# ClonoDynamics enables replicate-resolved inference of longitudinal T cell receptor repertoire dynamics

**DOI:** 10.64898/2026.09.24.754170

**Authors:** Ilenia Aversa, Raffaella Gallo, Antonio Abatino, Francesca Iannone, Lisa Isdraele, Caterina Giordano, Michael Marrano, Giuseppe Fiume, Camillo Palmieri, Giovanni Cuda

## Abstract

Longitudinal T cell receptor repertoire analysis can confound biological change with measurement noise and estimator-induced structure. We developed ClonoDynamics, a framework integrating latent abundance inference with replicate-decoupled estimates of directional change and cross-replicate displacement covariance. Pseudo-longitudinal series assembled from 12 independently processed technical measurements of one biological sample revealed apparent restoring dynamics caused by reusing measurements for conditioning and change estimation. Separating these roles between replicates largely removed this artifact. Synthetic controls preserving empirical detection patterns showed dose-dependent recovery of temporal accumulation, resolving the two larger imposed effects but not the smallest. Applied to TCRβ repertoires from ten healthy adults sampled weekly for six weeks, ClonoDynamics identified abundance-dependent directional change and shared fluctuations exceeding the technical reference. Within the analyzed abundance domain, these fluctuations showed no consistent positive accumulation over 1–5 weeks. ClonoDynamics provides an experimentally benchmarked framework for distinguishing estimator-induced patterns, replicate-consistent longitudinal variation, and temporal accumulation.

**One Sentence Summary:** ClonoDynamics combines technical replication and pseudo-longitudinal benchmarking to distinguish estimator-induced artifacts from replicate-consistent T cell receptor repertoire changes and assess their accumulation over time.

## INTRODUCTION

Longitudinal T-cell receptor (TCR) repertoire sequencing provides a way to track clonal expansion, contraction and persistence in health and after immune challenge^1–4^ . TCR clonotypes span several orders of magnitude in abundance, producing strongly skewed distributions in which relatively few abundant clones coexist with many rare ones^5–7^. This organization can retain signatures of repertoire formation and early clonal expansion^8^, while studies of healthy individuals reveal persistent clonotypes and relatively stable repertoire-level features over months to years^1–3^. Quantifying clonal fluctuations within this organization is therefore important for establishing a baseline against which immune perturbations can be assessed^2,3^.

T-cell survival and proliferation depend on self-peptide–MHC interactions, homeostatic cytokines and antigen exposure, with distinct requirements across naïve, memory and activated populations^9^. These differences can generate heterogeneous clonal growth and loss rates^9,10^, and models of fluctuating fitness provide mechanisms linking clonal dynamics to broad abundance distributions^5,8^. However, a clone-size distribution does not by itself establish how subsequent change depends on initial abundance or whether fluctuations accumulate over time.

A major obstacle is that sequencing measures only a noisy sample of the repertoire: finite sampling, library preparation and sequencing depth affect both abundance estimates and detection, particularly for rare clonotypes^11–14^. Multidimensional frameworks such as immuneREF characterize similarities between repertoires^15^, whereas replicate-calibrated probabilistic approaches, including NoisET^16^, model the relationship between latent frequencies and observed counts^11,16^. These approaches have been used to detect responding clones^4^ and infer stochastic repertoire dynamics in healthy individuals^3^. Complementary recapture models account for the dependence of repeated clonotype detection on initial abundance and sample richness^17^.

A separate problem arises when clonotypes are grouped by a noisy initial abundance that also enters the calculation of their subsequent change. This mathematical coupling can generate a negative association between initial abundance and displacement even without biological change^18^. Noise-aware abundance estimates do not, by themselves, validate a downstream abundance-conditioned estimator. Detection requirements can also change the clonotype population being analyzed^14,17^. An empirical reference without elapsed biological time can therefore complement observation models by testing which patterns are generated by the measurement and analysis procedures.

Here, we introduce ClonoDynamics, a replicate-resolved framework integrating latent abundance inference with experimentally benchmarked estimators of longitudinal change. Pseudo-longitudinal series from independently processed aliquots of one biological sample benchmark estimator behaviour, while synthetic controls preserving empirical detection patterns test sensitivity to temporal accumulation. Applied to weekly TCRβ repertoires from ten healthy adults with paired technical replicates, the framework separately quantifies abundance-dependent directional change, fluctuations shared between replicates and their lag dependence over 1–5 weeks.

## RESULTS

### Technical replicates characterize uncertainty and depth-dependent shifts in clonotype abundance estimates

We analysed TCRβ repertoires from 10 healthy adults sampled at six consecutive weekly time points, with paired technical replicates at each time point (Figure 1; Supplementary Table 1). Following exclusion of three technically unsuccessful subject–timepoint samples, 114 repertoires corresponding to 57 complete replicate pairs were available for descriptive characterization. Median analysed read depth was 496,140 reads per repertoire (range, 133,193–2,034,625), with a median of 110,648 observed clonotypes (range, 37,444–276,052; Supplementary Data 2).

**Figure 1.**
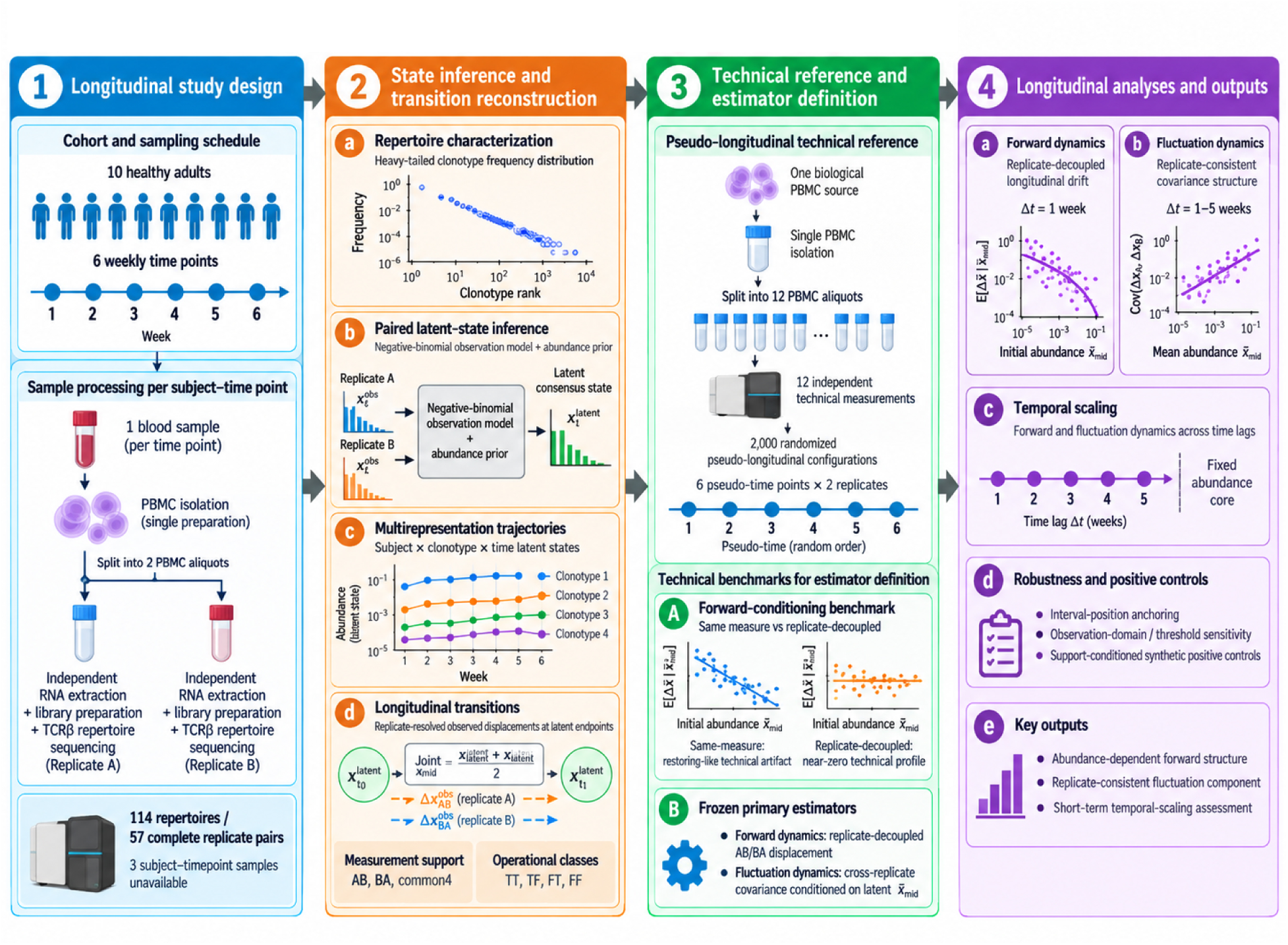
Study design, technical calibration, and longitudinal analysis framework of ClonoDynamics. (1) Longitudinal study design. Ten healthy adults were sampled weekly for six consecutive weeks. At each time point, PBMCs were isolated once and divided into two aliquots that were independently processed for RNA extraction, library preparation, and TCRβ repertoire sequencing. The study yielded 114 repertoires corresponding to 57 complete replicate pairs available for repertoire characterization. (2) State inference and transition construction. Repertoire architecture was characterized descriptively, and paired replicate counts were analysed with a pair-specific negative-binomial latent-state model. Latent consensus states, replicate-resolved observations, and observation annotations were retained through trajectory reconstruction and longitudinal transition assembly. Estimator-specific support comprised AB, BA, and common4, with TT, TF, FT, and FF retained as operational observation classes. (3) Technical reference and estimator definition. Twelve independently processed PBMC aliquots from a single biological source were reorganized into 2,000 pseudo-longitudinal configurations of six pseudo-time points and two replicates. Technical benchmarks identified same-measure coupling and defined the frozen primary estimators: replicate-decoupled AB/BA forward displacement and cross-replicate displacement covariance conditioned on latent midpoint abundance. (4) Longitudinal analyses and validation. Replicate-decoupled forward dynamics were evaluated at one week, whereas replicate-consistent fluctuations and their temporal scaling were analysed across lags of 1– 5 weeks. Support-conditioned synthetic positive controls tested recovery of prescribed accumulation, and robustness analyses evaluated interval position, common-start/common-end anchoring, operational observation domains, and observation thresholds (*α* = 0.10,0.05,0.025, and 0.01).

Clonotype frequencies were highly uneven, with many rare clonotypes and relatively few abundant ones (Figure 2A). Across six prespecified tail-threshold quantiles from 0.70 to 0.95, corrected Akaike Information Criterion (AICc) favoured the power-law model in 680 of 684 repertoire– threshold comparisons (99.4%), with similar results by Bayesian Information Criterion (BIC) (682/684, 99.7%; Figure 2B; Supplementary Data 2). In 678 comparisons, the lower-truncated log-normal optimum lay at its power-law boundary, indicating little support for additional tail curvature. The primary, Kolmogorov-Smirnov (KS)-selected power-law exponents were concordant between technical replicates (Spearman ρ = 0.888; median absolute difference, 0.030; 57 pairs; Figure 2C). These descriptive comparisons concern selected frequency tails and do not establish a power-law distribution across the entire repertoire^19^.

**Figure 2.**
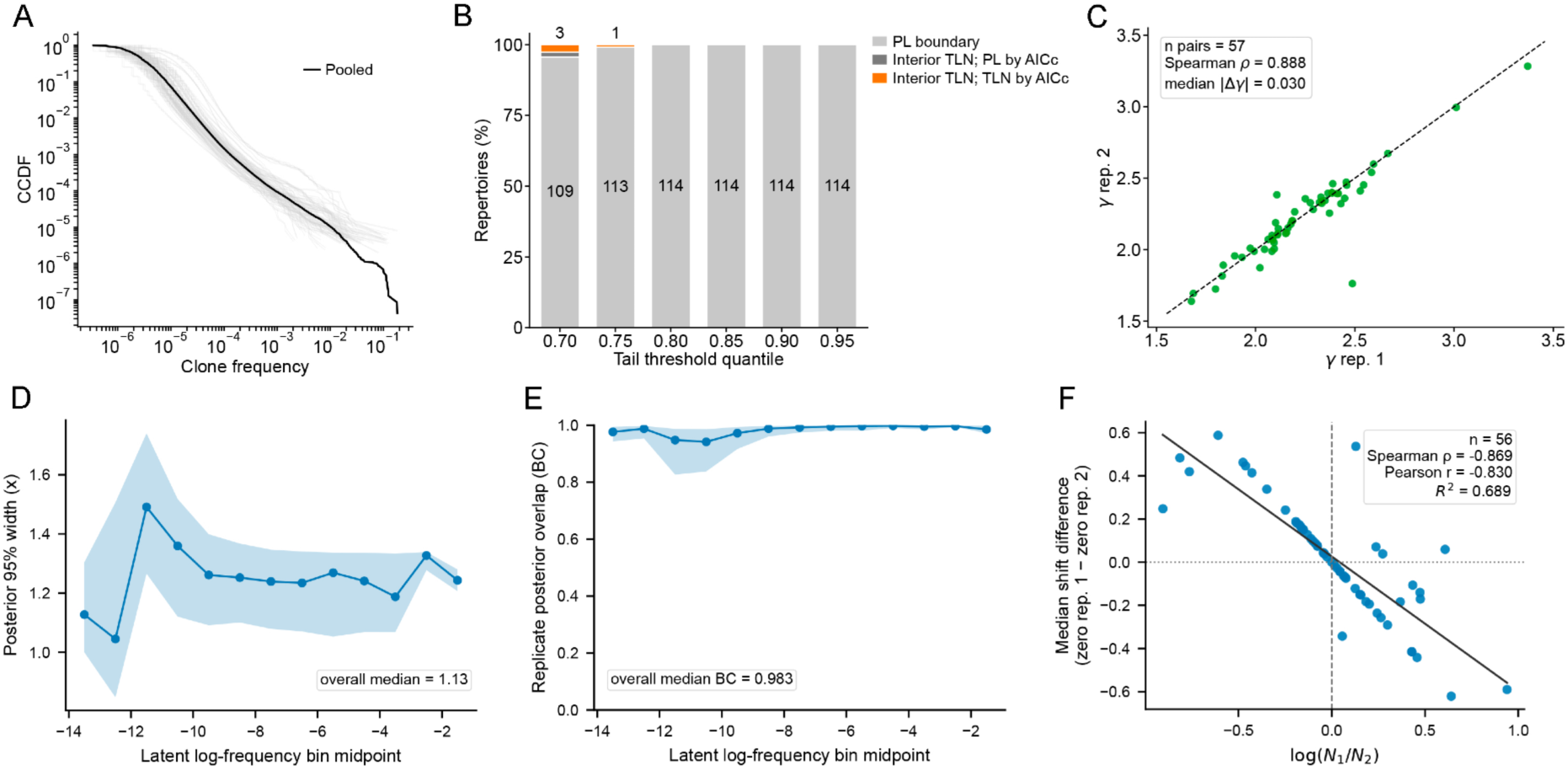
Repertoire architecture and diagnostics of replicate-resolved latent-state inference. (A) Complementary cumulative distribution functions (CCDFs) of clonotype relative frequencies across individual TCRβ repertoires (gray), with the pooled distribution shown in black. (B) Complexity-adjusted comparison of continuous power-law (PL) and jointly fitted lower-truncated log-normal (TLN) tail models across six prespecified empirical tail-threshold quantiles. Stacked bars distinguish repertoires in which the TLN optimum lies at the PL boundary, an interior TLN optimum is obtained but PL is retained by AICc, or an interior TLN is favored by AICc. Selected repertoire counts are shown within or above the bars. (C) Concordance of the fitted repertoire-level PL exponent, *γ*, between technical replicates. Each point represents one complete replicate pair, and the dashed line denotes identity. The inset reports the number of replicate pairs, Spearman correlation and median absolute inter-replicate difference. (D) Posterior 95% credible-interval width of the latent consensus as a function of inferred latent log-frequency. Points show abundance-bin medians and shading indicates the interquartile range. The inset reports the overall median posterior width. (E) Agreement between the two single-replicate latent posteriors, quantified by the exact discrete-posterior Bhattacharyya coefficient, as a function of inferred latent log-frequency. Points show abundance-bin medians and shading indicates the interquartile range. The inset reports the overall median Bhattacharyya coefficient. (F) Paired ONE_POSITIVE diagnostic relating sequencing-depth imbalance between technical replicates to the latent-versus-positive representation shift. Each point represents one subject–timepoint pair. The x-axis shows log(*N*_1_/*N*_2_), and the y-axis shows the difference between the median *x*_latent_ − *x*_positive_shift when replicate 1 versus replicate 2 is the zero-count replicate. The solid line shows the linear fit; dashed vertical and dotted horizontal lines denote zero. The inset reports the number of paired observations, Spearman and Pearson correlations, and the coefficient of determination. Panels D–F are descriptive diagnostics and do not impose abundance-, posterior-quality-, or detectability-based filtering. ONE_POSITIVE denotes clonotype states detected in exactly one of the two technical replicates and is not interpreted as biological presence or absence. Source data are provided in Supplementary Data 2 and 3.

We next used a pair-specific negative-binomial observation model to infer a shared underlying clonotype frequency from the two replicate counts and their sequencing depths^11,16^ . All clonotypes detected in either replicate were retained, including zero counts for paired non-detections. The paired posterior median log-frequency defined the latent consensus abundance; parameters from the descriptive tail fits were not transferred to this inference (Methods; Supplementary Figure 1; Supplementary Data 1). The model converged for 56 of 57 pairs. Exclusion of the non-converged fit for subject 5 at the second time point left 9,580,290 clonotype–timepoint states, with no further abundance or posterior-quality filtering (Supplementary Data 3).

Posterior uncertainty varied non-monotonically with estimated abundance (Figure 2D). The median 95% credible-interval width was 1.13 natural-log units (IQR, 0.94–1.47), corresponding to a median upper-to-lower frequency-limit ratio of 3.11 (Supplementary Figure 2A). The two single-replicate posterior distributions generally overlapped substantially, with a median Bhattacharyya coefficient of 0.983 (IQR, 0.939–0.996; Figure 2E; Supplementary Figure 2B). These quantities describe uncertainty and replicate agreement under the fitted model, not accuracy against an independent abundance reference.

Most states were detected in only one replicate (ONE_POSITIVE, 60.2%), whereas 39.8% were detected in both (TWO_POSITIVE). Relative to the corresponding summary based only on positive measurements, *x*_positive_, the latent consensus tended to be lower: median *x*_latent_ − *x*_positive_ was −0.375 for ONE_POSITIVE states and −0.205 for TWO_POSITIVE states (Supplementary Figure 2C; Supplementary Data 3).

Within ONE_POSITIVE states, this difference was associated with relative sequencing depth. Across 112 subject–timepoint–orientation summaries, greater depth in the zero-count replicate relative to the positive replicate was associated with a more negative median shift (Spearman ρ = −0.552; Supplementary Figure 2D). In the paired comparison, the difference between median shifts for the two zero-count orientations was negatively associated with log(*N*_1_/*N*_2_) (Spearman ρ = −0.869; 56 pairs; Figure 2F). These descriptive associations were calculated from grouped summaries across repeated visits from 10 participants, rather than treating clonotypes as independent depth measurements. Under the fitted model, the pattern is consistent with non-detection providing stronger evidence for low abundance in the more deeply sequenced replicate. Posterior-integrated state detectability closely tracked the factorized combination of replicate-specific predictive probabilities (Spearman ρ = 0.9999; median absolute difference, 0.00635; Supplementary Figure 2E). This internal diagnostic compares two treatments of shared frequency uncertainty rather than providing external calibration; additional diagnostic relationships are shown in Supplementary Figure 2F. The resulting state records preserved replicate-specific measurements and kept observed detection, predictive detectability and operational observability distinct, providing the inputs for longitudinal transition assembly.

### A common transition dataset defines estimator-specific analysis support

Within each participant, clonotype states were linked by amino-acid CDR3 sequence across all available sampling-time pairs. The resulting dataset contained 2,822,922 transitions from 928,016 unique subject–clonotype pairs, spanning all 15 nominal sampling-time pairs and temporal lags of 1–5 weeks. Following the upstream exclusions described above, 130 subject-specific intervals remained available. Their number decreased from 43 at a one-week lag to 9 at five weeks, whereas median transition counts per available interval remained broadly similar across lags, at approximately 21,000–23,700 (Supplementary Figure 3A,B; Supplementary Data 4).

Each transition retained replicate-specific counts, frequencies and sequencing depths at both endpoints, alongside latent abundance representations, uncertainty, predictive detectability and operational observation annotations. This structure preserved the measurements needed by different downstream estimators. Required variables were complete on their intended analysis support, and structural audits identified no mismatches (Supplementary Data 4).

Analysis eligibility was defined by the positive measurements required for conditioning and displacement estimation. The AB orientation used observed initial log-frequency in replicate 1 for conditioning and log-frequency change in replicate 2 for displacement, requiring replicate 1 to be positive initially and replicate 2 at both endpoints. BA reversed these roles. Across all lags, AB and BA were supported by 1,479,321 transitions (52.40%) and 1,480,279 transitions (52.44%), respectively. Cross-replicate fluctuation analysis required positive counts in both replicates at both endpoints, defining the common4 support of 1,182,069 transitions (41.87%). These requirements allowed observed displacements to be calculated without pseudo-counts. AB and BA eligibility fractions remained similar across lags, while common4 represented 39.0–42.3% of the assembled transitions (Supplementary Figure 3C; Supplementary Data 4).

Operational observability was annotated separately using the empirical endpoint observation statistic defined in Methods. At the reference threshold *α* = 0.05, endpoints were classified as T when *p* < *α* and F otherwise. Combining the endpoint labels yielded TT, TF, FT and FF transitions, accounting for 11.82%, 10.75%, 9.64% and 67.78% of the dataset, respectively, with broadly similar composition across lags (Supplementary Figure 3D; Supplementary Data 4). These labels described operational observation status, not biological persistence, emergence, extinction or absence, and did not restrict the primary AB, BA or common4 supports. We next benchmarked the corresponding estimators in the pseudo-longitudinal technical reference, where biological time was absent.

### Pseudo-longitudinal benchmarking reveals estimator-specific artifacts

To test estimator behaviour without elapsed biological time, we constructed a pseudo-longitudinal reference from 12 independently processed technical measurements of a single PBMC source. The same measurements were rearranged into 2,000 configurations of six pseudo-time points with two replicates each, using every measurement once per configuration. Pseudo-lags 1–5 reproduced the nominal longitudinal sampling geometry, not biological duration. Analyses were performed within configurations; reported ranges are the 2.5th–97.5th percentiles across configurations and describe randomization variability within this single-source reference, not biological confidence intervals (Supplementary Data 5).

At pseudo-lag 1, the primary replicate-decoupled estimator, combining AB and BA with equal weights, yielded a median abundance-dependence slope of −0.0019 (−0.0658 to 0.0630; Figure 3A). We then compared same-measure and replicate-decoupled estimates on identical *common4* transitions. Reusing the initial measurement for both conditioning and displacement estimation generated an apparent restoring pattern, with positive mean changes at low abundance and negative changes at high abundance. Its median slope was −0.4267 (−0.4856 to −0.3474), compared with −0.0016 (−0.0614 to 0.0549) after decoupling. The average absolute bin-specific mean change decreased by a median of 85.7% across configurations (Figure 3B; Supplementary Data 5). Thus, an apparent restoring pattern arose without biological time and was largely removed by separating the two measurement roles, consistent with mathematical coupling between initial value and change^18^.

**Figure 3.**
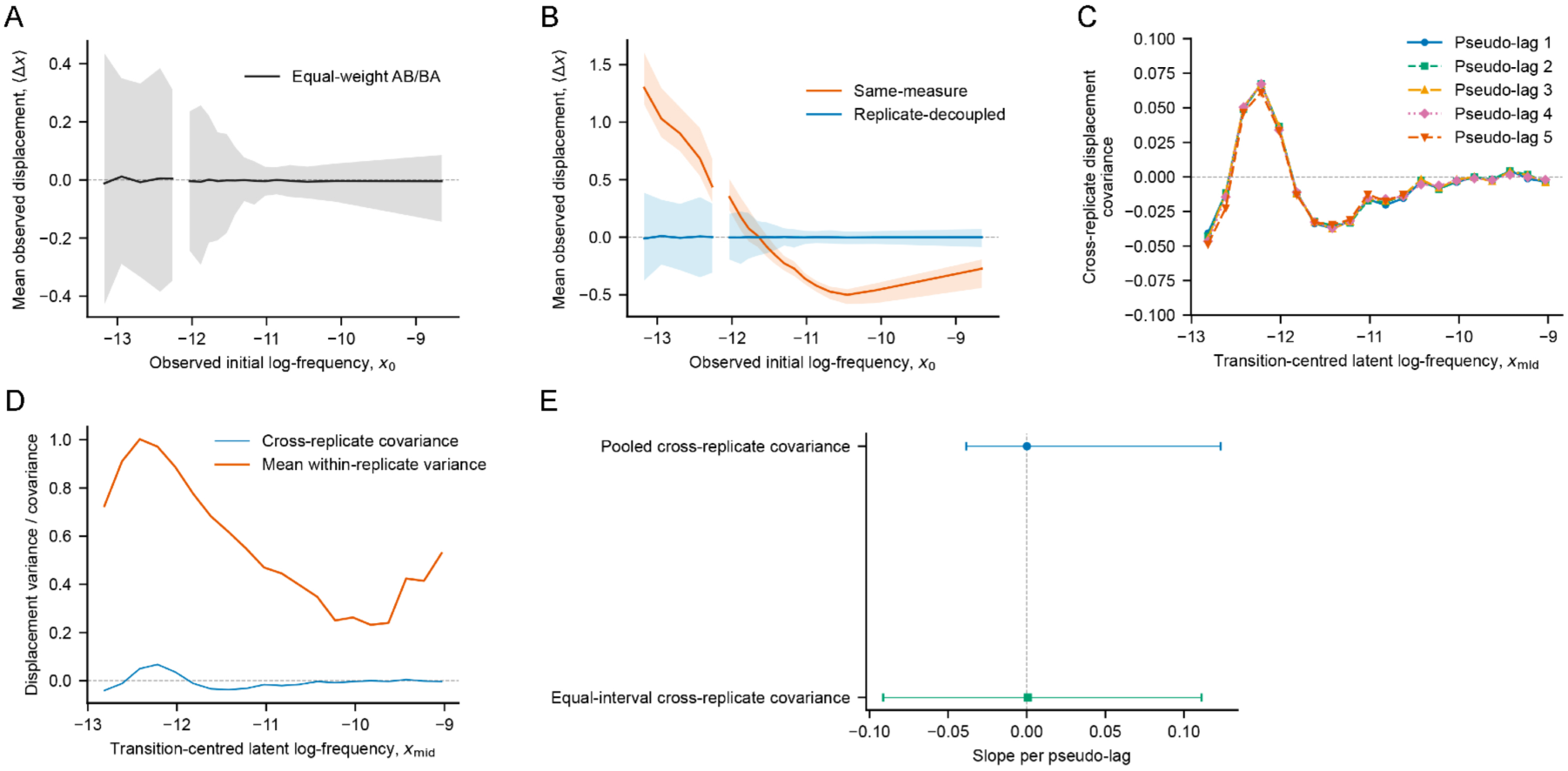
Pseudo-longitudinal technical-null benchmarks for forward dynamics and cross-replicate fluctuations. Lines and points show medians across 2,000 randomization configurations; shaded bands (A,B) and error bars (E) indicate the corresponding 2.5th–97.5th randomization percentiles. Reference lines mark zero. Panels B–E use *common4* transitions; bins not meeting the plotting-support criteria are omitted. (A) Mean observed log-frequency change at pseudo-lag 1 versus observed initial log-frequency, *x*_-_, estimated using the equal-weight AB/BA combination. (B) Matched same-measure (orange) and replicate-decoupled (blue) forward profiles at pseudo-lag 1, evaluated on identical transitions. (C) Cross-replicate displacement covariance, Cov(Δ, *x_A_*Δ*x_B_*), versus transition-centred latent log-frequency, *x_mid_*, for pseudo-lags 1–5. Colours and markers distinguish pseudo-lags. (D) Cross-replicate displacement covariance (blue) and mean within-replicate displacement variance (orange) across *x_mid_*bins at pseudo-lag 1. (E) Slopes of cross-replicate covariance versus pseudo-lag, calculated separately within each configuration using pooled transitions (top) or equally weighted pseudo-time intervals (bottom). Source data are provided in Supplementary Data 5.

A complementary diagnostic based on latent displacements also depended on the conditioning coordinate. Conditioning on initial latent abundance, *x*_0_, produced a negative median slope (−0.1907), whereas the transition-centred coordinates *x*_mid_ and *x*_∗_ yielded slopes near zero (Supplementary Figure 4A). Latent-state inference therefore did not, by itself, eliminate conditioning-dependent structure.

We next quantified the covariance between observed replicate-specific displacements on *common4*, conditioned on latent midpoint abundance, *x*_mid_. Binwise median profiles showed a small, signed abundance-dependent structure that was similar across pseudo-lags within supported bins (Figure 3C). At pseudo-lag 1, median covariance ranged from −0.041 to 0.067 across bins, compared with 0.232–1.002 for the mean within-replicate displacement variance (Figure 3D). The cross-replicate component was therefore much smaller than within-replicate variability but could not be assumed to vanish at every abundance.

Pooling *common4* transitions within each configuration and pseudo-lag, without abundance binning, yielded median covariances of 0.00006–0.00126 across lags (Supplementary Figure 4C). Randomization variability was greatest at pseudo-lag 5, which contained one pseudo-time interval per configuration and had a median *common4* fraction of 4.6%, compared with 9.9–11.5% at pseudo-lags 1–4 (Supplementary Figure 4B). These pooled summaries addressed aggregate technical behaviour and did not replace abundance-resolved profiles.

Temporal slopes calculated separately for all 2,000 configurations had medians close to zero: 0.00012 per pseudo-lag (−0.0385 to 0.1233) for pooled covariance and 0.00074 (−0.0913 to 0.1112) for equally weighted interval-specific covariances (Figure 3E; Supplementary Data 5). The technical reference therefore showed no systematic positive accumulation under these summaries. Together, the benchmarks supported replicate-decoupled observed estimators for forward analysis and an abundance-resolved technical reference for interpreting longitudinal cross-replicate covariance.

### Replicate-decoupled forward dynamics retain abundance dependence beyond the technical reference

We next applied the replicate-decoupled AB/BA estimator to the longitudinal cohort at a one-week lag. The reciprocal orientations produced concordant abundance-dependent profiles, with slopes of −0.241 for AB and −0.246 for BA and a Pearson correlation of 0.973 across 20 matched abundance bins (mean absolute AB–BA difference, 0.052; Figure 4A; Supplementary Figure 5C). Their equal-weight combination showed a negative relationship between observed initial log-frequency and subsequent mean displacement. A linear descriptor of this profile yielded a slope of −0.243 (95% subject-bootstrap CI, −0.290 to −0.213) and a fitted zero-crossing at *x*_0_ = −11.72 (−11.91 to −11.49). Subject-specific slopes were negative in all 10 participants (−0.389 to −0.184), and leave-one-subject-out estimates remained close to the full-cohort estimate (−0.252 to −0.233; Supplementary Figure 5A,B; Supplementary Data 6).

**Figure 4.**
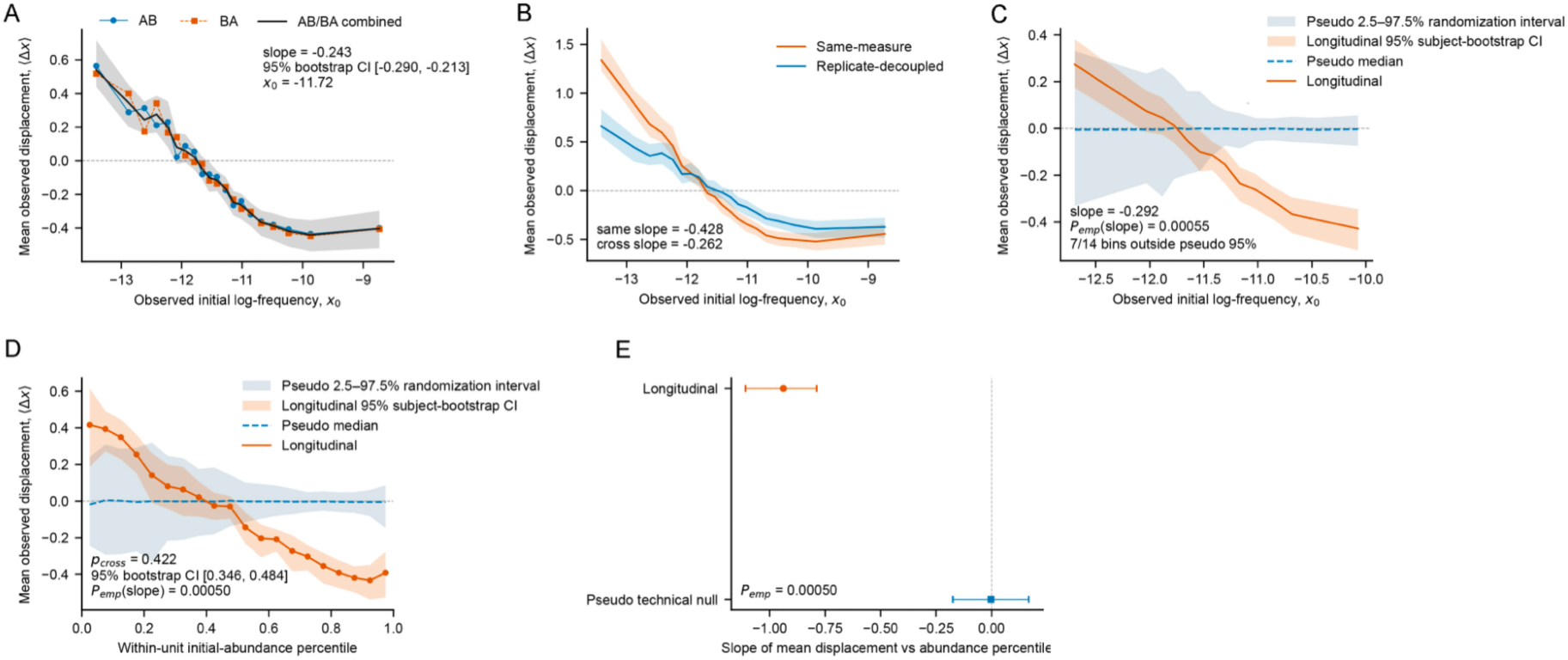
Longitudinal replicate-decoupled forward dynamics and comparison with the pseudo-longitudinal technical reference. (A) Mean observed log-frequency displacement as a function of initial observed log-frequency for the AB and BA orientations and their equal-weight combined profile. Shading indicates the 95% subject-bootstrap confidence interval of the combined estimate. The annotation reports the descriptive linear slope and fitted zero-crossing. (B) Matched same-measure and replicate-decoupled forward profiles evaluated on identical *common4* transitions. Shaded bands indicate 95% subject-bootstrap confidence intervals; annotations report the corresponding descriptive linear slopes. (C) Longitudinal and pseudo-longitudinal forward profiles over their shared absolute log-frequency support. Shading indicates the longitudinal 95% subject-bootstrap confidence interval and the pointwise pseudo 2.5th–97.5th randomization interval. The annotation reports the longitudinal slope, its lower-tail empirical probability under the pseudo reference (*P*_e3p_), and the number of shared bins falling below the pointwise pseudo interval. (D) Longitudinal and pseudo-longitudinal forward profiles after expressing initial abundance as a within-unit rank on a 0–1 scale. Ranks were defined within subject–orientation combinations for the longitudinal data and within configuration–orientation combinations for the pseudo reference. Shading indicates the longitudinal 95% subject-bootstrap confidence interval and the pointwise pseudo 2.5th–97.5th randomization interval. The annotation reports the fitted longitudinal zero-crossing rank and its 95% bootstrap interval, together with *P*_e3p_ for the longitudinal slope. (E) Longitudinal and pseudo-reference slopes of mean displacement versus within-unit abundance rank. Horizontal error bars indicate the longitudinal 95% subject-bootstrap confidence interval and the pseudo 2.5th–97.5th randomization interval, respectively. The vertical reference line denotes zero slope. Slopes and zero-crossings are descriptive summaries from linear fits to the corresponding binned profiles. *P*_e3p_ denotes the lower-tail empirical probability of the longitudinal slope under the pseudo-longitudinal reference. Source data are provided in Supplementary Data 6.

To isolate the effect of reusing the initial measurement, we compared same-measure and replicate-decoupled profiles on identical *common4* transitions. The same-measure profile had a more negative slope (−0.428; 95% bootstrap CI, −0.494 to −0.392) than the replicate-decoupled profile (−0.262; −0.309 to −0.230; Figure 4B). Direct measurement coupling therefore strengthened the abundance dependence, but the negative relationship remained after the conditioning and displacement measurements were separated.

We then compared longitudinal and pseudo-longitudinal profiles on shared observed-log-frequency support. Fourteen shared coordinates spanned −12.69 to −10.07, and 1,825 of the 2,000 pseudo configurations provided complete coverage of this grid. On this restricted domain, the longitudinal slope was −0.292, compared with a pseudo median of −0.0008 (2.5th–97.5th randomization percentiles, −0.088 to 0.091; lower-tail *P*_emp_ = 0.00055; Figure 4C). Seven of the 14 longitudinal estimates fell below the pointwise pseudo interval, all at the more abundant end of the shared range.

As a complementary comparison, we expressed initial abundance as a relative rank within each subject–orientation combination or pseudo configuration–orientation combination. This aligned relative positions within the respective abundance distributions rather than equating absolute frequencies. The longitudinal slope on the 0–1 rank scale was −0.938 (95% subject-bootstrap CI, −1.109 to −0.788), compared with a pseudo median of −0.0029 (2.5th–97.5th randomization percentiles, −0.175 to 0.168; lower-tail *P*_emp_ = 0.00050; Figure 4D,E). Thirteen of 20 bins lay outside the pointwise pseudo interval. The fitted zero-crossing was 0.422 on this rank scale (95% bootstrap CI, 0.346–0.484; Supplementary Figure 5D; Supplementary Data 6).

Together, these comparisons show that the one-week abundance-dependent profile persisted after replicate decoupling and differed from the single-source technical reference on both shared log-frequency and relative-rank scales. We next asked whether longitudinal changes also contained a fluctuation component shared between technical replicates.

### Cross-replicate covariance identifies replicate-consistent longitudinal fluctuations beyond the technical reference

Having identified a replicate-decoupled directional component, we next asked whether longitudinal clonotype changes also contained a component reproduced across the two technical replicates. We quantified this component as the covariance between replicate-specific observed log-frequency displacements, conditioned on transition-centred latent abundance, *x*_mid_, and restricted to *common4* transitions. The *common4* support contained 394,868, 307,424, 226,155, 173,355 and 80,267 transitions at lags of 1–5 weeks, respectively, corresponding to 39.0–42.3% of the complete transition universe (Supplementary Figure 6B; Supplementary Data 7).

Cross-replicate covariance was positive across all 20 abundance bins at every lag (Figure 5A). The 95% subject-bootstrap interval remained above zero in all bins at weeks 1–4 and in 19 of 20 bins at week 5. Median bin-wise covariance ranged from 0.476 to 0.569 across the five lags. At one week, median cross-replicate covariance was 0.516, compared with a median mean within-replicate displacement variance of 0.819 and a median replicate-specific excess of 0.217 (Figure 5B). The shared fraction, operationally defined as cross-replicate covariance divided by mean within-replicate variance, increased across the abundance axis and had a median of 0.735 at one week, with similar medians across lags (0.720–0.741; Supplementary Figure 6A). Thus, replicate-specific longitudinal displacements contained a substantial covariance component alongside additional replicate-specific variability.

**Figure 5.**
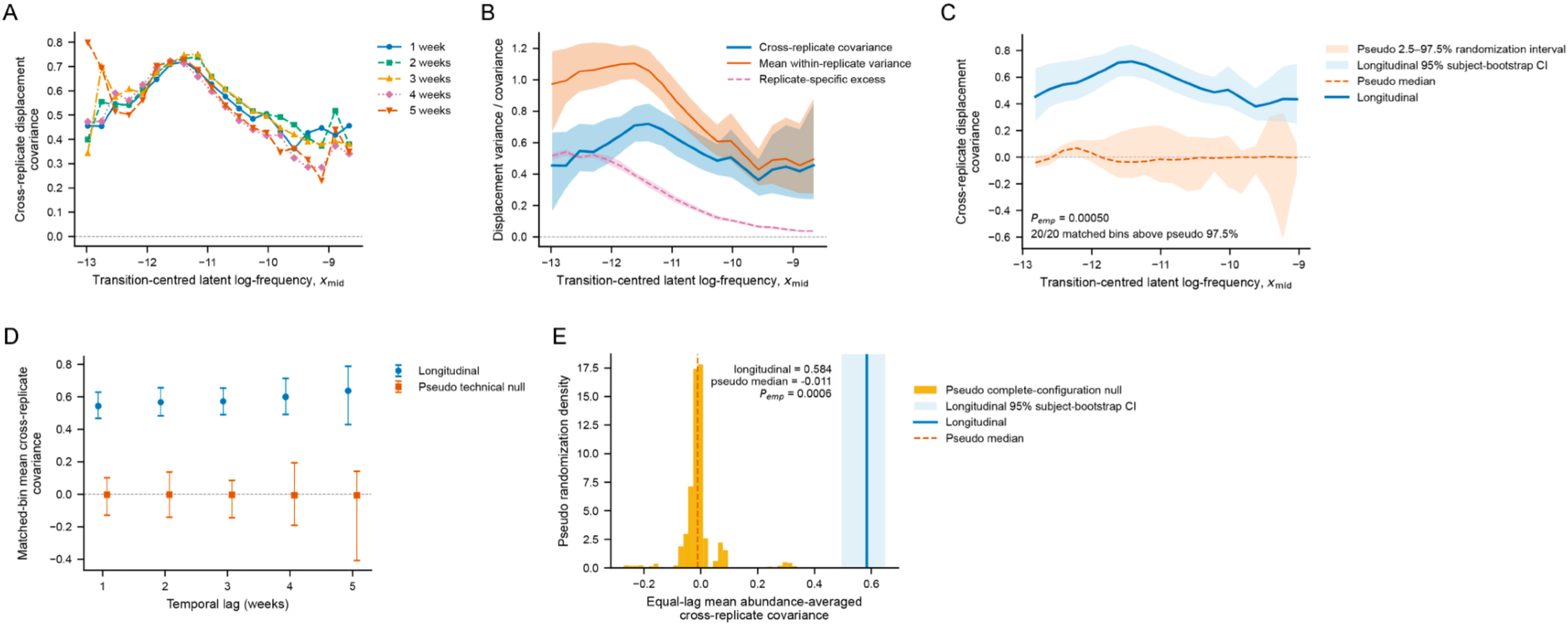
Longitudinal cross-replicate fluctuation covariance and comparison with the pseudo-longitudinal technical null. (A) Cross-replicate displacement covariance across transition-centred latent log-frequency, *x_mid_*, for temporal lags of 1–5 weeks. (B) Fluctuation decomposition at one week, showing cross-replicate covariance, mean within-replicate displacement variance and replicate-specific excess. Shaded regions indicate 95% subject-bootstrap confidence intervals. (C) Longitudinal and pseudo-longitudinal cross-replicate covariance at one week over matched absolute *x_mid_* support. Shaded regions indicate the longitudinal 95% subject-bootstrap confidence interval and the pseudo 2.5th–97.5th randomization interval. The annotation reports the empirical probability for the abundance-averaged covariance and the number of matched bins above the pseudo 97.5th percentile. (D) Abundance-averaged cross-replicate covariance over lag-specific matched support for temporal lags of 1–5 weeks. Error bars show longitudinal 95% subject-bootstrap confidence intervals and pseudo 2.5th–97.5th randomization intervals. (E) Equal-lag summary of the matched-support covariance. The histogram shows the pseudo complete-configuration null; the longitudinal estimate and its 95% subject-bootstrap confidence interval are shown in blue, and the pseudo median by the dashed line. The annotation reports the empirical probability of the longitudinal joint statistic under the pseudo null. Source data are provided in Supplementary Data 7.

We next compared this covariance with the pseudo-longitudinal technical reference on lag-specific matched absolute-abundance support. At one week, longitudinal covariance exceeded the pointwise pseudo 97.5th percentile in all 20 matched bins (Figure 5C). The equal-bin mean covariance over this support was 0.543 (95% subject-bootstrap CI, 0.468–0.628), compared with a pseudo median of −0.003 (2.5th–97.5th randomization percentiles, −0.129 to 0.101; upper-tail *P*_emp_ = 0.00050). Comparable separation was observed at weeks 2–4, with longitudinal estimates of 0.567, 0.572 and 0.600 and *P*_emp_ ≈ 5 × 10^−4^ at each lag (Figure 5D).

At week 5, the longitudinal estimate remained higher than the pseudo reference (0.637 versus −0.007), but the pseudo distribution was substantially broader (2.5th–97.5th percentiles, −0.408 to 0.141) and the empirical evidence was weaker (*P*_emp_ = 0.0234). Nine of 11 matched abundance bins exceeded the pointwise pseudo 97.5th percentile, compared with all matched bins at weeks 1–4 (Supplementary Figure 6C,D; Supplementary Data 7).

As a joint summary across lags, giving equal weight to the five lag-specific matched-support estimates yielded a longitudinal covariance of 0.584 (95% subject-bootstrap CI, 0.495–0.649). Across the 1,584 pseudo configurations eligible at all five lags, the corresponding null distribution had a median of −0.011 and 2.5th–97.5th percentiles of −0.110 to 0.092 (*P*_emp_ = 0.00063; Figure 5E). This joint statistic summarizes longitudinal–technical separation across lags; it is not a test of temporal scaling.

Together, these analyses identify a replicate-consistent longitudinal fluctuation component that exceeds the single-source pseudo-longitudinal technical reference. Because the matched abundance domain changes with lag, neither the lag-specific estimates nor their equal-lag summary quantify temporal accumulation. We therefore next evaluated temporal scaling using a fixed abundance domain and a common longitudinal subject set.

### Replicate-consistent fluctuations show no positive temporal accumulation over 1–5 weeks

We next asked whether the replicate-consistent fluctuation component increased with elapsed time. To compare the same abundance domain and the same participants at every lag, temporal scaling was evaluated within a fixed core of 17 contiguous *x*_mid_ bins spanning −12.41 to −8.55 and nine complete-case participants (Supplementary Figure 7A; Supplementary Data 8).

Within this fixed domain, core-averaged cross-replicate covariance was 0.563, 0.571, 0.537, 0.443 and 0.412 at lags of 1–5 weeks, respectively (Figure 6A). Covariance was therefore similar over the first three weeks and lower at weeks 4–5. The prespecified linear summary yielded a temporal slope of −0.043 per week (95% subject-bootstrap CI, −0.070 to −0.018), and none of the 2,000 subject-bootstrap slopes was positive (Supplementary Figure 7B; Supplementary Data 8).

**Figure 6.**
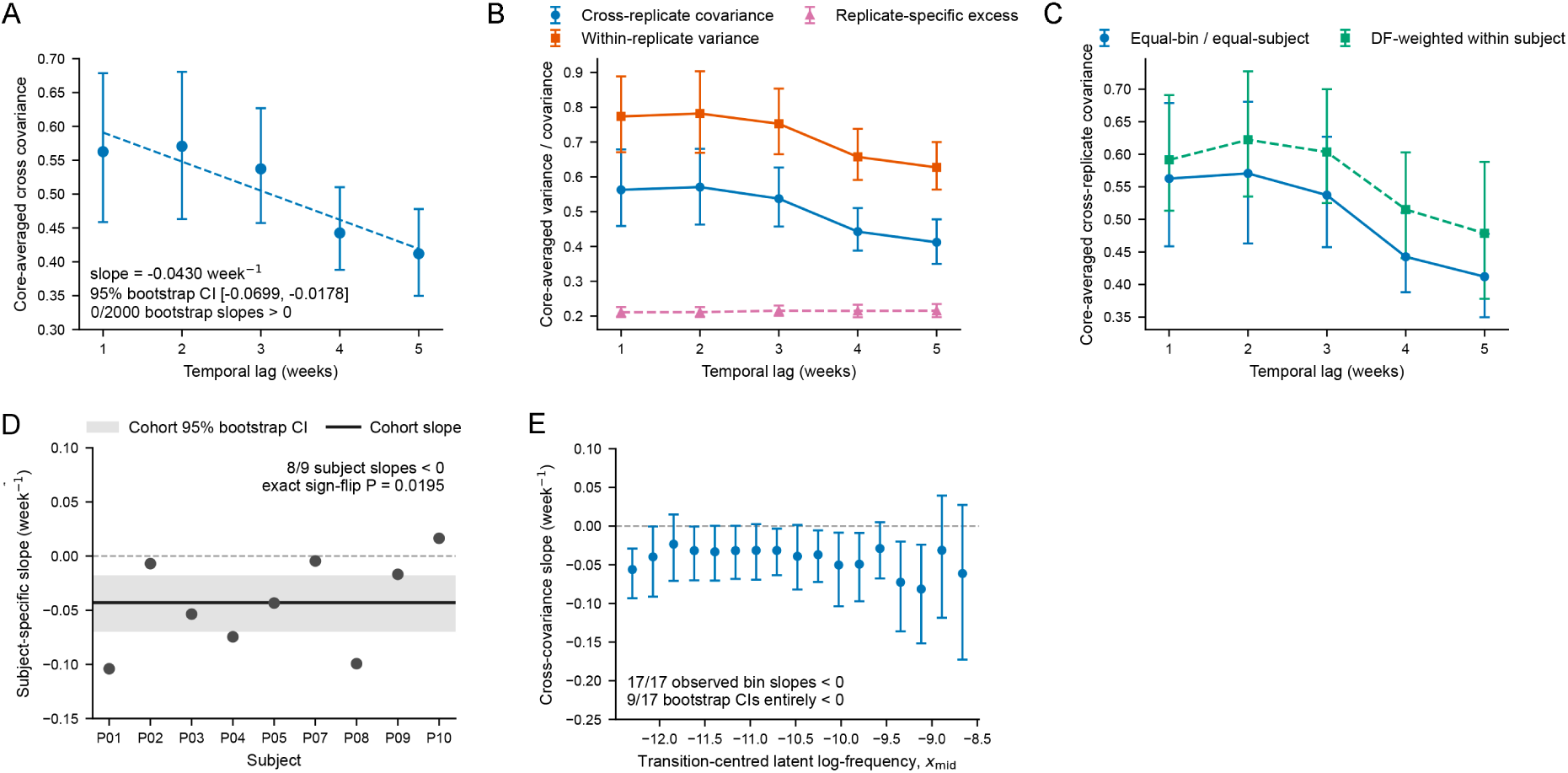
Temporal scaling of longitudinal cross-replicate fluctuation covariance. (A) Core-averaged cross-replicate covariance across temporal lags of 1–5 weeks, evaluated over the fixed 17-bin abundance core and the common set of nine complete-case participants. Error bars indicate 95% subject-bootstrap confidence intervals; the dashed line shows the descriptive linear temporal fit. The annotation reports the fitted slope, its 95% bootstrap interval, and the fraction of bootstrap slopes greater than zero. (B) Temporal profiles of cross-replicate covariance, mean within-replicate displacement variance, and replicate-specific excess over the same fixed abundance core. Error bars indicate 95% subject-bootstrap confidence intervals. (C) Sensitivity to abundance-bin weighting, comparing the primary equal-bin/equal-subject estimator with an alternative estimator in which abundance bins are weighted within subject by their covariance degrees of freedom. Error bars indicate 95% subject-bootstrap confidence intervals. (D) Subject-specific temporal slopes of cross-replicate covariance across the fixed abundance core. The horizontal line and shaded band indicate the cohort slope and its 95% subject-bootstrap confidence interval, respectively. The annotation reports the number of participants with negative slopes and the exact two-sided subject-level sign-flip test. (E) Temporal slopes estimated separately within each of the 17 abundance bins of the fixed core. Error bars indicate 95% subject-bootstrap confidence intervals. The annotation reports the number of bins with negative point estimates and the number whose bootstrap intervals lie entirely below zero. Horizontal reference lines in D and E denote zero slope. Source data are provided in Supplementary Data 8.

Mean within-replicate displacement variance showed a similar pattern, decreasing from 0.774 at one week to 0.627 at five weeks, with a slope of −0.0418 per week (95% bootstrap CI, −0.0710 to −0.0157). By contrast, replicate-specific excess remained approximately constant at 0.21 across the five lags, with a slope close to zero (0.0012 per week; 95% bootstrap CI, −0.0039 to 0.0068; Figure 6B). Thus, the decrease in within-replicate variance closely tracked that in cross-replicate covariance, while their difference remained approximately constant.

The negative temporal trend was retained under an alternative degrees-of-freedom weighting of abundance bins. This weighting increased the absolute covariance estimates but yielded a slope of −0.0333 per week (95% bootstrap CI, −0.0563 to −0.0122; Figure 6C; Supplementary Data 8). At the participant level, eight of nine slopes were negative, ranging from −0.104 to 0.016 per week; an exact two-sided subject-level sign-flip test gave *P* = 0.0195 (Figure 6D).

The signed trend was also distributed across the abundance core rather than confined to a single region. Point estimates were negative in all 17 bins, ranging from approximately −0.081 to −0.023 per week, with 95% subject-bootstrap intervals entirely below zero in 9 of 17 bins (Figure 6E; Supplementary Data 8).

A descriptive three-model comparison based on the five cohort-level lag estimates assigned the lowest AICc to the constant model, with ΔAICc = 0, 9.84 and 20.0 for the constant, signed-linear and positive-incremental models, respectively (Supplementary Figure 7C). We therefore interpret these data primarily as evidence against positive accumulation of replicate-consistent fluctuations over the observed 1–5-week interval. The lower covariance at weeks 4–5 supports a modest negative signed trend, but the five-lag profile does not establish a general decay law. Subject-level trajectories remained heterogeneous around the cohort pattern (Supplementary Figure 7D).

We next tested whether the analysis could recover known positive temporal accumulation using support-conditioned synthetic controls of increasing strength.

### Support-conditioned synthetic controls show dose-dependent recovery of imposed temporal accumulation

We next tested whether the absence of positive temporal accumulation in the longitudinal cohort could reflect limited analytical sensitivity. Four support-conditioned semi-synthetic scenarios preserved the analysis-eligible empirical sampling design, sequencing depths and replicate-specific detection patterns while introducing progressively stronger cumulative frequency variation. Full-frequency simulated values—the *oracle*—targeted increases of 0%, 25%, 50% and 100% in log-frequency displacement variance between one and five weeks. Positive doses were calibrated from the oracle before count generation, without using recovered estimates; the non-accumulating control retained visit-specific fluctuations but no cumulative component. All scenarios were analysed using the same fixed 17-bin abundance core and nine complete-case participants as the longitudinal analysis (Methods; Extended Methods; Supplementary Data 9).

Recovered temporal slopes increased with the prescribed accumulation dose (Figure 7A; Supplementary Figure 8). Slopes were −0.00446, −0.00086, 0.00412 and 0.01370 per week for the control and the 25%, 50% and 100% conditions, respectively. The 50% and 100% conditions had 95% subject-bootstrap intervals entirely above zero (0.00033–0.00759 and 0.00564–0.01986), whereas the intervals for the control and 25% condition included zero (Supplementary Data 9).

**Figure 7.**
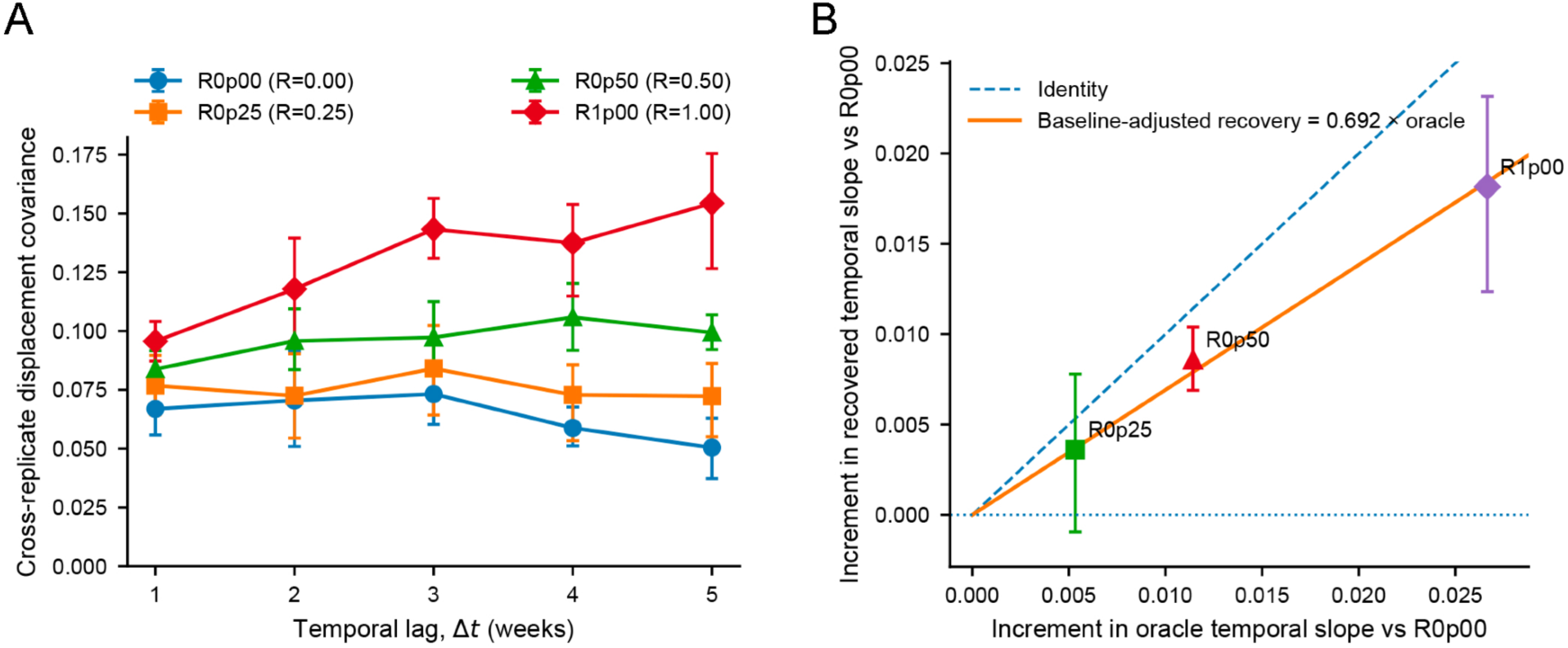
Support-conditioned synthetic positive controls for temporal accumulation. (A) Cross-replicate displacement covariance across temporal lags of 1–5 weeks for the four synthetic accumulation scenarios, corresponding to target oracle increases of 0%, 25%, 50% and 100%. Points show the recovered cohort estimates and vertical error bars the corresponding 95% subject-bootstrap intervals. (B) Baseline-adjusted recovery of the temporal slope for the three positive-dose scenarios relative to the non-accumulating R0p00 control. The horizontal axis shows the increment in oracle temporal slope and the vertical axis the corresponding increment recovered by the full analysis pipeline. Error bars indicate paired 95% subject-bootstrap intervals. The dashed line denotes identity, and the solid line shows the descriptive through-origin recovery fit. Source data are provided in Supplementary Data 9.

Because the control yielded a slightly negative recovered slope, we additionally quantified positive-dose increments relative to this common baseline. Recovered increments were 0.00360, 0.00858 and 0.01816 per week for the 25%, 50% and 100% conditions, corresponding to 67.6%, 75.1% and 68.1% of their respective oracle increments. Paired subject-bootstrap intervals excluded zero for the 50% and 100% increments but not for the 25% condition. A descriptive through-origin fit of recovered versus oracle increments yielded a recovery factor of 0.692 (Figure 7B; Supplementary Data 9). These recovery fractions are descriptive because oracle slopes are defined over the full simulated repertoire whereas recovered slopes are estimated on the selected observed-transition support; they are not correction factors applied to the empirical estimates.

These controls therefore show dose-dependent but attenuated recovery of imposed temporal accumulation under the tested observation geometry. The two larger prescribed effects were resolved, whereas the smallest remained unresolved. The results demonstrate that the analysis can recover positive temporal scaling of sufficient magnitude under these conditions, while not excluding weaker accumulation in the longitudinal cohort. This validation is conditional on the empirical detection structure of the available realization, and subject resampling does not constitute a power analysis across independent simulation realizations. We next examined whether the longitudinal temporal pattern was sensitive to sampling-interval composition.

### Interval-position and observation-domain controls qualify the interpretation of longitudinal dynamics

We next examined whether the longitudinal patterns were sensitive to sampling-interval composition or to the operational criteria used to define the observation domain. For cross-replicate covariance, neither cohort-level nor abundance-resolved interval-position tests showed BH-adjusted evidence of calendar-position structure at the evaluable lags of 1–3 weeks (Supplementary Figure 9A,B; Supplementary Data 10). Within-replicate variance showed a cohort-level interval-position signal at three weeks (*q* = 0.033), whereas replicate-specific excess did not. Restricting the temporal analysis to intervals sharing a common start or common end attenuated the negative cross-covariance slope. After requiring the same participants at every lag, both matched controls retained five participants: the common-start slope was −0.0095 per week (95% subject-bootstrap CI, −0.0335 to 0.0141) and the common-end slope was −0.0310 (−0.0965 to 0.0124), compared with −0.0430 in the primary analysis (Figure 8A; Supplementary Figure 9C,D). These controls therefore caution against interpreting the pooled negative trend as an intrinsic decay law, while the smaller anchored subject sets also limit precision.

**Figure 8.**
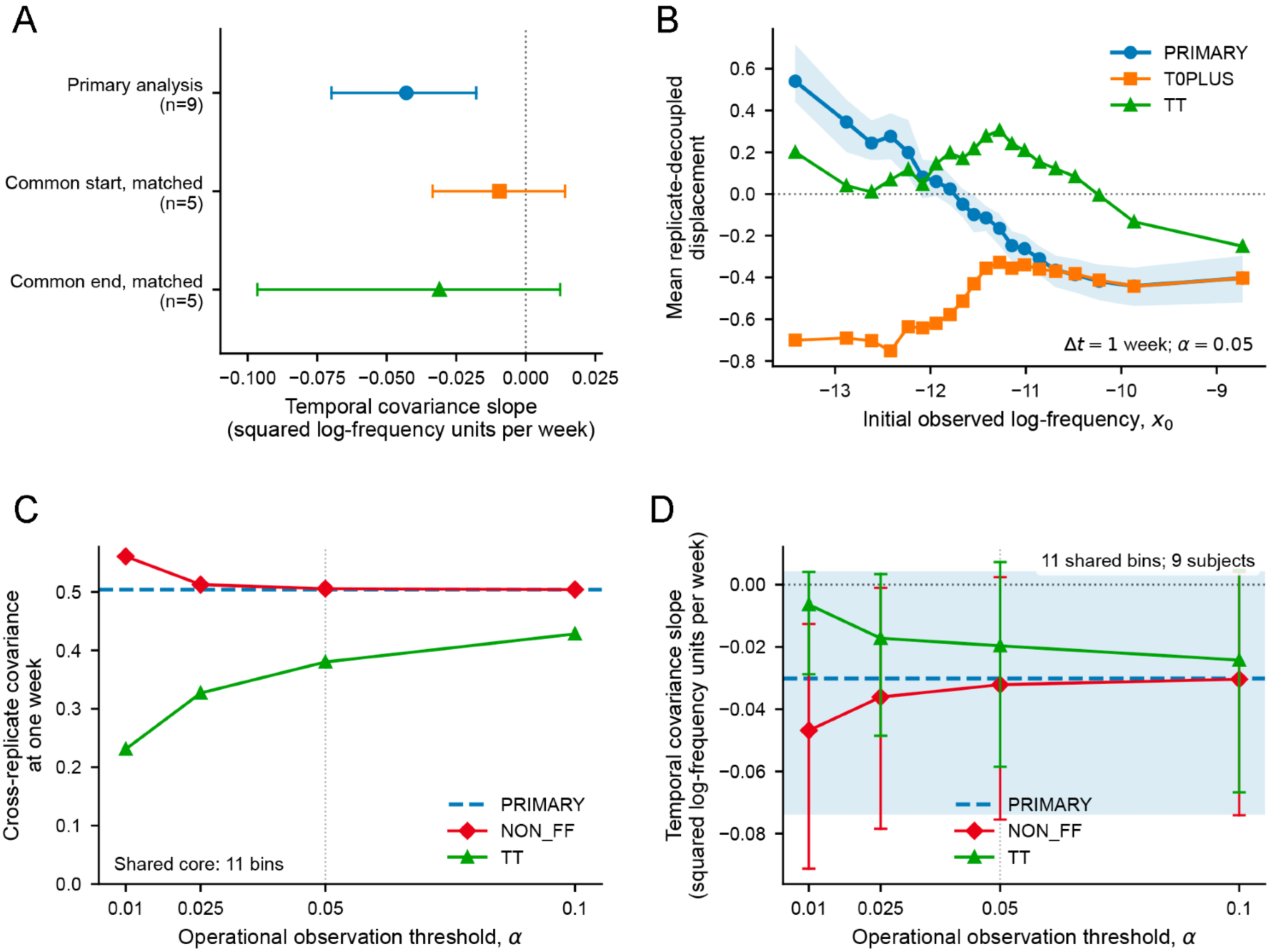
Interval-composition and operational observation-domain sensitivity analyses. (A) Temporal slopes of cross-replicate covariance for the primary Step 13 analysis and for matched common-start and common-end interval controls. Points indicate observed slopes and horizontal error bars the corresponding 95% subject-bootstrap intervals; the number of subjects contributing to each estimate is shown in parentheses. The vertical dotted line marks zero slope. (B) Replicate-decoupled forward displacement at a one-week lag for the PRIMARY, T0PLUS and TT operational domains at the reference threshold α=0.05. The shaded region denotes the 95% subject-bootstrap interval for the PRIMARY profile. The horizontal dotted line marks zero displacement. (C) Cross-replicate covariance at a one-week lag across operational observation thresholds for PRIMARY, NON_FF and TT, evaluated on the 11-bin cross-threshold common core. Points show the corresponding common-core estimates. The vertical dotted line marks the reference threshold α=0.05. (D) Temporal slopes of cross-replicate covariance across operational observation thresholds for PRIMARY, NON_FF and TT, evaluated on the same 11-bin common core and nine-subject cohort. Points indicate observed slopes and vertical error bars the corresponding 95% subject-bootstrap intervals. The vertical dotted line marks α=0.05 and the horizontal dotted line marks zero slope. Source data are provided in Supplementary Data 10 and 11.

We next examined sensitivity to the operational observation domain at the reference threshold *α* = 0.05. The PRIMARY forward analysis used replicate-based eligibility without operational filtering and had a negative abundance-dependence slope of −0.243. Restricting transitions to those operationally observable at the initial endpoint (T0PLUS = TT + TF) reversed the overall abundance dependence (slope, 0.101), whereas restricting to transitions observable at both endpoints (TT) substantially attenuated the negative slope (−0.058; Figure 8B; Supplementary Data 11). TF transitions showed a more negative profile (slope, −0.163) and contributed strongly negative displacements across lower-to-intermediate abundance regions (Supplementary Figure 10). Operational selection therefore changed the transition population being summarized rather than simply removing imprecise measurements. T/F labels denote operational observability and not biological presence, emergence, persistence or extinction.

We then repeated the operational-domain analyses across *α* = 0.10, 0.05, 0.025 and 0.01 without refitting latent abundances. Comparisons were restricted to common abundance support across thresholds. Increasing stringency progressively reduced the TT population and altered the restricted forward profiles (Supplementary Figure 11A–E). On the common 11-bin covariance core, one-week TT covariance decreased from 0.428 at *α* = 0.10 to 0.231 at *α* = 0.01, whereas NON_FF covariance increased from 0.504 to 0.561 over the same threshold range; PRIMARY was unchanged by construction (Figure 8C; Supplementary Figure 11F,G). Thus, the effect of operational selection on covariance magnitude depended on both the selected domain and the observation threshold.

Finally, temporal slopes were compared across domains and thresholds using the same nine participants and the 11-bin abundance core shared by all cross-threshold analyses. For this sensitivity analysis, covariance was pooled within each bin and then averaged equally across the common abundance core, as specified in Methods. PRIMARY was invariant to threshold by construction (slope, −0.0302 per week). NON_FF point estimates remained negative and became more negative with increasing stringency (−0.0304 to −0.0468), whereas TT slopes moved towards zero as the threshold became more stringent (−0.0242 to −0.0064); all TT bootstrap intervals included zero (Figure 8D; Supplementary Data 11). No consistently positive covariance–lag relationship emerged under any operational domain or threshold.

Together, these controls show that both interval composition and observation-domain selection affect the magnitude and abundance dependence of longitudinal estimates. They therefore constrain how the negative temporal trend and operationally restricted profiles should be interpreted. Across the tested domains and thresholds, however, we found no consistent evidence that the replicate-consistent fluctuation component increased as the interval between samples extended from one to five weeks.

## Discussion

ClonoDynamics addresses a central problem in longitudinal repertoire analysis: accounting for measurement noise does not automatically prevent artifacts introduced by conditioning or changes in analysis support. By integrating technical replication, pseudo-longitudinal benchmarking and estimator-specific support definitions, the framework distinguishes estimator-induced structure, variation reproduced across measurements and its dependence on elapsed time. In this healthy-adult cohort, replicate-decoupled forward profiles remained abundance dependent and cross-replicate displacement covariance exceeded the technical reference. Within the abundance domain supporting temporal analysis, however, this covariance showed no consistent positive accumulation over 1–5 weeks.

This contribution complements existing probabilistic approaches. Puelma Touzel and colleagues and NoisET established how replicate measurements can inform observation models and detection of clonal responses^11,16^, while subsequent studies inferred stochastic T-cell dynamics and extended related methods to memory B cells and plasmablasts^3,20^. ClonoDynamics instead benchmarks finite-time directional and fluctuation estimators without imposing a dynamical evolution model. Its primary displacements remain observed, whereas latent abundance coordinates are used for fluctuation conditioning. The methodological advance therefore lies in combining experimental estimator calibration with replicate decoupling and support-controlled comparisons, rather than introducing replicate-based noise modelling itself. The resulting estimates remain conditional on observation-model assumptions and measurement eligibility.

The pseudo-longitudinal experiment demonstrated the importance of estimator construction. Reusing a noisy initial measurement for both conditioning and displacement estimation generated a restoring-like profile, consistent with the established coupling between baseline and change^18^. Separating these roles largely removed this structure in the technical reference, but not in the longitudinal cohort. The remaining abundance dependence was concordant between reciprocal orientations and across participants, supporting a directional pattern not explained by the same-measure artifact alone under the conditions examined. This does not identify a particular homeostatic mechanism. Broad clone-size distributions can arise through fluctuating fitness and early-life expansion histories^5,8^; neither their shape nor a negative forward profile uniquely identifies feedback regulation. Competition, contraction after stimulation and subset redistribution remain possible contributors. The fitted slope and zero-crossing are finite-time descriptors, not an equilibrium size shared by individual clones.

Cross-replicate covariance provides complementary information. It was positive across the analysed abundance range and exceeded the pseudo-reference on matched support. Importantly, the technical reference itself exhibited small, abundance-dependent covariance, showing why a universal zero baseline would be inadequate. Nevertheless, covariance is not an unbiased partition of cellular and technical variance: positivity requirements, inferred conditioning coordinates and influences shared before replicate separation can affect it. The appropriate interpretation is therefore replicate-consistent longitudinal variation exceeding the technical conditions represented by the reference, rather than direct recovery of variance in cellular clone size.

The temporal analysis addressed a different question from longitudinal–technical separation. Fixing 17 abundance bins and nine participants limited changes in abundance range and subject composition across lags, although contributing clonotypes and intervals were not identical. Covariance was lower at four and five weeks, but common-start and common-end controls attenuated the negative slope and involved smaller participant sets. These results support the absence of consistent positive covariance–lag scaling in the tested window, not exact lag invariance, statistical equivalence to zero accumulation or a general decay law. The five-point model comparison likewise remains descriptive rather than establishing stationarity. This interpretation is compatible with longitudinal studies documenting repertoire persistence alongside ongoing clonal change^2,3^. In particular, geometric Brownian motion was motivated as an effective description over timescales longer than individual stimulation events^3^; a weekly, support-conditioned covariance profile is not directly interchangeable with its inferred diffusion coefficient or clonal lifetime.

The synthetic controls demonstrated dose-dependent recovery of imposed temporal accumulation, with positive slopes resolved for the two larger effects. Recovery was attenuated and the smallest dose remained unresolved. Because the generator preserved empirical detection masks and the scenarios shared random fields, these experiments establish conditional recoverability rather than a general detection limit. Subject bootstraps quantify uncertainty within the generated data, not power across independent simulation realizations. Weaker accumulation and changes outside the inherited observation support therefore remain possible.

Observation-domain sensitivity further clarifies what is being estimated. Applying operational T/F labels changed forward profiles and covariance amplitudes, demonstrating that selection changes the transition population rather than simply improving measurement precision. These labels are distinct from read positivity and do not indicate biological presence or extinction. The primary analyses also retain necessary measurement restrictions: AB/BA require their respective positive observations, and *common4* requires positivity in both replicates at both endpoints. This preferentially represents readily resampled clonotypes, not an intrinsically stable population or the entire rare-clonotype tail. Conclusions about temporal scaling consequently remain conditional on the analysed support. Previous studies similarly show that apparent persistence and recapture depend on sampling and experimental protocol^2,13,17^.

The main limitations concern generalizability and biological resolution. The study included ten healthy adults and a single-source technical reference, and its temporal conclusions apply to the analysed abundance domain over 1–5 weeks. RNA-based TCRβ sequencing measures relative expressed repertoire abundance rather than absolute cell numbers, while bulk blood profiling does not resolve the contribution of individual T-cell subsets. The framework therefore identifies reproducible longitudinal structure without assigning it to a unique cellular mechanism. Larger, phenotypically resolved cohorts and complementary cellular measurements will help establish how these patterns extend across populations, immune states and experimental protocols.

ClonoDynamics provides an experimentally benchmarked framework for distinguishing estimator-induced patterns from replicate-consistent repertoire variation and testing its dependence on elapsed time. In this healthy-adult cohort, abundance-dependent directional changes and substantial cross-replicate fluctuations exceeded the technical reference, yet fluctuation magnitude showed no consistent positive accumulation over 1–5 weeks. These findings demonstrate that reproducible short-term variation and temporal accumulation are distinct features of repertoire dynamics. By integrating technical replication, empirical benchmarking and support-controlled analysis, ClonoDynamics provides a reproducible basis for investigating how infection, vaccination and other immune perturbations alter these features.

## RESOURCE AVAILABILITY

### Lead contact

Further information and requests for resources, data, or code should be directed to and will be fulfilled by the lead contact, Prof. Camillo Palmieri.

### Materials availability

This study did not generate new unique reagents or renewable biological materials for distribution. Biological samples were collected and processed as described in the Methods and were used for T-cell receptor β-chain repertoire profiling.

### Data and code availability

Raw TCRβ sequencing data generated in this study have been deposited in the NCBI Sequence Read Archive under BioProject PRJNA1457834 and SRA Study SRP694925. Individual run accessions are reported in Supplementary Data 12.

The exact processed clonotype-level repertoire tables used as input to ClonoDynamics, together with the derived datasets, pseudo-longitudinal configuration manifests, analysis provenance, synthetic-validation summaries, integrity checksums, and final Supplementary Data workbooks required to reproduce the analyses reported in this study, are deposited in Zenodo under DOI 10.5281/zenodo.22925114.

The ClonoDynamics source code, including the analysis orchestration, scientific modules, robustness analyses, synthetic positive-control workflow, and publication-figure generation scripts, is available at https://github.com/ClonoDynamics/clonodynamics-pipeline. The frozen software release corresponding to the version used for this study is archived in Zenodo under DOI 10.5281/zenodo.22925873.

Any additional information required to reproduce or reanalyze the results reported in this paper is available from the lead contact upon reasonable request.

## Acknowledgments

We thank the staff of the Laboratory of Clinical Biochemistry and Clinical Molecular Biology, AOU “Renato Dulbecco” Hospital (Catanzaro, Italy), for their skilled technical assistance. We are also grateful to all study participants for their contribution to this work. This work was funded by the Italian Ministry of Health grant RF-2019-12370255.

## Author contributions

Conceptualization: C.P. and G.C.

Methodology: A.A., M.M., and C.G.

Investigation: I.A., R.G., A.A., L.I., F.I., M.M.

Software: A.A., R.G., P.C.

Formal analysis: C.P., A.A.

Data curation: I.A., R.G.

Funding acquisition: C.P. and G.C.

Supervision: G.F., C.P. and G.C.

Writing – original draft: I.A., R.G., L.I. and A.A.

Writing – review & editing: G.F., C.P., and G.C.

## Declaration of generative AI and AI-assisted technologies

During the preparation of this work, C.P. used ChatGPT (OpenAI) to support manuscript language editing, technical documentation, coding assistance, code review, debugging, and graphical rendering. All scientific questions, analytical definitions, estimands, parameter choices, workflow architecture, data interpretation, and final methodological decisions were defined and validated by the authors. AI-assisted code suggestions were inspected, tested, and revised by the authors before inclusion in the production pipeline, and all reported analyses were generated using the finalized, version-controlled ClonoDynamics codebase.

Figure 1 and Supplementary Figure 1 were generated using the image-generation functionality of ChatGPT from an author-specified description of the computational architecture. The resulting figures were subsequently checked against the executable ClonoDynamics workflow and iteratively corrected by the authors for scientific and topological accuracy. The authors take full responsibility for the content of the manuscript, software, analyses, and figures.

## EXPERIMENTAL MODEL AND SUBJECT DETAILS

### Study cohort and longitudinal sampling

Ten healthy adult volunteers aged 26–57 years (5 females and 5 males; Supplementary Table 1) were enrolled within a Biological Variation Data Critical Appraisal Checklist (BIVAC)-compliant framework. Regular clinical and biochemical monitoring throughout follow-up confirmed the absence of relevant inflammatory or infectious events during the study period^21,22^.

Peripheral blood was collected weekly for six consecutive weeks. At each time point, paired technical replicates were generated from the same blood draw. Following the technical failure of three subject–timepoint samples, the final dataset comprised 57 evaluable subject–timepoint samples and 114 TCRβ repertoires, organized as 57 complete technical-replicate pairs.

The study was approved by the Regional Ethical Review Board of Regione Calabria (protocol no. 69/2023). All participants provided written informed consent before enrolment.

## METHOD DETAILS

### PBMC isolation, TCRβ library preparation and sequencing

Peripheral blood mononuclear cells (PBMCs) were isolated by density-gradient centrifugation using Ficoll-Paque (Merck KGaA). At each longitudinal visit, the isolated PBMC suspension was divided into two aliquots, which were processed independently through RNA extraction and TCRβ libraries were prepared using the Oncomine™ TCR Beta-LR Assay (Thermo Fisher Scientific). Emulsion PCR, template enrichment, and Ion 530™ chip loading were performed using the Ion Chef™ System, followed by sequencing on an Ion GeneStudio™ S5 System.

The pseudo-longitudinal reference was generated by dividing a single PBMC preparation into 12 aliquots, each processed independently through the same workflow.

### TCRβ repertoire processing and clonotype assembly

Sample-specific BAM files generated by the Ion Torrent sequencing workflow were converted to FASTQ format using samtools fastq (SAMtools)^23^. The resulting FASTQ files were processed with MiXCR^24^ v4.4.2 using the thermofisher-human-rna-trb-oncomine-lr preset for the Oncomine™ TCR Beta-LR Assay. A floating left alignment boundary and a floating right alignment boundary on the C segment were used, and clonotypes were assembled on the basis of the CDR3 sequence with --assemble-clonotypes-by CDR3. The production command was:

mixcr analyze -Xmx50g thermofisher-human-rna-trb-oncomine-lr \

--floating-left-alignment-boundary \

--floating-right-alignment-boundary C \

--assemble-clonotypes-by CDR3 \ sample.fastq sample

For downstream ClonoDynamics analyses, amino-acid CDR3 sequence (aaSeqCDR3) and clonotype read count (readCount) were retained from the MiXCR clonotype tables. Rows sharing the same aaSeqCDR3 were collapsed by summing their readCount values, and readFraction was recalculated as the aggregated clonotype read count divided by the total read count of the corresponding repertoire. The resulting three-column clonotype tables (aaSeqCDR3, readCount, readFraction) constituted the exact processed repertoire inputs supplied to ClonoDynamics.

These processed input tables are deposited in the ClonoDynamics reproducibility dataset on Zenodo together with the analysis provenance and derived data, while the corresponding raw FASTQ files are available through the NCBI Sequence Read Archive.

### Computational workflow and reproducibility

All computational analyses were implemented in Python 3.9 within the modular ClonoDynamics framework for reconstructing longitudinal TCRβ clonotype dynamics from paired technical-replicate sequencing data. The source code, orchestration utilities, robustness modules, synthetic positive-control workflow, and figure-generation scripts are available at https://github.com/ClonoDynamics/clonodynamics-pipeline. The frozen software release corresponding to the version used in this study is archived in Zenodo under DOI 10.5281/zenodo.22925873. Detailed methodological definitions are provided in the Extended Methods, while the architecture of the computational workflow is summarized in Supplementary Figure 1 and Supplementary Data 1.

The workflow comprises seven main components: (i) repertoire characterization and measurement-noise-aware state inference from paired technical replicates; (ii) multirepresentation trajectory reconstruction and assembly of the longitudinal transition dataset; (iii) pseudo-longitudinal technical-reference construction and calibration of the primary replicate-resolved estimands; (iv) replicate-decoupled forward-dynamics analysis and comparison with the pseudo-longitudinal technical reference; (v) cross-replicate fluctuation analysis and temporal scaling across lags of 1–5 weeks; (vi) support-conditioned synthetic positive-control analyses used to assess recovery of known temporal accumulation under the empirical sampling and support structure; and (vii) robustness and sensitivity analyses addressing interval position, anchoring geometry, operational observation-domain selection, and cross-threshold dependence.

Execution is coordinated through a top-level interactive interface that separates Analysis and Figures workflows. The analysis orchestrator routes execution to the corresponding modular analysis blocks, resolves required upstream outputs, and writes results to standardized step-specific directories. Individual scientific modules can also be executed independently from the command line. Primary estimands and support definitions are frozen after the pseudo-longitudinal calibration stage and are then reused in the genuine longitudinal, synthetic-validation, and sensitivity analyses without re-optimization.

Intermediate datasets are written explicitly between stages. Large clonotype-, trajectory-, and transition-level tables are stored preferentially in Parquet format, whereas compact summaries, audit tables, manifests, run configurations, and provenance metadata are stored in CSV, TSV, or JSON format. Analysis parameters, random seeds, support definitions, subject universes, software versions, script hashes, and input metadata are recorded at the corresponding stages to preserve computational provenance.

Figure-generation scripts operate exclusively on finalized analysis outputs and do not recompute inferential quantities, bootstrap distributions, covariance estimates, temporal slopes, or synthetic-control results.

The exact processed repertoire inputs, the central longitudinal transition dataset, pseudo-reference configuration manifests, compact derived analysis outputs, synthetic-validation calibration and summaries, final Supplementary Data workbooks, and integrity checksums are archived in the ClonoDynamics reproducibility dataset on Zenodo under DOI 10.5281/zenodo.22925114. Raw FASTQ files remain deposited in the NCBI Sequence Read Archive and are not duplicated in the Zenodo archive.

### Repertoire characterization and latent-state inference from paired technical replicates

Repertoire abundance distributions were characterized independently in each technical replicate (Step 1). Continuous Pareto and lower-truncated log-normal models were compared at prespecified high-frequency tail cutoffs, following established approaches for the characterization and comparison of heavy-tailed distributions in immune repertoires^5,25,19^. This descriptive analysis neither excluded repertoires nor constrained latent-state inference.

For each complete subject–timepoint replicate pair, clonotypes detected in either replicate were retained, with zero counts for paired non-detections (Step 2). Counts were modelled as conditionally independent Negative-Binomial observations given a shared latent frequency, with a discrete power-law-shaped prior on a log-spaced frequency grid^11^. The prior exponent and dispersion parameter were jointly estimated by marginal likelihood for each pair^11^. The paired posterior median log-frequency provided the latent consensus coordinate, while observed replicate measurements remained distinct.

Posterior uncertainty, replicate agreement and posterior-predictive detectability were retained as diagnostics, not filters for successfully fitted states^26^. Operational observability was defined separately using the empirical lower-tail probability of fitted-model marginal log probability, with a reference threshold of α = 0.05. Only successful numerical fits entered the downstream state table; no longitudinal dynamical model was imposed. Full definitions are provided in Extended Methods 1, section (i).

### Reconstruction of clonotype trajectories and assembly of the longitudinal transition dataset

Clonotypes, operationally defined by amino-acid CDR3 sequence throughout this analysis, were assembled into within-subject trajectories and matched by amino-acid CDR3 sequence across available time pairs separated by 1–5 weeks (Steps 4–5), consistent with established longitudinal TCR-sequencing approaches for tracking individual clonotypes across serial samples^27,28,4,29^. Latent coordinates, observed replicate measurements and observation annotations remained distinct. Observed displacements required positive counts at both endpoints; no pseudo-counts were introduced.

Transition-level flags recorded each estimator’s measurement support: AB required replicate A at baseline and replicate B at both endpoints to be read-positive; BA used the reciprocal requirement. Cross-replicate covariance required both replicates to be read-positive at both endpoints (common4). These flags annotated rather than filtered the assembled table. Continuous endpoint observation statistics were retained for subsequent threshold-based classification. Coverage and structural consistency were audited in Step 6. Data contracts and optional posterior-uncertainty propagation are detailed in Extended Methods 1, section (ii).

### Pseudo-longitudinal technical calibration of conditioning and measurement representations

Technical benchmarking used 12 independently processed repertoire measurements from a single biological source, providing a reference in which between-measurement differences primarily reflected sampling and technical variability rather than longitudinal biological change^13,30^. The 12 measurements were arranged into 2,000 randomized configurations of six pseudo-timepoints and two replicates per timepoint. Pseudo-time represented technical ordering, not biological duration; the 2.5th–97.5th configuration percentiles therefore defined empirical randomization intervals rather than biological confidence intervals^31^.

Step 7 benchmarked observed replicate-decoupled forward dynamics and compared same-replicate and cross-replicate conditioning/displacement on matched common4 support to quantify measurement coupling. This design was motivated by the well-established problem that relating a baseline measurement to a change containing that same measurement can induce mathematical coupling and correlated measurement error^32^. Secondary diagnostics examined latent conditioning geometry.

Step 8 evaluated signed observed cross-replicate displacement covariance on common4, using latent midpoint abundance as the primary conditioning coordinate and alternative latent and observed coordinates as sensitivity analyses. The benchmark was used to identify estimand-specific conditioning and measurement representations rather than to define a universally preferred coordinate. Design, binning and randomization summaries are detailed in Extended Methods 1, section (iii).

### Replicate-decoupled forward-dynamics analysis and pseudo-longitudinal technical reference

At a one-week lag, the AB fold conditioned on observed baseline log-frequency in replicate A and measured displacement entirely in replicate B; BA reversed these roles (Step 9). Mean displacement profiles were estimated on their respective positivity supports in 20 shared observed-abundance bins, providing an abundance-resolved description of longitudinal clonotype dynamics^3^, and were then combined with exact 0.5/0.5 fold weighting. Operational observation classes did not restrict the primary analysis.

Ninety-five percent intervals were obtained from 2,000 biological-subject bootstrap resamples shared across folds, thereby preserving the within-subject dependence structure during resampling^33^. Linear slope and zero-crossing were descriptive summaries, supplemented by subject-specific, leave-one-subject-out and matched same-versus-cross checks. Step 10 compared genuine and pseudo profiles on shared absolute abundance support and in abundance-percentile space defined within subject/fold or configuration/fold, respectively. Biological bootstrap uncertainty remained distinct from pseudo-randomization variability; missing pseudo bins were not imputed, and comparisons did not extrapolate beyond shared support. Details are provided in Extended Methods 1, section (iv)

### Replicate-resolved fluctuation analysis and temporal scaling across lags of 1–5 weeks

Fluctuations were quantified as the signed sample covariance between observed replicate-specific log-frequency displacements, conditioned on latent midpoint abundance and restricted to *common4* (Step 11). Cross-replicate covariance was used to quantify fluctuation structure shared between replicate measurements, motivated by the standard additive-error result that errors independent of each other and of the underlying signal do not contribute to covariance^34^. Mean within-replicate variance and its excess over cross-replicate covariance were retained as comparators. Covariance was not truncated at zero, and no operational-class filter was applied.

Profiles across 1–5 weeks used 20 fixed latent-midpoint bins. Cohort covariance was reconstructed from pooled transition-level sufficient statistics, with uncertainty estimated using 2,000 subject-bootstrap resamples^33^. Step 12 compared covariance with the pseudo-reference on matched absolute midpoint support without imputing missing pseudo bins or subtracting the reference from longitudinal estimates.

Temporal analysis (Step 13) instead used a fixed contiguous abundance core and subjects with valid covariance in every core bin at every lag. Covariance was averaged equally across bins within subject, then equally across subjects. The signed slope across 1–5 weeks was estimated with 2,000 joint subject-bootstrap draws shared across lags. Analysis of fluctuation magnitude as a function of temporal separation has previously been used to characterize T-cell clone-size dynamics^3,8^. Constant, signed-linear and non-negative incremental-linear models were compared by the small-sample corrected Akaike information criterion (AICc) as secondary descriptors^35^. Full definitions and support criteria are provided in Extended Methods 1, section (v).

### Support-conditioned synthetic positive controls

Support-conditioned semi-synthetic controls were used to test recovery of prescribed temporal accumulation under known ground truth while retaining the analysis-eligible empirical sampling design, replicate depths, positivity masks and pair-specific dispersion parameters. Such experimentally calibrated simulations provide a means of evaluating computational methods under controlled signal while preserving data features relevant to their application domain^36,37^. Simulated frequencies combined visit-specific shocks with a cumulative random walk. A non-accumulating plateau and three positive doses targeted relative increases of 0.25, 0.50 and 1.00 in oracle log-frequency displacement variance between one and five weeks. Doses were calibrated from simulated frequencies before count generation, not from recovered estimates. Shared random fields linked scenarios, providing paired stochastic realizations across dose conditions^38^.

Each empirically positive clonotype received one read; remaining reads were allocated multinomially using Gamma-perturbed frequency weights, preserving replicate depth, richness and positivity support. Each scenario passed through production Steps 2, 4, 5, 11 and 13; simulated ground truth was withheld from inference. Recovery used the primary equal-bin/equal-subject slope on the empirical abundance grid and temporal core. Matched 2,000-draw subject bootstraps supported paired positive-dose-minus-plateau contrasts. This experiment assessed recovery conditional on the observed empirical support, rather than unconditional power across independently generated simulation realizations^36^. Mathematical definitions, calibration, generation and verification are provided in Extended Methods 2.

### Sensitivity analyses: calendar position, anchoring geometry, and observation-boundary effects

Step 14 evaluated calendar-position and common-start/common-end anchoring effects within the inherited temporal core and subject universe. Within-subject permutations assessed synchronized calendar-position structure while preserving the repeated-measures organization of the data^39^; joint subject bootstraps assessed anchored lag profiles, using 2,000 resamples for each procedure^33^. Multiple-testing adjustment for interval-position analyses used the Benjamini–Hochberg procedure^40^.

At α = 0.05, Step 15 compared the primary forward support with *T0PLUS* (TT + TF), TT and TF, and the primary *common4* fluctuation support with *NON_FF* (TT + TF + FT) and TT. Endpoint T/F labels indicated whether the stored empirical endpoint tail probability (p_value) was below the operational threshold; they did not indicate biological presence or absence.

Step 16 repeated the operational-domain analyses at α = 0.10, 0.05, 0.025 and 0.01 without refitting latent states or changing observed displacements, thereby testing sensitivity specifically to the operational observation boundary. Examining results under alternative reasonable definitions or cutoffs is a standard approach for assessing dependence of conclusions on analytic choices^41^. Paired subject bootstraps supported domain contrasts, with common abundance support used for cross-threshold comparisons. Temporal domain sensitivity used pooled covariance averaged equally across a common supported bin subset within the frozen subject universe; this separate sensitivity estimator did not replace the primary Step-13 equal-bin/equal-subject estimator. Full definitions are provided in Extended Methods 3.

## Supplemental information

**Supplementary Table 1.** Characteristics of enrolled participants and sampling overview.

**Document S1.** Supplementary Figures 1-11

**Document S2.** Extended Methods 1

**Document S3.** Extended Methods 2

**Document S4.** Extended Methods 3

**Supplementary Data 1. File Data_1.xlsx. ClonoDynamics computational architecture and module map.**

Manuscript-facing map of the current ClonoDynamics computational framework, linking analytical modules to their scientific roles, principal inputs and outputs, orchestration branches, analysis-versus-figure separation, randomization seeds, data-storage conventions, and reproducibility architecture.

**Supplementary Data 2. File Data_2.xlsx. Repertoire characterization and heavy-tail analysis.**

Replicate-level sequencing-depth and repertoire-richness summaries across the 114 TCRβ repertoires; Pareto tail fits and threshold-dependent Pareto-versus-lower-truncated-log-normal comparisons across prespecified empirical quantiles; paired technical-replicate concordance of fitted tail exponents; analysis-denominator and update checks; and source tables supporting Figure 2A–C.

**Supplementary Data 3. File Data_3.xlsx. Latent-state and observation-model diagnostics.**

Pair-specific latent-model parameters and fit-status tables; posterior uncertainty, replicate-posterior agreement, abundance-resolved quality metrics, ONE_POSITIVE versus TWO_POSITIVE state summaries, replicate-depth diagnostics, posterior-predictive detectability and operational-observation diagnostics, pair- and subject-level summaries, correlation analyses, and source data supporting Figure 2D–F and Supplementary Figure 2.

**Supplementary Data 4. File Data_4.xlsx. Longitudinal transition structure and estimand support.**

Characterization of the final longitudinal transition universe, including temporal-lag, subject and interval coverage; support for replicate-decoupled AB and BA forward estimands and common4 cross-replicate fluctuation analyses; TT/TF/FT/FF operational-class composition at the reference threshold; field-completeness, estimand-readiness and structural-consistency audits; and source data underlying Supplementary Figure 3.

**Supplementary Data 5. File Data_5.xlsx. Pseudo-longitudinal technical-null benchmarking.**

Configuration-level and ensemble analyses of the 2,000 pseudo-longitudinal randomization configurations generated from 12 independent technical repertoire measurements; replicate-decoupled AB/BA forward technical-null estimates, matched same-versus-cross controls, latent-conditioning geometry, common4 support, signed cross-replicate covariance and within-replicate variance components across pseudo-lags, conditioning sensitivities, temporal randomization summaries, configuration-level support and audit metrics, and source data underlying Figure 3 and Supplementary Figure 4.

**Supplementary Data 6. File Data_6.xlsx. Longitudinal replicate-resolved forward dynamics and technical-null comparison.**

Abundance-resolved observed AB, BA and symmetrically combined forward-displacement estimates at one-week lag, subject-cluster bootstrap intervals, matched same-versus cross-replicate controls, subject-specific and leave-one-subject-out robustness analyses, AB/BA fold concordance, absolute-abundance and within-unit percentile longitudinal-versus-pseudo technical-null comparisons, empirical randomization tests, fit-based crossover estimates, support diagnostics, and source data underlying Figure 4 and Supplementary Figure 5.

**Supplementary Data 7. File Data_7.xlsx. Longitudinal cross-replicate fluctuation dynamics and pseudo-longitudinal technical-null comparison.**

Abundance-resolved signed cross-replicate displacement covariance, mean within-replicate variance, replicate-specific excess and shared-fraction summaries across temporal lags of 1–5 weeks; common4 estimand support; biological subject-level fluctuation estimates; lag-specific absolute-abundance support matching between longitudinal and pseudo-longitudinal datasets; pointwise and abundance-averaged real-versus-pseudo covariance comparisons; empirical randomization tests, subject-bootstrap uncertainty, joint across-lag technical-null comparison, support and covariance-identity diagnostics, and source data underlying Figure 5 and Supplementary Figure 6.

**Supplementary Data 8. File Data_8.xlsx. Temporal scaling of longitudinal cross-replicate fluctuations.**

Fixed-abundance-core and complete-case-subject analyses of signed cross-replicate displacement covariance across temporal lags of 1–5 weeks; equal-bin/equal-subject primary estimates and covariance-degree-of-freedom-weighted sensitivity analyses; joint biological subject-bootstrap temporal slopes, subject-specific and abundance-resolved slopes, exact subject-level sign-flip tests, finite-lag constant, signed-linear and positive-incremental model comparisons, abundance-core and complete-case selection diagnostics, Step-14 analysis contract, and source data underlying Figure 6 and Supplementary Figure 7.

**Supplementary Data 9. File Data_9.xlsx. Support-conditioned synthetic positive-control validation.**

Oracle dose definitions and temporal variance profiles for the non-accumulating plateau and three positive-accumulation scenarios; recovered cross-replicate covariance profiles and temporal slopes on the frozen 17-bin abundance core and nine-subject complete-case cohort; scenario-minus-plateau paired slope increments, oracle increments and recovery fractions; subject-level temporal profiles; 2,000 matched biological subject-bootstrap slope distributions and paired contrasts; and source data underlying Figure 7.

**Supplementary Data 10. File Data_10.xlsx. Interval-position and subject-composition controls for temporal fluctuation scaling.**

Fixed-core calendar-position analyses following the primary Step-13 temporal-scaling result; within-subject permutation tests across interval start positions; abundance-resolved calendar-position sensitivity; common-start and common-end anchored temporal profiles; available-subject and matched-all-lag subject-composition controls; joint biological subject-bootstrap anchored slopes, subject-specific controls and matching diagnostics; and source data underlying Figure 8A and Supplementary Figure 9.

**Supplementary Data 11. File Data_11.xlsx. Operational observation-domain sensitivity and observation-threshold robustness.**

Fixed-threshold Step-15 analyses at reference alpha = 0.05, including TT/TF/FT/FF operational-class composition, observation-boundary characterization, replicate-decoupled forward dynamics across PRIMARY, T0PLUS, TT and TF domains, and cross-replicate fluctuation and temporal-scaling sensitivity across PRIMARY, NON_FF and TT domains; Step-16 cross-threshold analyses across alpha = 0.10, 0.05, 0.025, and 0.01, including common-support forward profiles and slopes, shared and replicate-specific fluctuation components, common-core temporal profiles and paired slope contrasts, reference-threshold replication audits and support diagnostics; and source data underlying Figure 8B–D and Supplementary Figures 10 and 11.

**Supplementary Data 12. File Data_12.xlsx.** SRA Accession for RepSeq data.

---

**Supplementary Table 1.** Characteristics of enrolled participants and sampling overview. Summary of demographic characteristics and sampling metrics for each participant, including age, gender, body mass index (BMI), number of blood withdrawals, and number of TCRβ repertoires successfully obtained.

| Participant ID | Age (years) | Gender | BMI, kg/m<sup>2</sup> | Number of withdrawals | Number of TCRβ repertoires* |
| --- | --- | --- | --- | --- | --- |
| #1 | 26 | M | 22.84 | 6 | 12 |
| #2 | 32 | M | 19.84 | 6 | 10 |
| #3 | 57 | F | 22.49 | 6 | 10 |
| #4 | 40 | M | 23.00 | 6 | 12 |
| #5 | 52 | F | 22.94 | 6 | 12 |
| #6 | 55 | F | 28.00 | 6 | 10 |
| #7 | 37 | F | 22.50 | 6 | 12 |
| #8 | 35 | F | 25.00 | 6 | 12 |
| #9 | 55 | M | 27.00 | 6 | 12 |
| #10 | 33 | M | 24.88 | 6 | 12 |

\* Repertoire counts refer to successfully generated measurements before exclusion of the non-converged latent-state fit for participant 5 at time point 2.

## References

1. Britanova, O.V., Shugay, M., Merzlyak, E.M., Staroverov, D.B., Putintseva, E.V., Turchaninova, M.A., Mamedov, I.Z., Pogorelyy, M.V., Bolotin, D.A., Izraelson, M., et al. (2016). Dynamics of Individual T Cell Repertoires: From Cord Blood to Centenarians. J. Immunol. 196, 5005–5013. 10.4049/jimmunol.1600005.

2. Chu, N.D., Bi, H.S., Emerson, R.O., Sherwood, A.M., Birnbaum, M.E., Robins, H.S., and Alm, E.J. (2019). Longitudinal immunosequencing in healthy people reveals persistent T cell receptors rich in highly public receptors. BMC Immunol. 20, 19. 10.1186/s12865-019-0300-5.

3. Bensouda Koraichi, M., Ferri, S., Walczak, A.M., and Mora, T. (2023). Inferring the T cell repertoire dynamics of healthy individuals. Proc. Natl. Acad. Sci. U. S. A. 120, e2207516120. 10.1073/pnas.2207516120.

4. Pogorelyy, M.V., Minervina, A.A., Touzel, M.P., Sycheva, A.L., Komech, E.A., Kovalenko, E.I., Karganova, G.G., Egorov, E.S., Komkov, A.Y., Chudakov, D.M., et al. (2018). Precise tracking of vaccine-responding T cell clones reveals convergent and personalized response in identical twins. Proc. Natl. Acad. Sci. U. S. A. 115, 12704–12709. 10.1073/pnas.1809642115.

5. Desponds, J., Mora, T., and Walczak, A.M. (2016). Fluctuating fitness shapes the clone-size distribution of immune repertoires. Proc. Natl. Acad. Sci. U. S. A. 113, 274–279. 10.1073/pnas.1512977112.

6. de Greef, P.C., Oakes, T., Gerritsen, B., Ismail, M., Heather, J.M., Hermsen, R., Chain, B., and de Boer, R.J. (2020). The naive T-cell receptor repertoire has an extremely broad distribution of clone sizes. eLife 9, e49900. 10.7554/eLife.49900.

7. Oakes, T., Heather, J.M., Best, K., Byng-Maddick, R., Husovsky, C., Ismail, M., Joshi, K., Maxwell, G., Noursadeghi, M., Riddell, N., et al. (2017). Quantitative Characterization of the T Cell Receptor Repertoire of Naïve and Memory Subsets Using an Integrated Experimental and Computational Pipeline Which Is Robust, Economical, and Versatile. Front. Immunol. 8, 1267. 10.3389/fimmu.2017.01267.

8. Gaimann, M.U., Nguyen, M., Desponds, J., and Mayer, A. (2020). Early life imprints the hierarchy of T cell clone sizes. eLife 9, e61639. 10.7554/eLife.61639.

9. Kawabe, T., Yi, J., and Sprent, J. (2021). Homeostasis of Naive and Memory T Lymphocytes. Cold Spring Harb. Perspect. Biol. 13, a037879. 10.1101/cshperspect.a037879.

10. Dessalles, R., Pan, Y., Xia, M., Maestrini, D., D’Orsogna, M.R., and Chou, T. (2022). How Naive T-Cell Clone Counts Are Shaped By Heterogeneous Thymic Output and Homeostatic Proliferation. Front. Immunol. 12, 735135. 10.3389/fimmu.2021.735135.

11. Puelma Touzel, M., Walczak, A.M., and Mora, T. (2020). Inferring the immune response from repertoire sequencing. PLoS Comput. Biol. 16, e1007873. 10.1371/journal.pcbi.1007873.

12. Mhanna, V., Bashour, H., Lê Quý, K., Barennes, P., Rawat, P., Greiff, V., and Mariotti-Ferrandiz, E. (2024). Adaptive immune receptor repertoire analysis. Nat. Rev. Methods Primers 4, 6. 10.1038/s43586-023-00284-1.

13. Barennes, P., Quiniou, V., Shugay, M., Egorov, E.S., Davydov, A.N., Chudakov, D.M., Uddin, I., Ismail, M., Oakes, T., Chain, B., et al. (2021). Benchmarking of T cell receptor repertoire profiling methods reveals large systematic biases. Nat. Biotechnol. 39, 236–245. 10.1038/s41587-020-0656-3.

14. Chaara, W., Gonzalez-Tort, A., Florez, L.-M., Klatzmann, D., Mariotti-Ferrandiz, E., and Six, A. (2018). RepSeq Data Representativeness and Robustness Assessment by Shannon Entropy. Front. Immunol. 9, 1038. 10.3389/fimmu.2018.01038.

15. Weber, C.R., Rubio, T., Wang, L., Zhang, W., Robert, P.A., Akbar, R., Snapkov, I., Wu, J., Kuijjer, M.L., Tarazona, S., et al. (2022). Reference-based comparison of adaptive immune receptor repertoires. Cell Rep. Methods 2, 100269. 10.1016/j.crmeth.2022.100269.

16. Koraichi, M.B., Touzel, M.P., Mazzolini, A., Mora, T., and Walczak, A.M. (2022). NoisET: Noise Learning and Expansion Detection of T-Cell Receptors. J. Phys. Chem. A 126, 7407– 7414. 10.1021/acs.jpca.2c05002.

17. Pavlova, A.V., Zvyagin, I.V., and Shugay, M. (2024). Detecting T-cell clonal expansions and quantifying clone survival using deep profiling of immune repertoires. Front. Immunol. 15, 1321603. 10.3389/fimmu.2024.1321603.

18. Tu, Y.-K., and Gilthorpe, M.S. (2007). Revisiting the relation between change and initial value: a review and evaluation. Stat. Med. 26, 443–457. 10.1002/sim.2538.

19. Clauset, A., Shalizi, C.R., and Newman, M.E.J. (2009). Power-Law Distributions in Empirical Data. SIAM Rev. 51, 661–703. 10.1137/070710111.

20. Mazzolini, A., Walczak, A.M., and Mora, T. (2026). Dynamics of memory B cells and plasmablasts in healthy individuals. Proc. Natl. Acad. Sci. U. S. A. 123, e2528141123. 10.1073/pnas.2528141123.

21. Aarsand, A.K., Røraas, T., Fernandez-Calle, P., Ricos, C., Díaz-Garzón, J., Jonker, N., Perich, C., González-Lao, E., Carobene, A., Minchinela, J., et al. (2018). The Biological Variation Data Critical Appraisal Checklist: A Standard for Evaluating Studies on Biological Variation. Clin. Chem. 64, 501–514. 10.1373/clinchem.2017.281808.

22. Iannone, F., Angotti, E., Lucia, F., Martino, L., Antico, G.C., Galato, F., Aversa, I., Gallo, R., Giordano, C., Abatino, A., et al. (2024). The biological variation of serum 1,25-dihydroxyvitamin D and parathyroid hormone, and plasma fibroblast growth factor 23 in healthy individuals. Clin. Chim. Acta Int. J. Clin. Chem. 557, 117863. 10.1016/j.cca.2024.117863.

23. Danecek, P., Bonfield, J.K., Liddle, J., Marshall, J., Ohan, V., Pollard, M.O., Whitwham, A., Keane, T., McCarthy, S.A., Davies, R.M., et al. (2021). Twelve years of SAMtools and BCFtools. GigaScience 10, giab008. 10.1093/gigascience/giab008.

24. Bolotin, D.A., Poslavsky, S., Mitrophanov, I., Shugay, M., Mamedov, I.Z., Putintseva, E.V., and Chudakov, D.M. (2015). MiXCR: software for comprehensive adaptive immunity profiling. Nat. Methods 12, 380–381. 10.1038/nmeth.3364.

25. Koch, H., Starenki, D., Cooper, S.J., Myers, R.M., and Li, Q. (2018). powerTCR: A model-based approach to comparative analysis of the clone size distribution of the T cell receptor repertoire. PLoS Comput. Biol. 14, e1006571. 10.1371/journal.pcbi.1006571.

26. Gelman, A., Meng, X.-L., and Stern, H. (1996). Posterior predictive assessment of model fitness via realized discrepancies. Stat. Sin. 6, 733–760.

27. DeWitt, W.S., Emerson, R.O., Lindau, P., Vignali, M., Snyder, T.M., Desmarais, C., Sanders, C., Utsugi, H., Warren, E.H., McElrath, J., et al. (2015). Dynamics of the cytotoxic T cell response to a model of acute viral infection. J. Virol. 89, 4517–4526. 10.1128/JVI.03474-14.

28. Shugay, M., Bagaev, D.V., Turchaninova, M.A., Bolotin, D.A., Britanova, O.V., Putintseva, E.V., Pogorelyy, M.V., Nazarov, V.I., Zvyagin, I.V., Kirgizova, V.I., et al. (2015). VDJtools: Unifying Post-analysis of T Cell Receptor Repertoires. PLOS Comput. Biol. 11, e1004503. 10.1371/journal.pcbi.1004503.

29. Minervina, A.A., Komech, E.A., Titov, A., Bensouda Koraichi, M., Rosati, E., Mamedov, I.Z., Franke, A., Efimov, G.A., Chudakov, D.M., Mora, T., et al. (2021). Longitudinal high-throughput TCR repertoire profiling reveals the dynamics of T-cell memory formation after mild COVID-19 infection. eLife 10, e63502. 10.7554/eLife.63502.

30. Trück, J., Eugster, A., Barennes, P., Tipton, C.M., Luning Prak, E.T., Bagnara, D., Soto, C., Sherkow, J.S., Payne, A.S., Lefranc, M.-P., et al. (2021). Biological controls for standardization and interpretation of adaptive immune receptor repertoire profiling. eLife 10, e66274. 10.7554/eLife.66274.

31. Ernst, M.D. (2004). Permutation Methods: A Basis for Exact Inference. Stat. Sci. 19, 676– 685. 10.1214/088342304000000396.

32. Chiolero, A., Paradis, G., Rich, B., and Hanley, J.A. (2013). Assessing the Relationship between the Baseline Value of a Continuous Variable and Subsequent Change Over Time. Front. Public Health 1, 29. 10.3389/fpubh.2013.00029.

33. Deen, M., and de Rooij, M. (2020). ClusterBootstrap: An R package for the analysis of hierarchical data using generalized linear models with the cluster bootstrap. Behav. Res. Methods 52, 572–590. 10.3758/s13428-019-01252-y.

34. Keogh, R.H., Shaw, P.A., Gustafson, P., Carroll, R.J., Deffner, V., Dodd, K.W., Küchenhoff, H., Tooze, J.A., Wallace, M.P., Kipnis, V., et al. (2020). STRATOS guidance document on measurement error and misclassification of variables in observational epidemiology: Part 1 – basic theory and simple methods of adjustment. Stat. Med. 39, 2197–2231. 10.1002/sim.8532.

35. Hurvich, C.M., and Tsai, C.-L. (1989). Regression and time series model selection in small samples. Biometrika 76, 297–307. 10.1093/biomet/76.2.297.

36. Morris, T.P., White, I.R., and Crowther, M.J. (2019). Using simulation studies to evaluate statistical methods. Stat. Med. 38, 2074–2102. 10.1002/sim.8086.

37. Sandve, G.K., and Greiff, V. (2022). Access to ground truth at unconstrained size makes simulated data as indispensable as experimental data for bioinformatics methods development and benchmarking. Bioinformatics 38, 4994–4996. 10.1093/bioinformatics/btac612.

38. Stout, N.K., and Goldie, S.J. (2008). Keeping the Noise Down: Common Random Numbers for Disease Simulation Modeling. Health Care Manag. Sci. 11, 399–406. 10.1007/s10729-008-9067-6.

39. Winkler, A., Webster, M., Vidaurre, D., Nichols, T., and Smith, S. (2015). Multi-level block permutation. NeuroImage 123, 253–268. 10.1016/j.neuroimage.2015.05.092.

40. Benjamini, Y., and Hochberg, Y. (1995). Controlling the False Discovery Rate: A Practical and Powerful Approach to Multiple Testing. J. R. Stat. Soc. Ser. B Methodol. 57, 289–300. 10.1111/j.2517-6161.1995.tb02031.x.

41. Thabane, L., Mbuagbaw, L., Zhang, S., Samaan, Z., Marcucci, M., Ye, C., Thabane, M., Giangregorio, L., Dennis, B., Kosa, D., et al. (2013). A tutorial on sensitivity analyses in clinical trials: the what, why, when and how. BMC Med. Res. Methodol. 13, 92. 10.1186/1471-2288-13-92.

