## Supplementary Figures for "ClonoDynamics enables replicate-resolved inference of longitudinal T cell receptor repertoire dynamics"

**Supplementary Figure S1. Computational architecture and dependency structure of the ClonoDynamics pipeline**

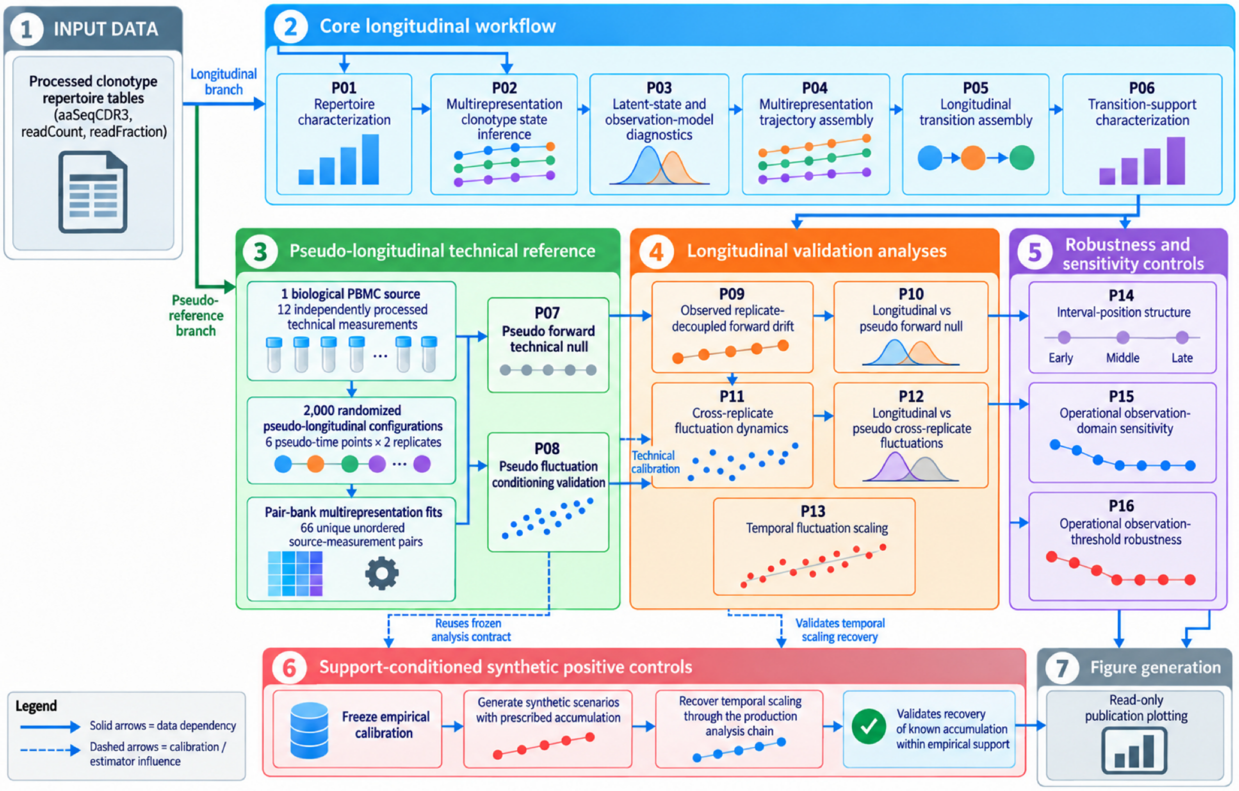


**Supplementary Figure 1. Computational architecture and dependency structure of the ClonoDynamics pipeline.**

Processed clonotype repertoire tables provide the inputs to the descriptive repertoire-characterization module (P01) and to the replicate-resolved longitudinal workflow comprising state inference and diagnostics, multirepresentation trajectory reconstruction, transition assembly, and support characterization (P02–P06). The independent pseudo-longitudinal technical reference is constructed from 12 technical measurements of a single biological PBMC source and analysed through the pseudo-forward (P07) and pseudo-fluctuation (P08) benchmarks, which inform the primary estimator choices used downstream.

Genuine longitudinal analyses comprise replicate-decoupled forward dynamics and comparison with the corresponding pseudo-reference (P09–P10), cross-replicate fluctuation dynamics and pseudo-reference comparison (P11–P12), and fixed-support temporal scaling (P13). Interval-position and anchoring controls (P14), operational observation-domain sensitivity (P15), and observation-threshold robustness (P16) assess the stability of the primary results without redefining the primary estimands.

A separate support-conditioned synthetic positive-control branch freezes the empirical calibration, generates scenarios with prescribed temporal accumulation, and evaluates recovery through the production analysis chain. Figure-generation modules operate downstream on finalized numerical outputs. Solid arrows indicate direct data dependencies; dashed arrows indicate calibration or estimator-definition influences.

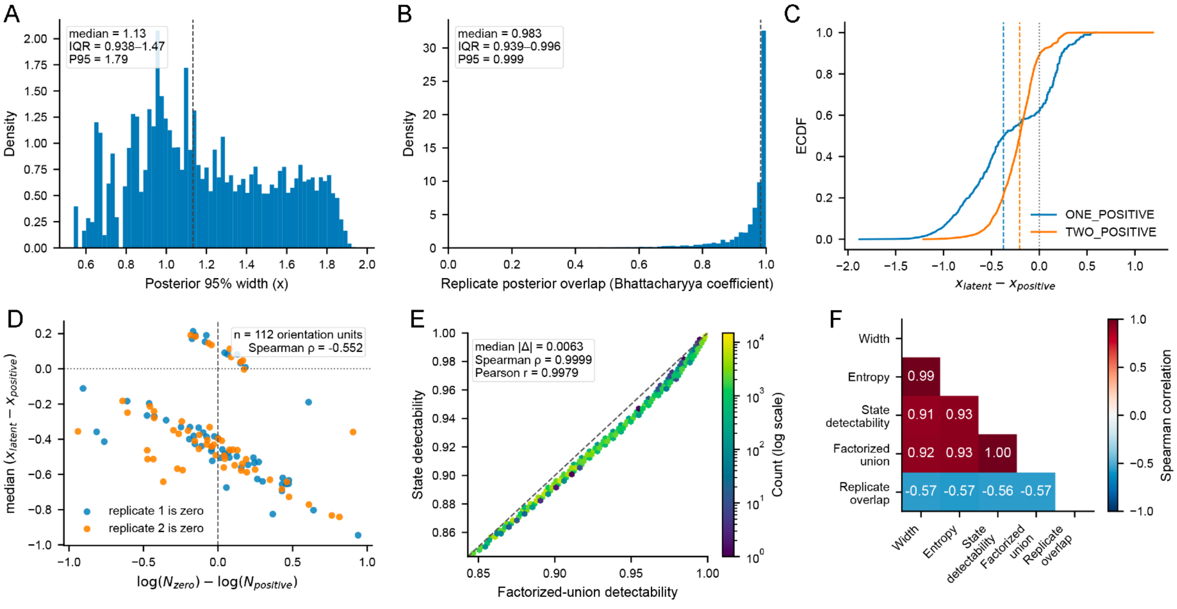


**Supplementary Figure 2. Extended diagnostics of latent-state and observation-model inference.**

(A) Distribution of posterior 95% credible-interval width in latent log-frequency space across clonotype–timepoint states. The dashed vertical line marks the overall median; the inset reports the median, interquartile range and 95th percentile.

(B) Distribution of agreement between the two single-replicate latent posteriors, quantified by the exact discrete-posterior Bhattacharyya coefficient. The dashed vertical line marks the overall median; the inset reports the median, interquartile range and 95th percentile.

(C) Empirical cumulative distributions of $x_{\text{latent}} - x_{\text{positive}}$ for ONE_POSITIVE and TWO_POSITIVE states. Colored dashed vertical lines indicate the corresponding medians, and the gray dotted line denotes zero difference.

(D) Relationship between relative sequencing depth and the latent-versus-positive representation shift within ONE_POSITIVE states. Each point represents one subject × time × zero-replicate orientation; colors indicate whether replicate 1 or replicate 2 was the zero-count replicate. The x-axis shows $\log(N_{\text{zero}}/N_{\text{positive}})$, and the y-axis shows the median $x_{\text{latent}} - x_{\text{positive}}$ within that orientation. Dashed vertical and dotted horizontal lines denote zero. The inset reports the number of orientation units and the Spearman correlation.

(E) Posterior-predictive state detectability versus factorized-union detectability. Hexagon color indicates the number of clonotype–timepoint states on a logarithmic scale, and the dashed line denotes identity. The inset reports the median absolute difference and the Spearman and Pearson correlations.

(F) Pairwise Spearman correlations among posterior credible-interval width, posterior entropy, state detectability, factorized-union detectability and replicate-posterior overlap. The lower triangular correlation matrix is shown, with correlation coefficients reported within cells.

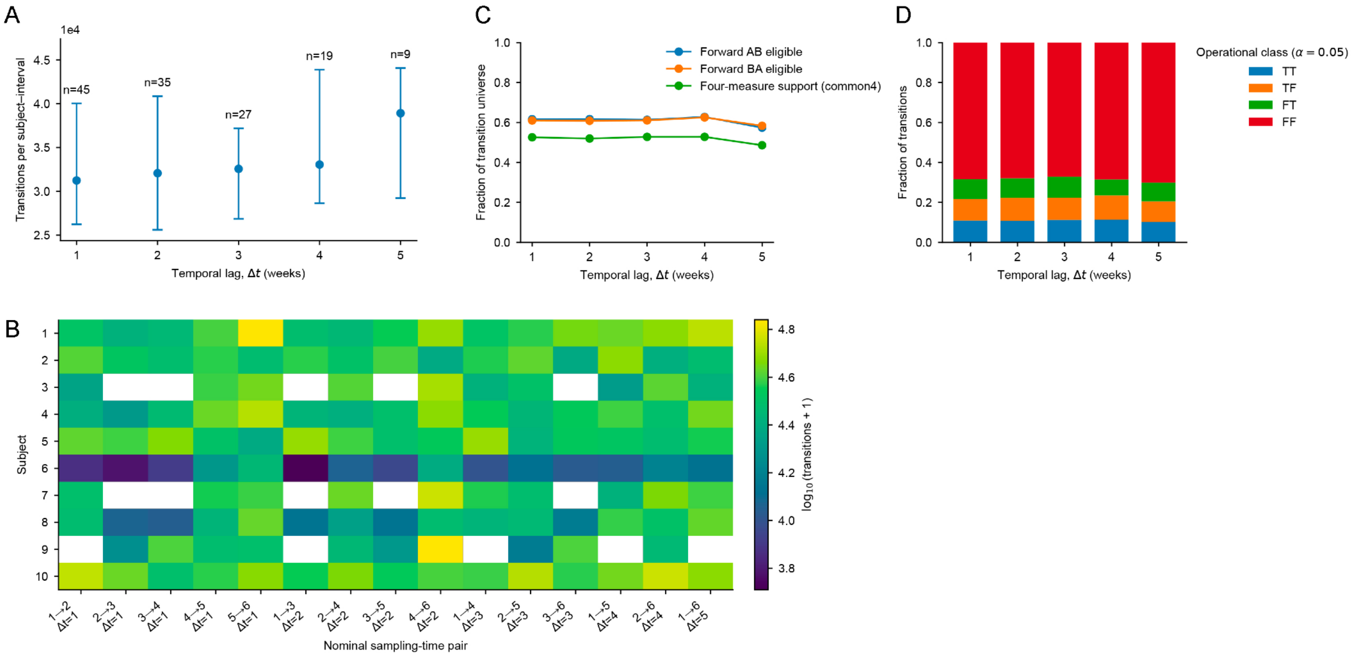


**Supplementary Figure 3. Structure and estimand support of the longitudinal transition dataset.**

(A) Transition support within available subject-specific intervals across temporal lags from 1 to 5 weeks. Points indicate the median number of clonotype transitions per subject–interval and error bars the corresponding interquartile range. Labels indicate the number of contributing subject–interval pairs at each lag.

(B) Subject-by-interval coverage matrix for the 15 nominal sampling-time pairs. Color represents log10(transitions + 1); white cells denote subject–interval combinations unavailable because one or both endpoint subject–timepoint representations were not available for downstream longitudinal analysis. Of the 150 subject–interval combinations possible under the complete six-timepoint design, 130 were available for longitudinal analysis. This reflects the three technically unsuccessful subject–timepoint samples and one additional subject–timepoint pair (subject 5, time point 2) for which latent-state fitting did not converge.

(C) Fraction of the assembled longitudinal transition dataset eligible for the replicate-decoupled forward AB and BA estimands and for the four-measure (*common4*) support used for cross-replicate fluctuation analysis, shown across temporal lags. Across all transitions, 52.40% and 52.44% were eligible for the AB and BA orientations, respectively, and 41.87% satisfied the *common4* criterion.

(D) Relative composition of reference operational endpoint-observation classes across temporal lags, defined at the reference threshold α = 0.05. TT denotes transitions operationally observable at both endpoints, TF and FT denote transitions crossing the operational observation boundary, and FF denotes transitions unobservable at both endpoints. Across the complete transition dataset, TT transitions represented 11.82%, TF+FT transitions 20.39%, and FF transitions 67.78%. These classes describe the operational observation domain and were retained as transition annotations rather than used as state-quality filters.

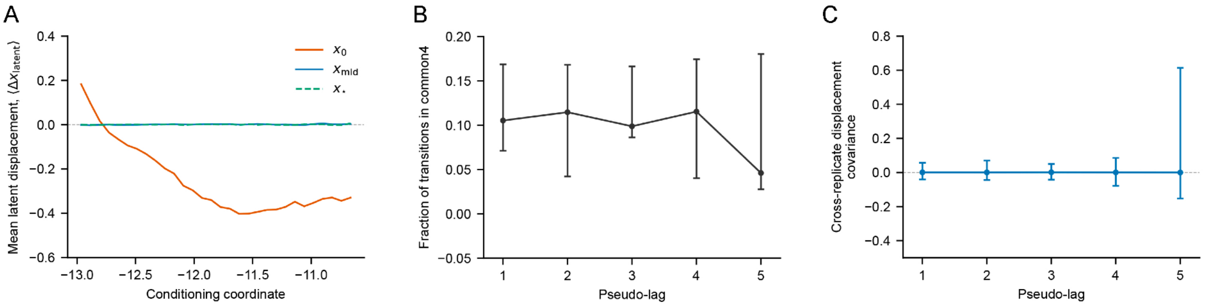


**Supplementary Figure 4. Conditioning-coordinate diagnostics, transition support and pooled covariance in the pseudo-longitudinal technical reference.**

Lines and points show medians across 2,000 randomization configurations; error bars (B,C) indicate the corresponding 2.5th–97.5th randomization percentiles. Horizontal reference lines in A and C mark zero.

(A) Mean latent log-frequency change as a function of the conditioning coordinate: initial latent abundance, $x_0$ (orange), or the transition-centred coordinates $x_{mid}$ (blue) and $x_*$ (green dashed).

(B) Fraction of all transitions meeting *common4* criteria within each configuration, summarized by pseudo-lag.

(C) Cross-replicate displacement covariance, $\mathrm{Cov}(\Delta, x_A \Delta x_B)$, calculated by pooling *common4* transitions within each configuration and pseudo-lag, without abundance binning.

Source data are provided in Supplementary Data 5.

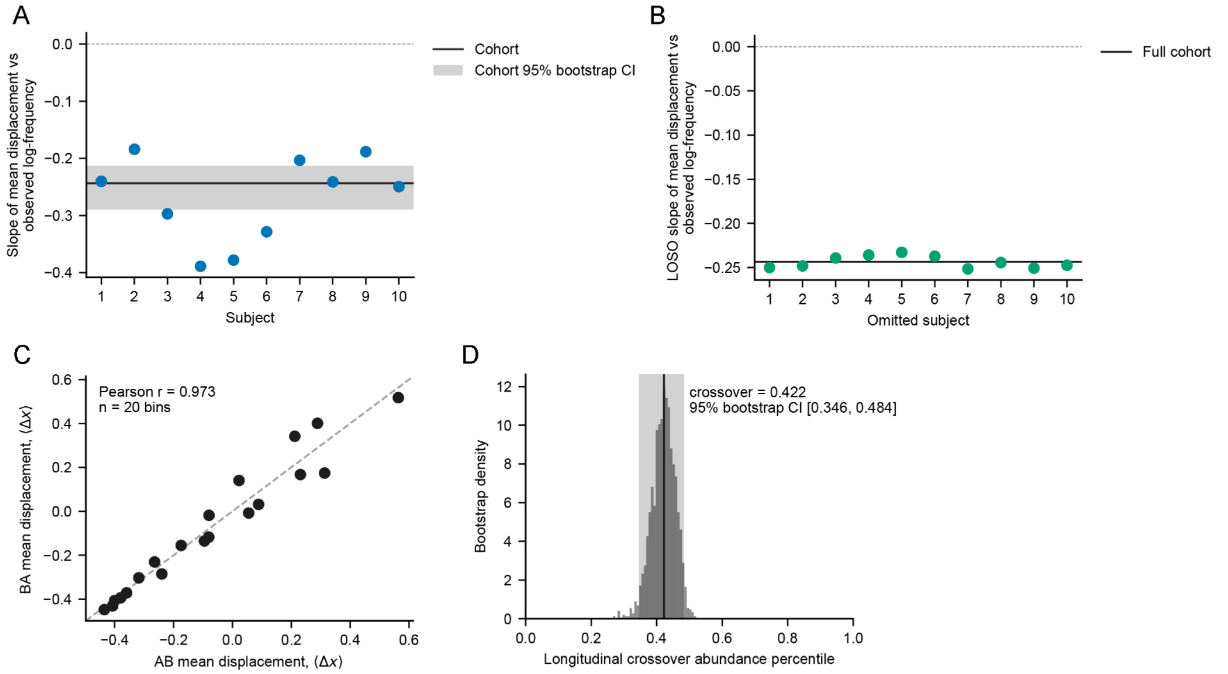


**Supplementary Figure 5. Subject-level robustness and concordance of longitudinal forward dynamics.**

(A) Subject-specific slopes of mean observed displacement versus initial observed log-frequency. The horizontal line indicates the cohort estimate and the shaded band its 95% subject-bootstrap interval.

(B) Leave-one-subject-out slopes, with the full-cohort estimate shown as a horizontal reference line.

(C) Bin-wise concordance between the AB and BA mean-displacement profiles; the dashed line denotes identity.

(D) Subject-bootstrap distribution of the longitudinal crossover abundance percentile. The vertical line marks the point estimate and the shaded region its 95% bootstrap interval.

Source data are provided in Supplementary Data 6.

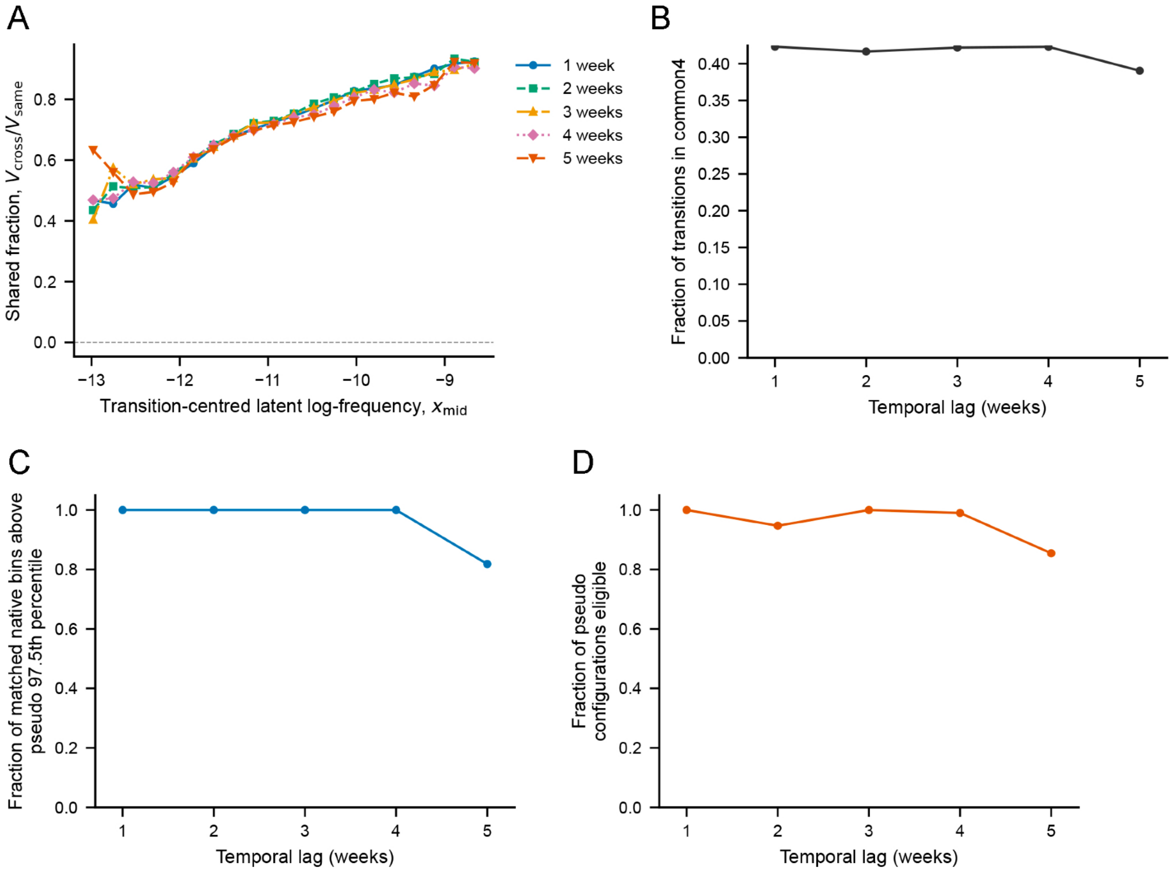


**Supplementary Figure 6. Shared fluctuation ratio and support diagnostics across temporal lags.**

(A) Shared-fluctuation ratio, defined as cross-replicate covariance divided by mean within-replicate displacement variance, across transition-centred latent log-frequency, $x_{mid}$, for temporal lags of 1–5 weeks.

(B) Fraction of longitudinal transitions meeting the *common4* support criterion at each temporal lag.

(C) Fraction of matched abundance bins in which longitudinal cross-replicate covariance exceeded the pseudo 97.5th percentile.

(D) Fraction of pseudo configurations eligible for the lag-specific matched-support comparison.

Source data are provided in Supplementary Data 7.

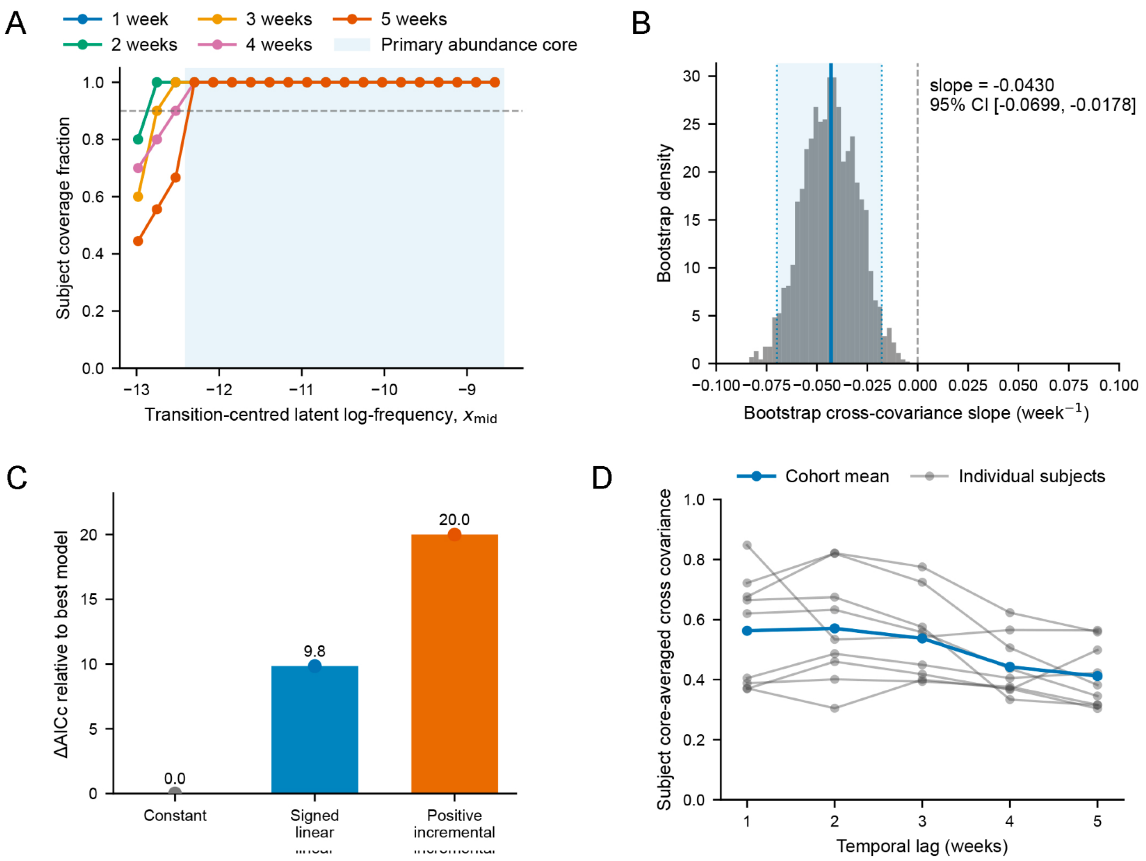


**Supplementary Figure 7. Support, bootstrap uncertainty and robustness of temporal fluctuation scaling.**

(A) Subject-coverage fraction across the fixed $x_{mid}$ abundance grid for temporal lags of 1–5 weeks. Shading indicates the abundance core retained for the primary temporal-scaling analysis.

(B) Subject-bootstrap distribution of the temporal slope of cross-replicate covariance. Vertical lines indicate the observed slope, its 95% bootstrap interval and zero.

(C) Shared-fluctuation ratio, defined as cross-replicate covariance divided by mean within-replicate displacement variance.

(D) Subject-specific core-averaged cross-replicate covariance across temporal lags, with the cohort mean shown in blue.

Source data are provided in Supplementary Data 8.

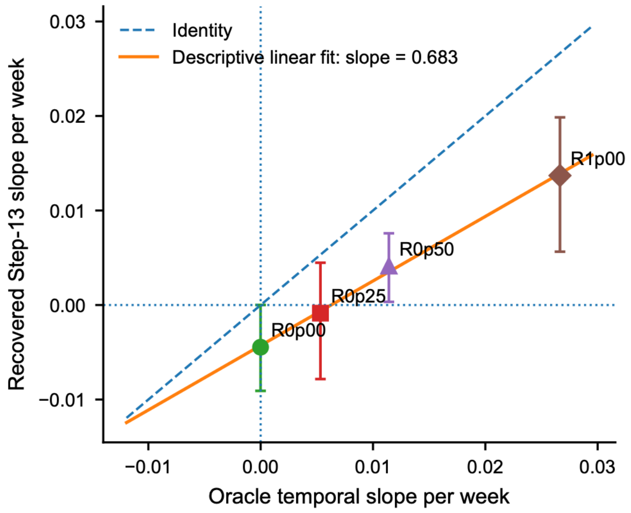


**Supplementary Figure 8. Oracle versus recovered temporal scaling in the support-conditioned synthetic controls.**

Oracle temporal slopes of simulated log-frequency displacement variance are plotted against the corresponding temporal slopes of cross-replicate covariance recovered after synthetic count generation and analysis through the same Step-13 workflow used for the empirical data. Points denote the recovered slope estimates for the four accumulation scenarios, and vertical error bars indicate their 95% subject-bootstrap confidence intervals. The dashed line denotes identity, and the solid line shows the descriptive linear fit across the four scenarios. Horizontal and vertical dotted lines mark zero slope.

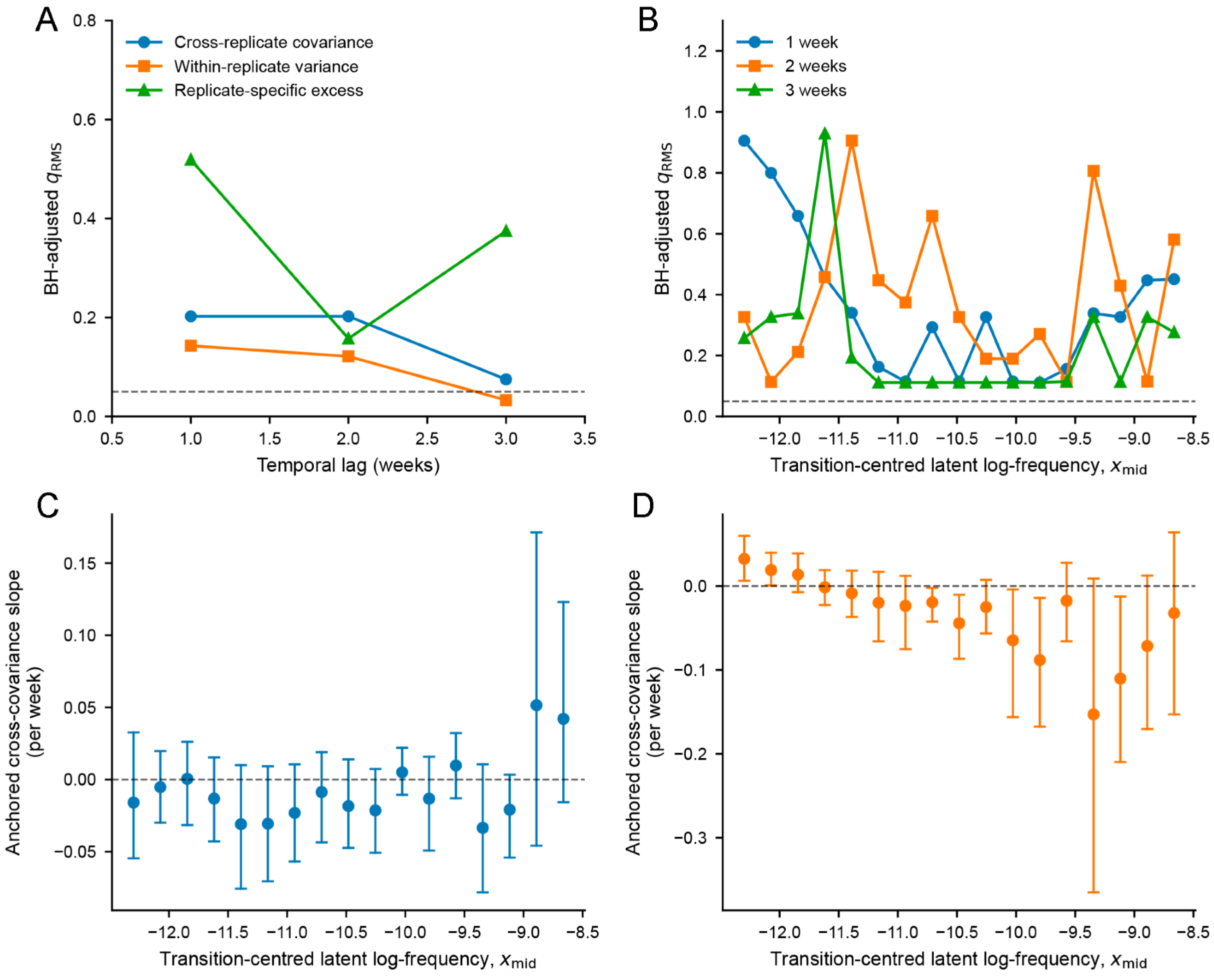


**Supplementary Figure 9. Interval-position and anchored temporal controls.**

**(A)** Benjamini–Hochberg-adjusted permutation q-values ($q_{\mathrm{RMS}}$) for the root-mean-square calendar-position statistic across temporal lags of 1–3 weeks for cross-replicate covariance, within-replicate variance and replicate-specific excess. The horizontal dashed line indicates $q = 0.05$.

**(B)** Abundance-resolved BH-adjusted $q_{\mathrm{RMS}}$ values for cross-replicate covariance at temporal lags of 1, 2 and 3 weeks as a function of transition-centred latent log-frequency, $x_{\mathrm{mid}}$.

**(C)** Abundance-resolved temporal slopes of cross-replicate covariance for the matched common-start control. Points indicate the estimated slope within each abundance bin and vertical bars indicate the corresponding 95% subject-bootstrap intervals.

**(D)** Abundance-resolved temporal slopes of cross-replicate covariance for the matched common-end control, displayed as in panel C. Horizontal dashed lines in C and D indicate zero slope.

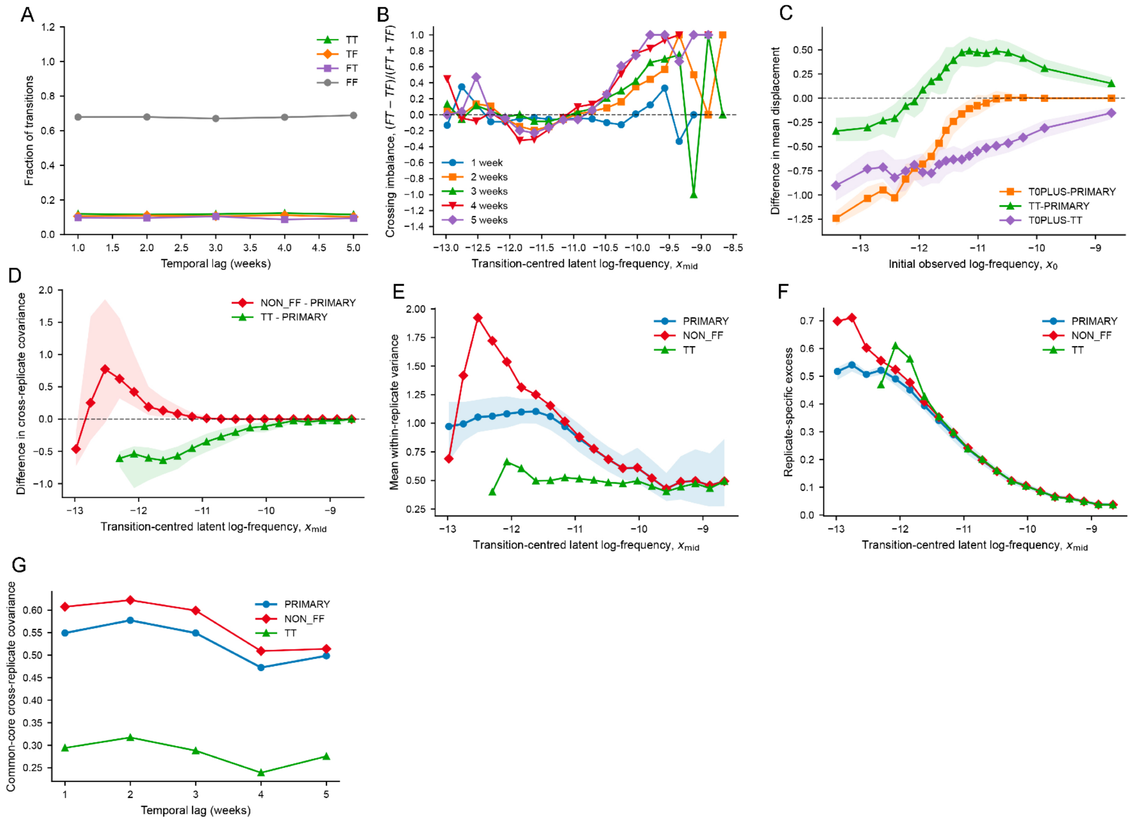


**Supplementary Figure 10. Fixed-threshold operational-domain sensitivity at $\alpha = 0.05$.**

TT, TF, FT and FF denote the operational endpoint classes at α = 0.05; T0PLUS = TT + TF, NON_FF = TT + TF + FT, and PRIMARY denotes the estimator-specific primary support. T/F labels refer to the operational observation boundary, not biological presence or absence.

**(A)** Fractions of TT, TF, FT and FF transitions across temporal lags of 1–5 weeks.

**(B)** Boundary-crossing imbalance, $(FT - TF)/(FT + TF)$, as a function of transition-centred latent log-frequency, $x_{\text{mid}}$, for temporal lags of 1–5 weeks.

**(C)** Differences between one-week forward profiles for T0PLUS–PRIMARY, TT–PRIMARY and T0PLUS–TT as a function of initial observed log-frequency, $x_0$. Shaded regions indicate the corresponding paired subject-bootstrap intervals.

**(D)** Differences in one-week cross-replicate covariance for NON_FF and TT relative to PRIMARY across $x_{\text{mid}}$, with paired subject-bootstrap intervals.

**(E)** Mean within-replicate variance at one week for PRIMARY, NON_FF and TT as a function of $x_{\text{mid}}$; the shaded region indicates the PRIMARY subject-bootstrap interval.

**(F)** Replicate-specific excess at one week for PRIMARY, NON_FF and TT across $x_{\text{mid}}$.

**(G)** Pooled common-core cross-replicate covariance across temporal lags of 1–5 weeks for PRIMARY, NON_FF and TT, using the separate temporal-domain sensitivity estimator. Dashed horizontal lines indicate zero where shown.

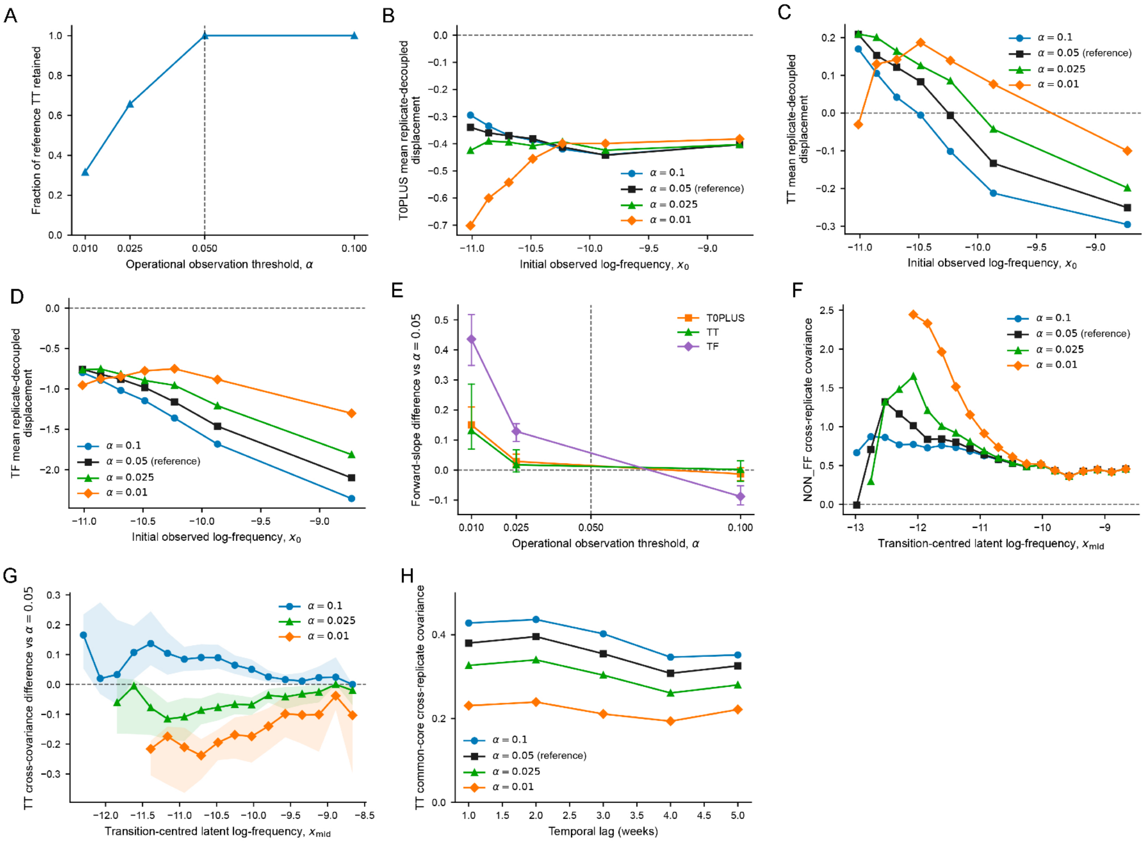


**Supplementary Figure 11. Cross-threshold operational robustness.**

**(A)** Fraction of transitions classified as TT at the reference threshold α = 0.05 that remain TT at each operational observation threshold (α = 0.01, 0.025, 0.05 and 0.10); the vertical dashed line marks the reference threshold $\alpha = 0.05$.

**(B–D)** One-week replicate-decoupled forward profiles for T0PLUS, TT and TF, respectively, across the operational thresholds shown, plotted as a function of initial observed log-frequency, $x_0$.

**(E)** Differences in forward slope relative to the reference threshold $\alpha = 0.05$ for T0PLUS, TT and TF; vertical bars indicate 95% paired subject-subject bootstrap intervals.

**(F)** One-week NON_FF cross-replicate covariance profiles across operational thresholds as a function of transition-centred latent log-frequency, $x_{\mathrm{mid}}$.

**(G)** Differences in one-week TT cross-replicate covariance relative to $\alpha = 0.05$ across $x_{\mathrm{mid}}$; shaded regions indicate 95% paired subject-bootstrap intervals where available.

**(H)** TT pooled common-core cross-replicate covariance across temporal lags of 1–5 weeks for the four operational thresholds, using the temporal-domain sensitivity estimator.
