## Extended_Methods_3 for "ClonoDynamics enables replicate-resolved inference of longitudinal T cell receptor repertoire dynamics"

### 1. Scope and frozen upstream architecture

The sensitivity analyses were designed to test whether the longitudinal conclusions depended on interval composition or on the operational observation domain after the main estimands had already been defined. They did not introduce alternative latent-state models or alternative dynamical outcomes.

The inherited fluctuation architecture was the finalized Step-11 estimand

$$V_{\text{cross}}(x, \Delta t) = \mathrm{Cov}(\Delta x_A, \Delta x_B \mid x_{\text{mid}}^{\text{latent}} = x, \Delta t, \text{common4}),$$

where

$$\Delta x_A = \Delta x_{\text{observed,rep1}}, \qquad \Delta x_B = \Delta x_{\text{observed,rep2}}.$$

The within-replicate comparator and replicate-specific excess were retained as

$$V_{\text{same}} = \frac{1}{2}\left[\mathrm{Var}(\Delta x_A) + \mathrm{Var}(\Delta x_B)\right],$$

$$V_{\text{excess}} = V_{\text{same}} - V_{\text{cross}}.$$

Step 14 inherited both the fixed `xmid_latent` abundance core and the complete-case biological-subject set selected by Step 13. It therefore tested composition sensitivity on the same support used for the primary temporal analysis rather than selecting a new abundance domain. The finalized `14-interval_position_structure.py` implementation explicitly rejects legacy TT-filtered, `xstar_latent`-conditioned variance/MSD inputs.

### 2. Calendar-position analysis

Step 14 read the subject × physical-interval × abundance-bin covariance table produced by Step 11. Let $\mathcal{B}$ denote the fixed Step-13 abundance core. For subject $s$ and physical interval $p = (t_0, t_1)$ at lag $h$, the core-level value for metric $m$ was

$$y_{sp}^{(m)} = \frac{1}{|\mathcal{B}|}\sum_{b \in \mathcal{B}} m_{spb},$$

provided every selected core bin satisfied the subject-interval support requirement. By default, the minimum transition count per subject × interval × bin inherited the Step-13 subject-cell requirement; the production value was the mathematical covariance minimum of two transitions.

For a fixed lag $h$, intervals were aligned by their nominal start position $t_0$. At calendar position $p$, the cohort profile was the equal-subject mean

$$\bar{y}_p = \frac{1}{n_p}\sum_{s \in \mathcal{S}_p} y_{sp},$$

where $\mathcal{S}_p$ is the set of complete-case Step-13 subjects with a finite value at that position. Calendar positions were retained only when the prespecified minimum number of subjects was available.

The primary synchronized-position statistic was

$$T_{\text{RMS}} = \sqrt{\frac{1}{P}\sum_{p=1}^{P}\left(\bar{y}_p - \bar{\bar{y}}\right)^2},$$

where $P$ is the number of retained positions. A secondary statistic was the absolute ordinary-least-squares slope of $\bar{y}_p$ versus $t_0$.

The permutation null preserved each subject's observed set of finite interval values and missingness pattern. For each subject independently, finite values were randomly permuted among that subject's available calendar positions; the missing positions were not filled. The cohort profile and both statistics were then recalculated. The procedure therefore preserved subject identity, lag, metric distribution, fixed abundance composition, and subject-specific missingness while removing synchronized association with calendar position. Restricting permutations to rearrangements compatible with the repeated-measures dependence structure follows the general exchangeability principle for structured permutation inference (Winkler et al., 2015).

For $B = 2000$ permutations, an upper-tail empirical P value for statistic $T$ was calculated with the finite-sample +1 correction (Phipson and Smyth, 2010):

$$p_{\text{emp}} = \frac{1 + \sum_{b=1}^{B} \mathbf{1}(T_b^* \geq T_{\text{obs}})}{B + 1}.$$

Benjamini–Hochberg false-discovery-rate correction was performed separately for the RMS and absolute-slope statistics (Benjamini and Hochberg, 1995). For the primary core-level analysis, correction was applied within metric across valid lags. A secondary abundance-resolved analysis repeated the same permutation design within individual core bins and applied correction within metric across all valid lag × bin tests.

#### 3. Common-start and common-end anchoring

Anchoring controlled the composition of physical intervals contributing at different lags. Two geometries were defined:

- `common_start`: transitions with $t_0 = \min(t_0)$;
- `common_end`: transitions with $t_1 = \max(t_1)$.

For each subject, anchor, and lag, the metric was averaged equally across the same frozen Step-13 abundance core. Two subject-composition modes were evaluated. In `available` mode, all Step-13 complete-case subjects with a finite anchored value at a given lag contributed, so subject composition could vary across lag. In `matched_all_dt` mode, a subject was retained only when the anchored value was finite at every requested lag, producing an identical subject set across the lag series.

For each anchored profile, the signed descriptive model

$$M(\Delta t) = K + D(\Delta t - 1)$$

was fitted by ordinary least squares, with at least four finite lag values required. Biological uncertainty was quantified using $B = 2000$ subject-cluster bootstrap replicates, preserving the biological subject as the resampling unit (Deen and de Rooij, 2020). A subject sample was drawn once per replicate and its multiplicities were propagated jointly across all lags before refitting $D$. Percentile intervals and the fractions of bootstrap slopes above and below zero were reported. For `matched_all_dt`, subject-specific slopes and exact sign-flip tests were additionally retained as conservative subject-level diagnostics. Step 14 did not rerun the Step-13 finite-lag model comparison.

#### 4. Operational observation classes at the reference threshold

Step 15 used the empirical endpoint observation statistic from Step 5. At the fixed reference threshold $\alpha = 0.05$,

$$T_k = \mathbf{1}[p_k < \alpha], \qquad F_k = 1 - T_k,$$

with endpoint classes

$$TT = (T_0, T_1), \quad TF = (T_0, F_1), \quad FT = (F_0, T_1), \quad FF = (F_0, F_1).$$

These labels were operational observation states. They were not interpreted as biological presence, disappearance, extinction, or persistence.

Step 15 first characterized class composition by lag and across the fixed Step-11 `xmid_latent` abundance grid. Within the non-FF population, the crossing fraction was

$$f_{\text{cross}} = \frac{n_{TF} + n_{FT}}{n_{TT} + n_{TF} + n_{FT}},$$

and, when at least one crossing occurred, the directional imbalance was

$$I_{\text{cross}} = \frac{n_{FT} - n_{TF}}{n_{TF} + n_{FT}}.$$

These quantities describe traversal of the operational observation boundary only.

#### 5. Forward observation-domain sensitivity

The forward estimator itself was unchanged from Step 9. With technical replicates A and B,

$$AB: \quad x_A^{\text{obs}}(t_0) \longrightarrow \Delta x_B^{\text{obs}},$$

$$BA: \quad x_B^{\text{obs}}(t_0) \longrightarrow \Delta x_A^{\text{obs}}.$$

AB and BA were estimated separately and combined only after binning with exact equal fold weight. The fixed Step-9 observed-abundance grid was retained.

Four operational domains were evaluated:

- `PRIMARY`: the manuscript-primary Step-9 estimand-specific support, with no operational-class filter;
- `TOPLUS`: operationally T at $t_0$, with the future endpoint unrestricted, i.e. $TT \cup TF$;
- `TT`: operationally T at both endpoints;
- `TF`: the descriptive boundary-crossing component of `TOPLUS`.

A combined abundance bin was retained only when both reciprocal folds satisfied the support threshold; no single-fold fallback was allowed. One subject-cluster bootstrap was shared across all folds and operational domains, enabling paired contrasts among `TOPLUS`, `TT`, and the invariant primary curve. The principal contrasts were `TOPLUS-PRIMARY`, `TT-PRIMARY`, and `TOPLUS-TT`.

#### 6. Fluctuation observation-domain sensitivity

Fluctuation-domain sensitivity retained the Step-11 variables and grid exactly. The base support remained `common4`, after which operational classes were used only to define nested analysis domains:

- `PRIMARY`: all `common4` transitions;
- `NON_FF`: `common4` transitions in $TT \cup TF \cup FT$;
- `TT`: `common4` transitions classified TT.

Within every subject × lag × abundance-bin cell, sufficient statistics were accumulated for `cross_cov`, `same_var_mean`, and `replicate_specific_excess`. Pooled cohort estimates were reconstructed by aggregating those sufficient statistics across subjects. The same joint subject bootstrap was propagated across `PRIMARY`, `NON_FF`, and `TT`, allowing paired domain differences. A bootstrap cell contributed only if the reconstructed transition and subject support thresholds remained satisfied.

This analysis quantified how operational-domain selection changed the amplitude or abundance dependence of the shared fluctuation estimand. It did not interpret differences between domains as an additive decomposition into technical and biological variance.

#### 7. Temporal sensitivity to the operational domain

The Step-13/14 complete-case subject universe and abundance core were frozen before applying operational-domain restrictions. Filtering to `NON_FF` or `TT` can remove support from individual subject × lag × abundance cells, so Step 15 did not replace the primary Step-13 equal-bin/equal-subject estimator with a silently reselected cohort.

Instead, a matched sensitivity estimand was constructed. Starting from the frozen core, the largest contiguous bin subset with valid pooled covariance support for `PRIMARY`, `NON_FF`, and `TT` at every requested lag was selected. The biological-subject universe remained the Step-14 universe. Pooled covariance was computed within each retained bin, bins were averaged equally within lag, and a signed slope

$$M(\Delta t) = K + D(\Delta t - 1)$$

was fitted separately for each operational domain. The same subject bootstrap draw was used across domains and lags. A bootstrap temporal slope was retained only when the entire fixed common operational core satisfied the declared support thresholds at every lag. This `matched_core_pooled_covariance_equal_bin` estimator was a sensitivity estimator and was not substituted for the primary Step-13 temporal estimator.

#### 8. Numerical observation-threshold robustness

Step 16 varied the operational threshold over the prespecified grid below as a sensitivity analysis of the operational observation boundary, rather than as threshold optimization (Thabane et al., 2013):

$$\alpha \in \{0.10,\ 0.05,\ 0.025,\ 0.01\},$$

with $\alpha_{\text{ref}} = 0.05$. For each endpoint,

$$T_k(\alpha) = \mathbf{1}[p_k < \alpha].$$

Only these endpoint labels and the derived TT/TF/FT/FF transition classes were recalculated. Pair-specific latent-state fits, observed replicate frequencies, replicate displacements, posterior uncertainty, and the primary Step-9/Step-11 estimands were not recomputed.

Step 16 called the finalized Step-15 implementation at every $\alpha$, preserving the same support rules, abundance grids, subject universe, bootstrap seed, and bootstrap draw indexing. The manuscript-primary `PRIMARY` domains were therefore $\alpha$-invariant by construction; the threshold sweep evaluated only the operationally restricted domains.

For forward slopes, one largest contiguous Step-9 abundance support valid for `TOPLUS`, `TT`, and `TF` at every threshold was used for cross-threshold comparison. This prevented threshold-dependent loss of low-abundance bins from masquerading as a slope change. Native-support slopes were retained only as descriptive diagnostics.

For temporal sensitivity, one cross-threshold common operational core was defined as the largest contiguous subset of the frozen Step-13/14 core with valid `PRIMARY`, `NON_FF`, and `TT` support for every threshold and every lag. Equal-bin temporal covariance profiles and signed slopes were calculated on this same core for all thresholds.

Because subject universes, seeds, and bootstrap draw IDs were identical across $\alpha$, threshold-specific curves and slopes were compared with paired bootstrap differences against $\alpha = 0.05$. A forward bootstrap slope was retained only when all bins in the cross-threshold common forward support were finite. A temporal bootstrap slope was retained only when the entire cross-threshold common operational core satisfied the support requirements at every lag.

#### 9. Interpretation boundaries and software implementation

The analyses were implemented in `14-interval_position_structure.py`, `15-detectability_boundary_sensitivity.py`, and `16-analyze_observation_threshold_robustness.py`, which respectively addressed interval composition, fixed-threshold operational-domain sensitivity, and numerical-threshold robustness. None refitted latent abundance, subtracted the pseudo-longitudinal Step-12 technical reference, interpreted operational T/F as biological presence/absence, or replaced the manuscript-primary Step-9, Step-11, or Step-13 estimands.

### References

Benjamini Y, Hochberg Y. Controlling the false discovery rate: a practical and powerful approach to multiple testing. *Journal of the Royal Statistical Society: Series B (Methodological)*. 1995;57(1):289–300. doi:10.1111/j.2517-6161.1995.tb02031.x.

Deen M, de Rooij M. ClusterBootstrap: An R package for the analysis of hierarchical data using generalized linear models with the cluster bootstrap. *Behavior Research Methods*. 2020;52(2):572–590. doi:10.3758/s13428-019-01252-y.

Phipson B, Smyth GK. Permutation P-values should never be zero: calculating exact P-values when permutations are randomly drawn. *Statistical Applications in Genetics and Molecular Biology*. 2010;9:Article 39. doi:10.2202/1544-6115.1585.

Thabane L, Mbuagbaw L, Zhang S, et al. A tutorial on sensitivity analyses in clinical trials: the what, why, when and how. *BMC Medical Research Methodology*. 2013;13:92. doi:10.1186/1471-2288-13-92.

Winkler AM, Webster MA, Vidaurre D, Nichols TE, Smith SM. Multi-level block permutation. *NeuroImage*. 2015;123:253–268. doi:10.1016/j.neuroimage.2015.05.092.
