## Extended_Methods_2 for "ClonoDynamics enables replicate-resolved inference of longitudinal T cell receptor repertoire dynamics"

#### Support-conditioned synthetic positive controls: mathematical model and computational reproduction

This supplement describes the construction and analysis of semi-synthetic positive controls for temporal accumulation in TCRβ repertoire dynamics. Part I defines the experiment, the quantities compared and their uncertainty. Part II provides a step-by-step reproduction protocol using the corrected repertoire-count tables together with the finalized empirical ClonoDynamics outputs required to freeze the validation calibration. Appendix A documents the current repository entry points, code layout and archived execution records. The likelihood and fitting procedure of the production state-inference model are described in the main Methods; they are not replaced by a simulation-specific inference model.

### Part I. Mathematical model and recovery analysis

#### 1. Objective and empirical conditioning

We assessed whether the longitudinal analysis could recover a prescribed increase in clonotype displacement variability with elapsed time under the observation structure of the empirical study. The controls were semi-synthetic: underlying frequency trajectories and read counts were generated, whereas the sampling design, replicate-specific sequencing depths and clonotype detection patterns were inherited from the empirical data. Consequently, the experiment evaluated recovery conditional on observed support, rather than jointly simulating biological abundance and the process by which clonotypes become detectable. This design follows the general use of simulation studies to evaluate recovery against known ground truth while preserving application-relevant data features (Burton et al., 2006; Morris et al., 2019; Sandve and Greiff, 2022).

The empirical design comprised 10 participants sampled at six nominal weekly time points, with two technical replicates per available visit. The 114 available repertoires formed 57 complete replicate pairs. Eligibility for downstream analysis was recorded separately from physical sample availability: 56 pairs were eligible after state inference, defining 130 within-subject sampling-time intervals. A pair excluded because of unsuccessful inference was not interpreted as a biologically absent visit. Its raw repertoire files remained available for construction of the subject-level reference, but it did not contribute synthetic count repertoires to the analysis-eligible design.

Let $s$ index subjects, $i$ clonotypes, $t \in \{1, \dots, 6\}$ nominal sampling times and $r \in \{A, B\}$ technical replicates. All logarithms are natural. We distinguish empirical counts $C^{\mathrm{emp}}_{sir}(t)$, simulated frequencies $f^{(d)}_{si}(t)$ in scenario $d$, and synthetic counts $C^{(d)}_{sir}(t)$. Superscript $d = 0$ denotes the non-accumulating plateau control.

#### 2. Subject-level reference repertoires and frozen calibration

For each subject, the reference universe $\mathcal{I}_s$ was the union of amino-acid CDR3 clonotypes observed in all available empirical repertoire files. Repeated `aaSeqCDR3` rows were aggregated within each repertoire by summing their read counts. No never-observed clonotypes were introduced, and temporal order was not used to construct reference frequencies.

If $\mathcal{R}_s$ denotes the available empirical repertoires of subject $s$, $m_s = |\mathcal{R}_s|$ and $N_{sr}(t)$ their sequencing depths, the reference was

$$\bar{f}_{si} = \frac{1}{m_s} \sum_{(t,r)\in\mathcal{R}_s} \frac{C^{\mathrm{emp}}_{sir}(t)}{N_{sr}(t)}, \qquad f^{\mathrm{ref}}_{si} = \frac{\bar{f}_{si}}{\sum_{j\in\mathcal{I}_s} \bar{f}_{sj}}.$$

An unobserved clonotype contributed zero to a repertoire-specific frequency. This construction gave equal weight to repertoires, rather than weighting them by sequencing depth. The pooled-count reference exported by the calibration builder was a separate quantity and was not the reference used for this dose series.

The frozen calibration recorded the analysis-eligibility mask, replicate depths, clonotype-positive sets and pair-specific dispersion parameters, together with the empirical abundance-bin grid and temporal-analysis core. The fitted Step-2 parameter `k` was retained as $\kappa_{st}$, the Gamma shape parameter used in synthetic count allocation. Empirical temporal estimates were retained as documentary fields only; they were not targets for selecting the cumulative signal or adjusting its recovery.

#### 3. Simulated frequency trajectories

For each subject and clonotype, scenario-specific log weights were generated as

$$\ell_{si}^{(d)}(t) = \log f_{si}^{\text{ref}} + \sigma_{\text{fast}} z_{si}(t) + \sqrt{q_d}\, W_{si}(t),$$

where $z_{si}(t)$ are independent standard-normal visit-specific shocks before frequency normalization. The unit random walk started at the first nominal time with $W_{si}(1) = 0$ and evolved according to

$$W_{si}(t_j) = W_{si}(t_{j-1}) + \sqrt{t_j - t_{j-1}}\, \varepsilon_{si}(t_j), \qquad \varepsilon_{si}(t_j) \sim \mathcal{N}(0, 1).$$

The fast shocks and random-walk increments were generated separately. The fast component was fixed at $\sigma_{\text{fast}} = 0.20$ in every scenario; $q_d$, corresponding to `q_bio`, controlled the variance of cumulative log-weight increments per week. Frequencies were normalized independently at each subject and time point:

$$f_{si}^{(d)}(t) = \frac{\exp\{\ell_{si}^{(d)}(t)\}}{\sum_{j \in \mathcal{J}_s} \exp\{\ell_{sj}^{(d)}(t)\}}.$$

The implementation used a stable log-softmax calculation. Trajectories were generated at all six nominal times, including times that were not eligible for count generation. Only eligible visits were subsequently used to create observed synthetic repertoires.

##### 3.1 Common random numbers

The master simulation seed was `20260918`. Fast shocks and unit random-walk increments were determined by subject and time using NumPy `SeedSequence` inputs `[master, subject, 117031, 501, time]` and `[master, subject, 117031, 502, time]`, respectively. The same random fields and reference-clonotype ordering were retained across scenarios, applying the common-random-numbers principle to reduce irrelevant Monte Carlo variation in paired scenario contrasts (Stout and Goldie, 2008). Changing $q_d$ therefore rescaled the same unit random-walk path rather than drawing a new path; the plateau and positive-dose scenarios shared identical fast shocks.

Count-generation streams were initialized separately using `[master, subject, 910247, 500]`. Within each subject, eligible visits were processed in time order and the two replicates sequentially. Separate Gamma and multinomial variates were used for the two replicates. Reusing a count-stream initialization across doses does not imply that every subsequent count-generation variate is matched across scenarios; the explicitly matched random fields are the fast shocks and unit random-walk paths.

#### 4. Oracle definition and dose calibration

The known simulated frequencies provided an oracle reference computed without using counts or downstream recovery estimates. For every analysis-eligible interval $(t_0, t_1)$, the true displacement was

$$\Delta x_{si}^{\text{true},(d)}(t_0, t_1) = \log f_{si}^{(d)}(t_1) - \log f_{si}^{(d)}(t_0).$$

Its population variance across the full subject reference universe, of size $J_s = |\mathcal{J}_s|$, was

$$v_s^{(d)}(t_0, t_1) = \frac{1}{J_s} \sum_{i \in \mathcal{J}_s} \left[ \Delta x_{si}^{\text{true},(d)}(t_0, t_1) - \overline{\Delta x}_s^{\text{true},(d)}(t_0, t_1) \right]^2.$$

This corresponds to `ddof=0`. Let $\mathcal{P}_s(h)$ be the eligible intervals of subject $s$ with lag $h = t_1 - t_0$, and let $\mathcal{S}_h$ contain subjects with at least one such interval. The oracle profile was

$$V_d(h) = \frac{1}{|\mathcal{S}_h|} \sum_{s \in \mathcal{S}_h} \frac{1}{|\mathcal{P}_s(h)|} \sum_{(t_0,t_1) \in \mathcal{P}_s(h)} v_s^{(d)}(t_0, t_1).$$

Thus interval variances were first averaged within subject and lag, then averaged equally across subjects available at that lag. The oracle used analysis-eligible visits but all reference clonotypes; it was not restricted to `common4` transitions, inferred-abundance bins or the nine-subject complete-case recovery cohort.

Positive doses targeted the relative increase

$$R_d = \frac{V_d(5) - V_d(1)}{V_d(1)}$$

at values of 0.25, 0.50 and 1.00. For each target, the calibrator bracketed $q_d$, expanded the upper bound when required, and applied up to 22 bisection iterations with an absolute target tolerance of $5 \times 10^{-4}$. The candidate with the smallest absolute target error was retained. Common random fields were fixed throughout this search. Synthetic counts, inferred states, empirical cross-replicate covariance and recovered temporal slopes were not used to select the dose.

For the zero-accumulation scenario, $q_0$ was fixed at zero without tuning its finite-sample oracle profile to have an exactly zero slope or exactly zero realized $R$. This plateau retained the non-accumulating fast component; it was distinct from the generator's strict `null` preset, which sets both components to zero.

| Scenario | Target $R$ | $\sigma_{\text{fast}}$ | Calibrated $q_d$ |
|---|---|---|---|
| `R0p00` | 0.00 | 0.20 | 0 |
| `R0p25` | 0.25 | 0.20 | 0.005333333333333334 |
| `R0p50` | 0.50 | 0.20 | 0.01142857142857143 |
| `R1p00` | 1.00 | 0.20 | 0.026666666666666672 |

These are the parameter values recorded for the reported series. A new reproduction reads the values generated in `oracle/01_scenario_table.csv`, rather than inserting rounded values from this table. Target and realized $R$, oracle profiles and the search trace are exported separately. The oracle temporal slope, $\beta_d^{\text{oracle}}$, was an unweighted least-squares linear fit of $V_d(h)$ across $h = 1, \ldots, 5$.

#### 5. Read-count generation conditional on empirical support

For each eligible subject, visit and replicate, let

$$A_{str} = \{i : C_{sir}^{\text{emp}}(t) > 0\}$$

be the frozen empirical-positive clonotype set and let $K_{str} = |A_{str}|$. Given simulated frequencies, independent Gamma factors were drawn for clonotypes in this set:

$$G_{sir}^{(d)}(t) \sim \text{Gamma}\left(\kappa_{st},\ \text{scale} = 1/\kappa_{st}\right), \qquad i \in A_{str}.$$

Frequency-weighted allocation probabilities were then defined by

$$p_{sir}^{(d)}(t) = \frac{f_{si}^{(d)}(t) G_{sir}^{(d)}(t)}{\sum_{j \in A_{str}} f_{sj}^{(d)}(t) G_{sjr}^{(d)}(t)}.$$

Each empirical-positive clonotype received one mandatory read. The remaining reads were allocated as

$$\left(M_{sir}^{(d)}(t)\right)_{i\in A_{str}} \sim \text{Multinomial}\left(N_{sr}(t)-K_{str}, \left(p_{sir}^{(d)}(t)\right)_{i\in A_{str}}\right),$$

$$C_{sir}^{(d)}(t) = \begin{cases} 1+M_{sir}^{(d)}(t), & i \in A_{str}, \\ 0, & i \notin A_{str}. \end{cases}$$

The generator required $0 < K_{str} \leq N_{sr}(t)$ and a finite positive $\kappa_{st}$. If the sum of Gamma-weighted frequencies was non-finite or non-positive, the numerical safeguard used the unperturbed simulated frequencies within the active set before normalization.

This construction preserved the exact sequencing depth, observed richness and clonotype-specific positivity mask of each empirical replicate. Synthetic repertoire data with known generating structure are widely used for immunoinformatics benchmarking, although the present generator was deliberately conditioned on empirical support rather than designed to simulate de novo receptor repertoires (Weber et al., 2020). It was not an unconditioned draw of independent negative-binomial counts. Original empirical count vectors were not reused as synthetic observations, although their frequencies, fitted dispersion and positive sets informed the calibration.

Each scenario comprised 112 synthetic repertoires from the 56 eligible replicate pairs. Count outputs were written to `repertoires/` with `aaSeqCDR3`, `readCount` and recalculated `readFraction` columns. Known log-frequency trajectories were written separately to `ground_truth/` and never supplied to production state inference.

##### 5.1 Support verification

For each subject and clonotype, let $k_{si}$ be the number of eligible visits positive in at least one replicate, and $b_{si}$ the number positive in both replicates. The generator checked the implied all-interval and four-measure support totals using

$$N_{\text{transitions}} = \sum_s \sum_i \binom{k_{si}}{2}, \qquad N_{\text{common4}} = \sum_s \sum_i \binom{b_{si}}{2}.$$

For the six-timepoint design, all visit pairs have lags within 1–5 weeks. The frozen totals were 2,822,922 transitions and 1,182,069 `common4` transitions. Replicate-positive, union-positive and both-positive state counts were also checked against the calibration. These generator-level checks were kept distinct from checks on the support actually retained after synthetic state inference and downstream analysis.

#### 6. Recovery through the longitudinal-analysis pipeline

Synthetic count repertoires were processed using the production state-inference, trajectory-assembly and transition-assembly workflow. State inference estimated abundances from the synthetic technical-replicate counts; it did not read the oracle trajectories. The downstream transition table retained replicate-specific counts, depths and observed log-frequencies alongside the model-based state representation.

For positive counts, the observed log-frequency and displacement were

$$x_{sir}^{\text{obs},(d)}(t) = \log\left[\frac{C_{sir}^{(d)}(t)}{N_{sr}(t)}\right], \qquad \Delta x_{sir}^{\text{obs},(d)}(t_0,t_1) = x_{sir}^{\text{obs},(d)}(t_1) - x_{sir}^{\text{obs},(d)}(t_0).$$

Fluctuation recovery used `common4`: both replicates had to contain positive counts at both interval endpoints. The conditioning coordinate was the midpoint of the inferred endpoint log-frequencies,

$$x_{\text{mid},si}^{(d)}(t_0,t_1) = \frac{\hat{x}_{si}^{(d)}(t_0) + \hat{x}_{si}^{(d)}(t_1)}{2}.$$

The empirical 20-bin abundance grid was retained across scenarios. The reproduction configuration defines this grid with equal-width binning and the empirical 0.01 and 0.99 quantile limits; synthetic Step 11 reads the exported empirical edges instead of estimating new ones.

Within subject $s$, lag $h$ and bin $b$, let $\mathcal{T}_{shb}^{(d)}$ contain the eligible transitions and let $n_{shb}^{(d)}$ denote their count. Each transition is one clonotype–interval record. The subject-level cross-replicate covariance was

$$\widehat{C}_{shb}^{(d)} = \frac{1}{n_{shb}^{(d)} - 1} \sum_{u \in \mathcal{T}_{shb}^{(d)}} \left( \Delta x_{uA}^{(d)} - \overline{\Delta x}_{A,shb}^{(d)} \right) \left( \Delta x_{uB}^{(d)} - \overline{\Delta x}_{B,shb}^{(d)} \right).$$

Both means were computed on the same transition set. Covariance was calculated across transitions pooled within subject, lag and bin, rather than by averaging interval-specific covariances. It remained signed and was not truncated at zero. No pseudo-longitudinal covariance was subtracted, and operational TT/TF/FT/FF classifications did not define the primary recovery population.

#### 7. Fixed abundance core and temporal summaries

The temporal core was selected from the empirical grid before analysis of the synthetic scenarios. At each lag, a subject–lag–bin cell was valid when its cross-replicate covariance was finite and at least two transitions contributed. A bin qualified if it contained valid cells for at least

$$\max\{2, \lceil 0.90\, n_{\text{available}}(h) \rceil\}$$

subjects at every requested lag. The largest contiguous qualifying bin run defined the core; ties were resolved in favour of the run starting at lower abundance. The complete-case cohort comprised subjects with a valid primary covariance in every selected bin at every lag.

The reported analysis used 17 bins, IDs 3–19, spanning $x_{\text{mid}} = -12.4129$ to $-8.5494$, and nine complete-case subjects: 1, 2, 3, 4, 5, 7, 8, 9 and 10. These same bins were supplied to synthetic Step 13. Complete-case subjects were nevertheless determined from each scenario's outputs and checked for agreement with the empirical cohort; they were not assumed to be unchanged merely because the detection masks were fixed. Membership of individual transitions in inferred-abundance bins was allowed to differ across doses.

The minimum of two transitions for a subject-level covariance cell is distinct from the Step-11 pooled-bin reporting threshold of 50 transitions. The latter was not imposed as a minimum of 50 transitions in every subject–lag–bin cell.

Let $\mathcal{B}$ be the fixed core and $\mathcal{S}$ the complete-case subject set. The primary temporal summary gave equal weight to bins within subject, then equal weight to subjects:

$$m_s^{(d)}(h) = \frac{1}{|\mathcal{B}|} \sum_{b \in \mathcal{B}} \widehat{C}_{shb}^{(d)}, \qquad M_d(h) = \frac{1}{|\mathcal{S}|} \sum_{s \in \mathcal{S}} m_s^{(d)}(h).$$

An unweighted least-squares fit across the five lag values described the temporal profile as

$$M_d(h) = a_d + \beta_d^{\text{recovered}}(h-1).$$

The fitted signed slope, in squared natural-log-frequency units per week, was the primary recovery descriptor. The displacement covariance was not divided by lag before fitting. A positive effect was described as resolved when the lower endpoint of its 95% subject-bootstrap interval exceeded zero.

Step 13 additionally exported a sensitivity estimate using bin weights $n_{shb}^{(d)} - 1$ within subject while retaining equal subject weights. It also evaluated mean within-replicate displacement variance and replicate-specific excess on the same fixed cohort. Subject slopes, sign-flip tests, binwise slopes and finite-lag model descriptors were retained as ancillary outputs; the dose-response summaries selected `equal_bin_equal_subject` and `cross_cov` explicitly.

#### 8. Joint subject bootstrap and incremental recovery

Uncertainty was estimated using 2,000 subject-bootstrap draws with seed 123, with the biological subject retained as the cluster-level resampling unit (Deen and de Rooij, 2020). Subjects were ordered by identifier. For each bootstrap draw, nine subject indices were sampled with replacement and the same draw was propagated across all lags, metrics and estimators. Scenario analyses used the same ordered subject set and seed. Their matching was checked by reconstructing the subject draws and the primary bootstrap curves and slopes from the subject-level outputs.

For draw $k$, with resampled subjects $s_1^{*(k)}, \dots, s_9^{*(k)}$, the primary curve was

$$M_d^{*(k)}(h) = \frac{1}{9}\sum_{j=1}^{9} m_{s_j^{*(k)}}^{(d)}(h).$$

The temporal slope was refitted to each resampled curve. The 2.5th and 97.5th percentiles supplied the interval endpoints. These resamples did not regenerate frequencies or counts and did not refit the state-inference model.

Absolute recovered slopes were retained for all four scenarios. Secondary recovery summaries compared each positive dose with the plateau:

$$\Delta\beta_d^{\text{recovered}} = \beta_d^{\text{recovered}} - \beta_0^{\text{recovered}}, \qquad \Delta\beta_d^{\text{oracle}} = \beta_d^{\text{oracle}} - \beta_0^{\text{oracle}}.$$

Paired bootstrap differences were formed within matching draws,

$$\Delta\beta_d^{*(k)} = \beta_d^{\text{recovered},*(k)} - \beta_0^{\text{recovered},*(k)}.$$

Intervals for these contrasts were the corresponding percentiles of the paired difference distribution, not differences between marginal interval endpoints. The summarizer matches records by bootstrap ID; the separate verification step establishes that these IDs identify the same subject resamples across scenarios.

For nonzero oracle increments, incremental recovery was

$$\rho_d = \frac{\Delta\beta_d^{\text{recovered}}}{\Delta\beta_d^{\text{oracle}}}.$$

Its bootstrap distribution divided each paired recovered increment by the fixed oracle increment. A descriptive through-origin fit across positive doses gave

$$\hat{\lambda} = \frac{\sum_{d>0} \Delta\beta_d^{\text{oracle}} \Delta\beta_d^{\text{recovered}}}{\sum_{d>0} \left(\Delta\beta_d^{\text{oracle}}\right)^2}.$$

An unconstrained descriptive linear fit of absolute recovered against oracle slopes across all four scenarios was also exported. Neither fit was used to recalibrate the doses or correct the empirical results. The oracle and recovered quantities have different support and weighting definitions; their ratio is therefore a conditional recovery benchmark, not a universal measure of estimator accuracy.

This experiment comprises one coupled realization per dose under the specified reference, random fields and count-generation procedure. Subject-bootstrap draws quantify resampling uncertainty within these realizations. They do not constitute independent simulations, establish a power curve or define a general detection limit for temporal accumulation. The validation does not assess recovery outside the inherited detection-support structure.

### Part II. Computational workflow and execution protocol

#### 9. Scope and software requirements

The positive-control workflow is implemented as a dedicated validation block within the current ClonoDynamics repository. It does not define a separate inference model and does not replace the production longitudinal analysis. Instead, it freezes the finalized empirical calibration, calibrates the oracle dose series, generates support-conditioned synthetic repertoires, and then reuses the same production Steps 2, 4, 5, 11 and 13 used for the empirical analysis.

The current repository entry point is

```
python3 code/orchestrator_clonodynamics.py
```

and the positive-control implementation is located under

```
code/03_validation/positive_controls/
```

with the production longitudinal dependencies retained under `code/01_core/` and `code/03_validation/`.

The reported production environment used Python 3.9.22. The frozen release records the principal package versions and MiXCR version in `requirements-freeze.txt` and `run_manifest.yaml`. Reproduction of an archived analysis should use the tagged software release and its recorded environment rather than an arbitrary later version of the repository, consistent with general recommendations for reproducible computational research (Sandve et al., 2013).

The positive-control workflow does **not** rerun or redefine the finalized empirical analysis as part of the validation block. It requires the corrected empirical repertoire tables and the completed empirical outputs needed to freeze the calibration. In particular, the calibration reads the empirical Step-1 repertoire characteristics, Step-2 fitted pair parameters and QC, the Step-5 transition-assembly report, the Step-11 abundance grid/support outputs and the Step-13 temporal core and complete-case cohort. The empirical results therefore remain the upstream reference, while synthetic scenarios are analyzed independently through the production recovery chain.

#### 10. Input files, repository layout and frozen prerequisites

##### 10.1 Corrected empirical repertoire tables

The corrected empirical input consists of 114 clonotype-level repertoire tables arranged as 57 complete technical-replicate pairs. Each file corresponds to one subject–time–replicate combination and contains the clonotype-level fields used by ClonoDynamics:

```
aaSeqCDR3
readCount
readFraction
```

Rows sharing the same `aaSeqCDR3` have already been collapsed by summing `readCount`, and `readFraction` has been recalculated from the resulting repertoire-specific total. The naming convention is

```
subject_time-replica.tsv
```

for example:

```
1_1-1.tsv
1_1-2.tsv
1_2-1.tsv
1_2-2.tsv
10_6-1.tsv
10_6-2.tsv
```

The archived reproducibility dataset contains the exact processed repertoire inputs supplied to the production pipeline. Missing visits are represented by absent files rather than by artificial placeholder repertoires.

##### 10.2 Current code layout

The relevant public repository structure is

```
clonodynamics-pipeline/
   code/
       orchestrator_clonodynamics.py
       01_core/
           1-repertoire_characterization.py
           2-multirepresentation_clonotype_state_inference.py
           noiseK_latent.py
           4-multirepresentation_trajectory_assembly.py
           5-longitudinal_transition_assembly.py
       03_validation/
           11-cross_replicate_fluctuation_dynamics.py
           13-temporal_fluctuation_scaling.py
           positive_controls/
               01_freeze_empirical_calibration.py
               02_calibrate_oracle_doses.py
               03_generate_support_conditioned_repertoires.py
               04_summarize_temporal_recovery.py
               05_plot_synthetic_validation.py
               verify_workflow.py
   requirements-freeze.txt
   run_manifest.yaml
```

The master orchestrator contains the current validated orchestration logic. The scientific scripts remain separately executable for inspection, testing and direct reproduction. Script identity is tracked through the repository release and, where generated by the workflow, script hashes and run signatures.

##### 10.3 Required empirical outputs

The positive-control block requires a completed empirical ClonoDynamics analysis. The essential upstream products are:

```
1-repertoire_characterization/
2-multirepresentation_clonotype_state_inference/
5-longitudinal_transition_assembly/
11-cross_replicate_fluctuation_dynamics/
13-temporal_fluctuation_scaling/
```

The calibration builder uses these outputs to freeze:

- the raw and analysis-eligible sampling design;
- replicate-specific sequencing depths and positive clonotype sets;
- pair-specific Step-2 dispersion parameters;
- the empirical Step-11 abundance-bin grid and support structure;
- the Step-13 abundance core and complete-case subject cohort.

The validation block therefore inherits the empirical analysis contract instead of reconstructing or re-optimizing it.

#### 11. Current execution workflow

##### 11.1 Launch the positive-control block

Run the master interface from the repository root:

```
python3 code/orchestrator_clonodynamics.py
```

Select:

```
Analysis
→ positive_controls
```

The positive-control block requests the corrected longitudinal input dataset, the completed empirical/core and validation outputs, and a destination directory for positive-control results. The orchestrator records the selected paths and execution configuration and routes the workflow through the validated positive-control components.

For exact reconstruction of the reported execution, use the archived tagged software release and the run-specific configuration/provenance files rather than relying on interactive defaults from a later repository state.

##### 11.2 Freeze the empirical calibration

`01_freeze_empirical_calibration.py` reads the corrected empirical repertoire tables and the finalized empirical outputs. It creates the frozen calibration directory, which records the raw and analysis-eligible sampling designs, Step-2 pair parameters, observation-support targets, subject-union references, empirical bin edges, temporal-core definition and source audits.

The reported calibration checks the cohort-specific design against the observed study structure. A mismatch is treated as a provenance or compatibility failure; the workflow should not be altered simply to force agreement with the archived values.

The principal calibration products are stored under

```
positive_controls/calibration/
```

and include the frozen sampling designs, Step-2 parameter table, observation-structure targets, Step-11 bin edges, Step-13 core definition, complete-case subjects, source-repertoire audit and validation contract.

##### 11.3 Calibrate the oracle dose series

`02_calibrate_oracle_doses.py` reads the frozen calibration and constructs the four reported scenarios:

```
R0p00
R0p25
R0p50
R1p00
```

using the parameterization defined in Part I. The reported series uses

```
sigma_fast = 0.20
master seed = 20260918
target R = 0, 0.25, 0.50, 1.00
```

with the calibrated `q_bio` values written to the oracle scenario table. Later workflow stages read the generated scenario table directly rather than inserting rounded parameter values manually.

Oracle products are written under

```
positive_controls/oracle/
```

and include the scenario table, oracle temporal profiles, calibration/search records and manifest.

##### 11.4 Generate the synthetic repertoire scenarios

For each oracle scenario, `03_generate_support_conditioned_repertoires.py` generates synthetic repertoire-count tables using the support-conditioned count model described in Part I.

Each scenario contains:

```
repertoires/
ground_truth/
00_manifest.json
01_generated_sampling_summary.csv
02_oracle_pair_summary.csv
03_support_audit_by_subject.csv
```

Only `repertoires/` is supplied to the production inference chain. `ground_truth/` remains separate and is used only for oracle evaluation and validation.

The generator checks the declared support contract, including sequencing depth, richness and replicate-specific positivity structure. A support mismatch stops the workflow rather than being repaired by changing the generated data after the fact.

##### 11.5 Analyse each synthetic scenario with the production pipeline

Each synthetic repertoire set is processed through the production recovery chain:

```
Step 2 → Step 4 → Step 5 → Step 11 → Step 13
```

using the same scientific implementations as the empirical analysis.

For the positive-control recovery analysis:

- Step 2 infers states from the synthetic technical-replicate counts;
- Step 4 assembles multirepresentation trajectories;
- Step 5 constructs the finite-time transition table;
- Step 11 estimates signed cross-replicate covariance on the frozen empirical abundance grid;
- Step 13 estimates temporal scaling on the frozen empirical core.

Synthetic Step 11 receives the empirical bin edges frozen during calibration. Synthetic Step 13 receives the empirical core-bin definition. These fixed supports prevent scenario-specific abundance-grid selection from becoming part of the dose response.

The scenario-specific analysis directories retain their production-style step outputs. Within the positive-control workspace, the Step-2 scenario output directory may retain the internal name

```
2-clonotype_state_inference/
```

for compatibility with the validation engine; this is an execution-directory name and does not replace the public Step-2 script name `2-multirepresentation_clonotype_state_inference.py`.

##### 11.6 Verify support, reconstruction and bootstrap pairing

`verify_workflow.py` checks the compatibility of the generated scenarios and their recovered analyses with the frozen empirical validation contract.

The verification includes checks of:

- scenario parameters and seeds against the oracle table;
- analysed replicate-pair eligibility;
- transition and `common4` support;
- Step-11 abundance edges;
- Step-13 core bins and complete-case subjects;
- the Step-11/Step-13 analysis link;
- reconstruction of the primary subject-bootstrap curves and slopes;
- matching of bootstrap draw identities across scenarios before paired contrasts are calculated.

The verification products are stored under

```
positive_controls/verification/
```

and include the check table, verification summary, reconstructed subject draws and verified paired increments.

Verification tests consistency of the declared computational design. It does not test whether a recovered slope has a desired sign and does not use the oracle outcome to modify the production estimators.

##### 11.7 Produce numerical recovery summaries

`04_summarize_temporal_recovery.py` reads the oracle products and scenario-specific Step-13 outputs and extracts the primary `equal_bin_equal_subject` / `cross_cov` recovery estimates.

The numerical summaries are written under

```
positive_controls/summary/
```

and retain:

- absolute recovered temporal profiles and slopes;
- positive-dose-minus-plateau paired increments;
- oracle increments;
- incremental recovery ratios;
- descriptive oracle-versus-recovered fits;
- validation metadata.

Paired contrast intervals are calculated from matching bootstrap draw IDs, not by subtracting marginal confidence-interval endpoints.

##### 11.8 Plotting and manuscript source data

The scientific recovery calculations are complete before plotting. `05_plot_synthetic_validation.py` and the read-only publication plotting layer consume the finalized numerical outputs without regenerating synthetic data, refitting state inference or recomputing temporal slopes.

The manuscript-facing numerical summaries support Supplementary Data 9. Plot-specific file names may evolve with the publication layout, but such changes do not alter the underlying recovery calculations.

#### 12. Execution records, reruns and reproducibility boundaries

The current positive-control orchestration records run configuration and provenance at the workflow level. The archived production workspace includes, where applicable:

```
00_positive_controls_orchestrator_config.json
00_positive_controls_step_manifest.csv
source_code_inventory.json
commands.tsv
calibration/
oracle/
verification/
summary/
```

Large scenario repertoires and scenario-specific computational intermediates are deterministic products of the frozen inputs, scenario table, random seeds and software release. They need not be duplicated in the compact reproducibility archive when they can be regenerated from those frozen components.

The reported positive-control experiment comprises one coupled realization per dose under the specified empirical reference, random fields and count-generation procedure. Subject-bootstrap draws quantify resampling uncertainty within these realizations; they are not independent synthetic experiments and do not define a general power curve.

An interrupted or partially completed execution should be handled according to the current orchestrator's compatibility and existing-output checks. Completed compatible outputs may be reused, whereas incompatible outputs should not be silently overwritten. The archived configuration files, commands, manifests, script hashes/signatures, input checksums and scientific outputs constitute the provenance record of the reported analysis.

### Appendix A. Current command and provenance reference

This appendix summarizes the current public entry points. Exact effective commands for the reported run are retained in the archived `commands.tsv` and configuration/manifests generated by the positive-control workflow.

#### A.1 Master entry point

From the repository root:

```
python3 code/orchestrator_clonodynamics.py
```

Select:

```
Analysis
→ positive_controls
```

The master orchestrator then requests the empirical processed input directory, completed empirical analysis outputs and the positive-control destination directory.

#### A.2 Scientific programs used by the validation block

The positive-control block invokes or depends on the following production and validation programs:

```
code/01_core/2-multirepresentation_clonotype_state_inference.py
code/01_core/4-multirepresentation_trajectory_assembly.py
code/01_core/5-longitudinal_transition_assembly.py
code/03_validation/11-cross_replicate_fluctuation_dynamics.py
code/03_validation/13-temporal_fluctuation_scaling.py

code/03_validation/positive_controls/01_freeze_empirical_calibration.py
code/03_validation/positive_controls/02_calibrate_oracle_doses.py
code/03_validation/positive_controls/03_generate_support_conditioned_repertoires.py
code/03_validation/positive_controls/04_summarize_temporal_recovery.py
code/03_validation/positive_controls/05_plot_synthetic_validation.py
code/03_validation/positive_controls/verify_workflow.py
```

`code/01_core/noiseK_latent.py` is imported by Step 2 as the latent-state inference engine.

#### A.3 Frozen analysis constants used in the reported validation

The principal reported settings are:

```
reference operational alpha = 0.05
Step-11 lags = 1, 2, 3, 4, 5 weeks
Step-11 abundance bins = 20
Step-11 empirical abundance limits = 0.01-0.99 quantiles
Step-11 minimum pooled bin support = 50 transitions
subject bootstrap draws = 2000
bootstrap seed = 123

synthetic master seed = 20260918
sigma_fast = 0.20
oracle targets R = 0, 0.25, 0.50, 1.00
```

```
oracle calibration iterations = 22
oracle target tolerance = 0.0005
```

The calibrated `q_bio` values are read from `oracle/01_scenario_table.csv`. The Step-11 bin edges and Step-13 core bins are read from the frozen empirical calibration rather than hard-coded from a figure or from rounded manuscript values.

##### A.4 Archived reproducibility records

The compact ClonoDynamics reproducibility dataset retains the empirical calibration, oracle definitions, verification outputs, numerical summaries and relevant run provenance. The full synthetic scenario directories are intentionally omitted from the compact archive because they can be regenerated from:

1. the corrected processed empirical repertoires;
2. the frozen empirical analysis outputs and calibration;
3. the oracle scenario table;
4. the recorded random seeds;
5. the tagged ClonoDynamics software release.

The manuscript-facing numerical summaries are collected in Supplementary Data 9. Complete workflow provenance remains in the archived machine-readable configuration, manifest and checksum files.

#### References


Burton A, Altman DG, Royston P, Holder RL. The design of simulation studies in medical statistics. *Statistics in Medicine*. 2006;25(24):4279–4292. doi:10.1002/sim.2673.

Deen M, de Rooij M. ClusterBootstrap: An R package for the analysis of hierarchical data using generalized linear models with the cluster bootstrap. *Behavior Research Methods*. 2020;52(2):572–590. doi:10.3758/s13428-019-01252-y.

Morris TP, White IR, Crowther MJ. Using simulation studies to evaluate statistical methods. *Statistics in Medicine*. 2019;38(11):2074–2102. doi:10.1002/sim.8086.

Sandve GK, Greiff V. Access to ground truth at unconstrained size makes simulated data as indispensable as experimental data for bioinformatics methods development and benchmarking. *Bioinformatics*. 2022;38(21):4994–4996. doi:10.1093/bioinformatics/btac612.

Sandve GK, Nekrutenko A, Taylor J, Hovig E. Ten simple rules for reproducible computational research. *PLoS Computational Biology*. 2013;9(10):e1003285. doi:10.1371/journal.pcbi.1003285.

Stout NK, Goldie SJ. Keeping the noise down: common random numbers for disease simulation modeling. *Health Care Management Science*. 2008;11(4):399–406. doi:10.1007/s10729-008-9067-6.

Weber CR, Akbar R, Yermanos A, et al. immuneSIM: tunable multi-feature simulation of B- and T-cell receptor repertoires for immunoinformatics benchmarking. *Bioinformatics*. 2020;36(11):3594–3596. doi:10.1093/bioinformatics/btaa158.
