## Extended_Methods_1 for "ClonoDynamics enables replicate-resolved inference of longitudinal T cell receptor repertoire dynamics"

### Scientific methods: primary ClonoDynamics analysis

**Scope:** repertoire characterization, replicate-resolved state inference, longitudinal transition construction, pseudo-longitudinal technical calibration, genuine forward dynamics, cross-replicate fluctuation dynamics, and temporal scaling (Steps 1–13).

This document is the **living GitHub methods documentation** for the primary ClonoDynamics analysis. Its scientific content is aligned with the extended methods prepared for the associated manuscript, while repository paths and execution references follow the current public code layout. The manuscript version should remain frozen with the corresponding paper release/tag.

For routine execution, use the unified launcher:

```
python3 code/orchestrator_clonodynamics.py
```

Current code mapping:

- Steps 1–6: `code/01_core/`
- pseudo-reference Steps 7–8: `code/02_pseudo_reference/`
- validation Steps 9–13: `code/03_validation/`
- publication plotting: `code/05_plotting/`

Back to the ClonoDynamics README

#### Table of contents

1. (i) Repertoire characterization and latent-state inference from paired technical replicates
    - Observed repertoire preprocessing and heavy-tail characterization
    - Pairing of technical replicates and replicate-resolved observed measurements
    - Pair-specific Negative-Binomial latent model
    - Joint and single-replicate latent posteriors
    - Realized detection and posterior-predictive detectability
    - Operational observability and inclusion policy
    - Code implementation
2. (ii) Assembly and characterization of the longitudinal transition dataset
    - Standardized multirepresentation trajectory assembly
    - Generic finite-time transition assembly
    - Replicate-resolved observed transition layer and estimand support
    - Operational endpoint annotation within the unrestricted transition universe
    - Full-posterior endpoint uncertainty propagation
    - Transition-universe and estimand-support characterization
3. (iii) Pseudo-longitudinal technical calibration and definition of the primary estimators
    - Pseudo-longitudinal technical-reference ensemble and inferential unit
    - Mathematical coupling and rationale for replicate decoupling
    - Step-7 primary observed replicate-decoupled forward benchmark
    - Matched same-versus-cross coupling control
    - Secondary latent conditioning-geometry diagnostic
    - Step-8 replicate-consistent fluctuation calibration
    - Randomization summaries and final estimator policy
4. (iv) Replicate-decoupled forward dynamics and comparison with the pseudo-longitudinal technical reference
    - Observed replicate-decoupled forward estimator
    - Abundance binning and fold combination
    - Subject-level uncertainty and curve descriptors
    - Matched same-measure control
    - Reference to the pseudo-longitudinal technical reference
5. (v) Replicate-resolved fluctuation dynamics, technical-reference comparison and temporal scaling
    - Cross-replicate fluctuation estimand

- Abundance grid, lag-resolved profiles and uncertainty
- Reference to the pseudo-longitudinal technical reference
- Fixed-core temporal scaling across 1–5 weeks
- Primary temporal estimator and sensitivity weighting
- Signed temporal slope, bootstrap inference and finite-lag models

#### (i) Repertoire characterization and latent-state inference from paired technical replicates

##### Observed repertoire preprocessing and heavy-tail characterization

Observed repertoire architecture was characterized using `1-repertoire_characterization.py`. Each technical replicate was analyzed independently. Read counts were converted to numeric values, non-finite and non-positive entries were discarded, and rows sharing the same CDR3 amino-acid sequence (`aaSeqCDR3`) were collapsed by summing their read counts. The input `readFraction` field, when present, was not used. For a repertoire with clonotype counts $c_i$, sequencing depth was

$$D = \sum_i c_i,$$

and the observed relative frequency of clonotype $i$ was recalculated as

$$f_i = \frac{c_i}{D}.$$

The number of distinct amino-acid CDR3 sequences after collapsing defined observed repertoire richness.

The high-frequency tail was first characterized using a continuous Pareto model, following established heavy-tail fitting approaches and prior descriptions of heavy-tailed TCR clone-size distributions (Clauset et al., 2009; Desponds et al., 2016; Koch et al., 2018),

$$p(f \mid f_{\text{min}}, \gamma) = (\gamma - 1) f_{\text{min}}^{\gamma-1} f^{-\gamma}, \qquad f \geq f_{\text{min}},\ \gamma > 1.$$

For a fixed lower boundary $f_{\text{min}}$, the maximum-likelihood estimate of the tail exponent was

$$\hat{\gamma} = 1 + \frac{n_{\text{tail}}}{\sum_{i=1}^{n_{\text{tail}}} \log(f_i / f_{\text{min}})}.$$

Candidate values of $f_{\text{min}}$ were taken from the observed frequency values, with a minimum tail size of 50 clonotypes. The selected boundary minimized the Kolmogorov-Smirnov distance

$$D_{\text{KS}} = \sup_f \left| F_{\text{emp}}(f) - F_{\text{Pareto}}(f) \right|.$$

Because tail-model conclusions can depend on the lower boundary, an additional threshold-dependent comparison was performed at empirical frequency quantiles 0.70, 0.75, 0.80, 0.85, 0.90, and 0.95. At each threshold, the same tail observations were evaluated under both the continuous Pareto model and a lower-truncated log-normal model,

$$p_{\text{TLN}}(f) = \frac{\exp\left[-(\log f - \mu)^2/(2\sigma^2)\right]}{f\sigma\sqrt{2\pi}\left[1 - \Phi\left((\log f_{\text{min}} - \mu)/\sigma\right)\right]}, \qquad f \geq f_{\text{min}}.$$

For numerical optimization, the lower-truncated log-normal was fitted through the equivalent coordinate $y = \log(f/f_{\text{min}}) \geq 0$, with density proportional to

$$\exp(-\beta y - \psi y^2).$$

The boundary $\psi = 0$ corresponds to the continuous Pareto family. The implementation explicitly compared the optimized truncated-log-normal interior with this boundary. Because the truncated-log-normal model has one additional fitted parameter and contains the Pareto model on its boundary, raw log-likelihood differences were retained as numerical diagnostics rather than used as a formal likelihood-ratio test. Model preference was summarized primarily by the small-sample corrected Akaike information criterion,

$$\mathrm{AICc} = \mathrm{AIC} + \frac{2k(k+1)}{n-k-1},$$

with BIC retained as a sensitivity criterion.

This repertoire-level analysis was descriptive. Sequencing depth, repertoire richness, Pareto fit success, fitted exponent, KS distance, tail fraction, and Pareto-versus-truncated-log-normal preference were not used as upstream exclusion criteria. In particular, the Step-1 Pareto exponent was neither passed to nor used to constrain the latent model.

**Pairing of technical replicates and replicate-resolved observed measurements**

Latent-state inference and construction of the multirepresentation clonotype-state table were implemented in `2-multirepresentation_clonotype_state_inference.py`, which calls `noiseK_latent.py` for latent-frequency inference. Only complete subject-timepoint replicate pairs were fitted. Within each replicate, duplicate `aaSeqCDR3` entries were collapsed by summing read counts. Replicates were then outer-joined by CDR3 amino-acid sequence. Consequently, every clonotype detected in at least one replicate was represented, whereas a clonotype not detected in the other replicate received an observed count of zero. A $(0, 0)$ row was not generated because the table was defined on the union of detected clonotypes.

For replicate $r$, sequencing depth was

$$N_r = \sum_i c_{ir},$$

and replicate-resolved observed relative frequency was

$$f_{ir}^{\mathrm{obs}} = \frac{c_{ir}}{N_r}.$$

For positive counts,

$$x_{ir}^{\mathrm{obs}} = \log f_{ir}^{\mathrm{obs}}.$$

When $c_{ir} = 0$, the corresponding observed log-frequency was left undefined rather than assigned an arbitrary finite pseudocount value. These replicate-resolved measurements were preserved because their independence is required by the downstream replicate-decoupled forward and cross-replicate fluctuation estimators.

**Pair-specific Negative-Binomial latent model**

For clonotype $i$, the two technical-replicate counts were assumed conditionally independent given a shared latent relative frequency $f_i$:

$$c_{ir} \mid f_i, N_r, \kappa \sim \mathrm{NB}(\mu_{ir} = N_r f_i, \mathrm{size} = \kappa).$$

The implementation used the mean/size parameterization

$$E(C_{ir} \mid f_i) = N_r f_i,$$

$$\mathrm{Var}(C_{ir} \mid f_i) = N_r f_i + \frac{(N_r f_i)^2}{\kappa}.$$

Thus, $\kappa$ captures technical overdispersion beyond Poisson sampling, consistent with replicate-based repertoire measurement models in which true clonotype frequency is latent and read-count variability is explicitly overdispersed (Puelma Touzel et al., 2020). A zero count is a valid Negative-Binomial observation and contributes to the likelihood without implying $f_i = 0$.

Latent frequencies were evaluated on a logarithmically spaced grid $\{f_1, \ldots, f_G\}$ with $G = 500$ and $f_{\text{max}} = 1$. Unless otherwise specified, the lower limit was

$$f_{\text{min}} = \frac{1}{\max\left[(N_1 + N_2)/2, 1\right]}.$$

The prior implemented in `noiseK_latent.py` is a normalized discrete probability mass over this grid,

$$\pi_j(\gamma) = \frac{f_j^{-\gamma}}{\sum_{k=1}^{G} f_k^{-\gamma}}.$$

No grid-cell-width quadrature weights are applied. The implemented prior is therefore a discrete power-law-shaped prior on a log-spaced grid, rather than numerical quadrature of an exact continuous Pareto density. The Step-2 exponent $\gamma$ is independently estimated and is not inherited from the descriptive Step-1 tail fit.

For each complete replicate pair, $\gamma$ and $\kappa$ were estimated jointly by marginal likelihood. For clonotype $i$,

$$P(c_{i1}, c_{i2} \mid \gamma, \kappa) = \sum_{j=1}^{G} \pi_j(\gamma) P(c_{i1} \mid f_j, N_1, \kappa) P(c_{i2} \mid f_j, N_2, \kappa),$$

and the pair-level objective was

$$\ell(\gamma, \kappa) = \sum_i \log P(c_{i1}, c_{i2} \mid \gamma, \kappa).$$

Optimization was performed with L-BFGS-B using starting values $\gamma = 1.6$ and $\kappa = 50$ and numerical lower bounds of $10^{-6}$ for both parameters. Marginal likelihood calculations were evaluated in clonotype chunks, with a default chunk size of 20,000, to limit peak memory use without altering the fitted objective.

**Joint and single-replicate latent posteriors**

After pair-specific parameter estimation, the joint posterior for each clonotype was

$$P(f_j \mid c_{i1}, c_{i2}) = \frac{\pi_j P(c_{i1} \mid f_j, N_1, \hat{\kappa}) P(c_{i2} \mid f_j, N_2, \hat{\kappa})}{\sum_k \pi_k P(c_{i1} \mid f_k, N_1, \hat{\kappa}) P(c_{i2} \mid f_k, N_2, \hat{\kappa})}.$$

Posterior mean, median, mode, 2.5th and 97.5th percentiles were calculated for both relative frequency and log-frequency, together with the posterior standard deviation in log-frequency space and posterior entropy,

$$H_i = -\sum_j p_{ij} \log p_{ij}.$$

The paired posterior represents a model-based latent consensus state for one biological sampling time. Its posterior median log-frequency (`x_latent_median`) was retained as the canonical latent point coordinate when downstream analyses required a noise-aware conditioning representation. It was not interpreted as directly observed or as ground-truth clonotype abundance.

Using the same pair-specific $\hat{\gamma}$, $\hat{\kappa}$, and latent-frequency grid, separate single-replicate posteriors $P(f \mid c_{i1})$ and $P(f \mid c_{i2})$ were also reconstructed. Their posterior summaries were retained for replicate-agreement diagnostics and latent-representation sensitivity analyses. Because both posterior distributions are evaluated on the same discrete grid, their overlap was summarized by the Bhattacharyya coefficient

$$BC_i = \sum_j \sqrt{p_{i1,j} p_{i2,j}}.$$

These single-replicate latent summaries do not constitute the primary AB/BA forward measurements in the final pipeline; the primary forward estimator uses the independently observed replicate-resolved log-frequencies.

For backward compatibility and representation diagnostics, an auxiliary conditional-positive abundance summary was also retained. When both replicates were positive,

$$f_i^+ = \sqrt{f_{i1}^{\text{obs}} f_{i2}^{\text{obs}}},$$

whereas for a clonotype detected in only one replicate the observed frequency of that positive replicate was used. This auxiliary quantity is not the primary fluctuation displacement in the final analysis.

**Realized detection and posterior-predictive detectability**

Realized technical detection was recorded separately for each replicate as $c_{ir} > 0$. These indicators describe sequencing outcomes and were not interpreted as biological presence or absence.

Model-based posterior-predictive detectability was calculated under the fitted Negative-Binomial observation model. With the default detection threshold of one read, the conditional dropout probability in replicate $r$ is

$$q_r(f) = P(C_r = 0 \mid f, N_r, \kappa) = \left( \frac{\kappa}{\kappa + N_r f} \right)^{\kappa},$$

and therefore

$$P(C_r \geq 1 \mid f) = 1 - q_r(f).$$

Replicate-specific posterior-predictive detectabilities were obtained by integrating the corresponding conditional probabilities over the joint paired-replicate posterior. For the paired state, conditional independence of replicate observations given $f$ gives

$$p_{\text{drop,state}} = E_{\text{post}} \left[ q_1(f) q_2(f) \right],$$

and

$$p_{\text{detect,state}} = 1 - p_{\text{drop,state}}.$$

Because both replicate predictions are integrated over the same uncertain latent frequency, paired-state detectability is not generally identical to a product constructed from separately posterior-averaged replicate detectabilities. These quantities are model-based technical detectability measures and were not treated as independently calibrated probabilities of biological presence. Their use as model diagnostics follows the broader posterior-predictive model-checking framework (Gelman et al., 1996).

##### Operational observability and inclusion policy

Operational observability was defined separately from posterior-predictive detectability. For each clonotype, the fitted model provides the marginal log probability of the observed replicate count pair,

$$\log P_i = \log \left[ \sum_j \pi_j P(c_{i1} \mid f_j, N_1, \hat{\kappa}) P(c_{i2} \mid f_j, N_2, \hat{\kappa}) \right].$$

In the reference analysis, empirical $\log P$ distributions were pooled within subject. A global pool was used as fallback for subjects with fewer than two successfully fitted replicate pairs. The operational lower-tail probability was

$$p_i = \widehat{F}_{\text{pool}}(\log P_i),$$

where $\widehat{F}_{\text{pool}}$ is the empirical cumulative distribution function of the corresponding reference pool. A state was classified as operationally observable when

$$p_i < \alpha,$$

with $\alpha = 0.05$ in the reference analysis. This annotation subsequently defines the operational endpoint classes used in the longitudinal TT/TF/FT/FF analyses. Re-aggregation at alternative values of $\alpha$ changes the empirical operational annotation but does not require refitting the Negative-Binomial latent model.

All complete replicate pairs were submitted to latent inference without upstream quality-based exclusion. Posterior width, entropy, replicate agreement, depth imbalance, ONE_POSITIVE/TWO_POSITIVE status, and posterior-predictive detectability were retained as diagnostics only and were not used to filter successfully fitted states. Numerical optimizer failure was treated differently because it does not yield a valid fitted latent model: unsuccessful pair fits were documented but were not admitted to the canonical aggregated state table. Downstream inclusion therefore depended on successful numerical inference, not on cohort-relative QC thresholds.

##### Code implementation

The analyses described in this section correspond to the following production scripts:

- `1-repertoire_characterization.py`: repertoire preprocessing, descriptive Pareto-tail fitting, and Pareto-versus-lower-truncated-log-normal robustness analysis.
- `2-multirepresentation_clonotype_state_inference.py`: technical-replicate pairing, construction of replicate-resolved observed measurements, pair-specific latent fitting, state-table aggregation, and operational observability.
- `noiseK_latent.py`: Negative-Binomial latent-frequency model, pair-specific empirical-Bayes estimation of $\gamma$ and $\kappa$, joint and single-replicate posteriors, posterior uncertainty summaries, and posterior-predictive detectability.

#### (ii) Assembly and characterization of the longitudinal transition dataset

##### Standardized multirepresentation trajectory assembly

Longitudinal trajectory construction was implemented in `4-multirepresentation_trajectory_assembly.py` (Step 4), in the general tradition of cross-sample clonotype tracking in repertoire-sequencing studies (Shugay et al., 2015; Pogorelyy et al., 2018; Minervina et al., 2020). Its canonical input was the combined Step-2 clonotype-state table. The production state key was

$$\text{subject} \times \text{aaSeqCDR3} \times \text{time},$$

with optional additional grouping variables for pseudo-longitudinal or null workflows. The production input was required to contain exactly one row per state key. Duplicate state keys were treated as an upstream data-contract violation and caused the assembly to stop; they were not averaged, summed, or otherwise collapsed.

Step 4 preserved three conceptually distinct state layers. First, the model-based latent consensus state was propagated directly from the paired-replicate posterior without refitting,

$$f^{\text{latent}} = f_{\text{latent,median}}, \qquad x^{\text{latent}} = x_{\text{latent,median}}.$$

When available, the 2.5th and 97.5th posterior quantiles, posterior log-frequency standard deviation, posterior entropy, fitted-model marginal log probability, and single-replicate latent posterior summaries were retained as additional model-based fields.

Second, replicate-resolved observed measurements were reconstructed directly from the measured counts and sequencing depths,

$$f_r^{\text{obs}} = \frac{c_r}{N_r},$$

and, for positive counts only,

$$x_r^{\text{obs}} = \log f_r^{\text{obs}}.$$

If $c_r = 0$, the corresponding observed log-frequency remained undefined; no pseudocount or finite abundance surrogate was introduced. Realized replicate positivity was stored separately for each technical replicate. An auxiliary positive-only representation was also carried forward for compatibility and sensitivity analyses: for TWO_POSITIVE states it was the geometric mean of the two observed relative frequencies, whereas for ONE_POSITIVE states it was the observed frequency of the single positive replicate. This auxiliary representation was not used as the primary forward or fluctuation measurement layer.

Third, observation annotations were retained separately from abundance representations. The continuous empirical `p_value`, posterior-predictive `p_detect_state` and `p_dropout_state`, and the reference operational-observability indicator were carried forward. The reference indicator was reconstructed as

$$\text{observable}_{\text{ref}} = \mathbf{1}(p_{\text{value}} < \alpha_{\text{ref}}),$$

with $\alpha_{\text{ref}} = 0.05$ in the reference analysis. Upstream and reconstructed reference observability were validated for consistency. The continuous endpoint `p_value` was retained so that alternative operational thresholds could later be evaluated without rebuilding the state table.

Step 4 was strictly an assembly layer. It did not refit the observation model, alter posterior estimates, estimate dynamics, or remove states according to abundance, posterior width, posterior entropy, replicate agreement, realized positivity class, posterior-predictive detectability, `p_value`, or operational observability. Instead, malformed identifiers, negative counts, non-positive sequencing depths, invalid latent states, out-of-range probabilities, or duplicate state keys generated explicit errors.

##### Generic finite-time transition assembly

Finite-time transitions were assembled with `5-longitudinal_transition_assembly.py` (Step 5). For each subject, and separately within any explicit extra grouping variables, all available time pairs $t_0 < t_1$ satisfying

$$1 \leq t_1 - t_0 \leq 5$$

were considered. Endpoint tables were inner-joined by `aaSeqCDR3`; consequently, a transition was generated when the same subject-specific clonotype was represented in the Step-4 state table at both endpoints. Step 5 did not synthesize unobserved both-zero states and did not remove endpoint pairs according to operational observation class.

For each transition,

$$\Delta t = t_1 - t_0,$$

and the latent consensus endpoints were recorded as $x_0^{\text{latent}}$ and $x_1^{\text{latent}}$. The latent displacement and midpoint were

$$\Delta x^{\text{latent}} = x_1^{\text{latent}} - x_0^{\text{latent}},$$

$$x_{\text{mid}}^{\text{latent}} = \frac{x_0^{\text{latent}} + x_1^{\text{latent}}}{2}.$$

The latent displacement was retained as a representation-level quantity and was not defined by the assembly step as the primary biological forward-drift estimand.

When endpoint posterior log-frequency standard deviations $\sigma_0$ and $\sigma_1$ were finite and strictly positive, Step 5 additionally constructed

$$w_0 = \frac{1}{\sigma_0^2 + 10^{-12}}, \qquad w_1 = \frac{1}{\sigma_1^2 + 10^{-12}},$$

and

$$x_\star^{\text{latent}} = \frac{w_0 x_0^{\text{latent}} + w_1 x_1^{\text{latent}}}{w_0 + w_1}.$$

If valid endpoint uncertainties were unavailable, $x_\star^{\text{latent}}$ was set to $x_{\text{mid}}^{\text{latent}}$; the source of the coordinate was recorded explicitly. The approximate uncertainty of the difference based only on endpoint marginal standard deviations was also retained as

$$\sigma_{\Delta x,\text{approx}} = \sqrt{\sigma_0^2 + \sigma_1^2}.$$

These quantities were model-based conditioning or sensitivity representations rather than estimand-specific filters.

**Replicate-resolved observed transition layer and estimand support**

Observed replicate-specific endpoint measurements were retained as counts, sequencing depths, relative frequencies, and log-frequencies at $t_0$ and $t_1$. For replicate $r$,

$$\Delta x_r^{\text{obs}} = x_r^{\text{obs}}(t_1) - x_r^{\text{obs}}(t_0).$$

Because $x_r^{\text{obs}}$ was defined only for positive counts, $\Delta x_r^{\text{obs}}$ was defined only when replicate $r$ was positive at both endpoints. Step 5 did not impute these values at an observation boundary.

The generic transition table included explicit support flags for the primary downstream estimands. For the AB orientation, replicate 1 supplies the initial conditioning measurement and replicate 2 supplies the displacement. Therefore,

$$\text{AB eligible} = I(c_{1,t_0} > 0)\, I(c_{2,t_0} > 0)\, I(c_{2,t_1} > 0).$$

For the reciprocal BA orientation,

$$\text{BA eligible} = I(c_{2,t_0} > 0)\, I(c_{1,t_0} > 0)\, I(c_{1,t_1} > 0).$$

The strict support for the cross-replicate fluctuation estimator was

$$\text{common4} = \prod_{r=1}^{2} I(c_{r,t_0} > 0) I(c_{r,t_1} > 0),$$

that is, both technical replicates were positive at both endpoints. On this support, both observed displacements are defined and later analyses can estimate the conditional cross-replicate covariance

$$\text{Cov}\left(\Delta x_1^{\text{obs}}, \Delta x_2^{\text{obs}} \mid x_{\text{mid}}^{\text{latent}}, \Delta t\right).$$

These flags encoded measurement eligibility only. They were not quality scores and were not used during Step 5 to remove transitions.

##### Operational endpoint annotation within the unrestricted transition universe

Step 5 retained the continuous empirical observation statistic at both endpoints, `p_value_t0` and `p_value_t1`, together with the reference operational-observability indicators. Reference endpoint labels were defined by

$$T: \; p_{\text{value}} < \alpha_{\text{ref}}, \qquad F: \; p_{\text{value}} \geq \alpha_{\text{ref}},$$

and combined into the four reference classes TT, TF, FT, and FF. These labels describe operational endpoint observability at the specified threshold and were not interpreted as biological persistence, expansion, contraction, extinction, or absence. No observation-class filtering was performed during transition assembly. Preserving the continuous endpoint `p_value` fields allowed downstream sensitivity analyses to reconstruct the operational domain at alternative $\alpha$ values on exactly the same transition universe.

##### Full-posterior endpoint uncertainty propagation

Step 5 supports an additional model-based uncertainty layer based on the complete discrete latent posterior caches generated upstream. In the production analysis in which this layer was requested, each endpoint posterior was sampled independently. For transition $t_0 \to t_1$ and $M = 1000$ Monte Carlo draws,

$$x_0^{(m)} \sim P_0(x), \qquad x_1^{(m)} \sim P_1(x), \qquad m = 1, \ldots, M,$$

and

$$\Delta x^{(m)} = x_1^{(m)} - x_0^{(m)}.$$

Sampling was performed from the endpoint discrete posterior probability vectors on their stored latent-frequency grids after conversion to log-frequency. A deterministic seed derived from the base seed, subject, temporal pair, and any explicit extra grouping variables was used to make the propagation reproducible.

For each successfully propagated transition, the endpoint posterior means and medians and the displacement mean, median, 2.5th percentile, 97.5th percentile, and standard deviation were stored. Directional posterior probabilities were estimated as

$$P(\Delta x > 0 \mid \text{data}) \approx \frac{1}{M} \sum_{m=1}^{M} I(\Delta x^{(m)} > 0),$$

and analogously for $P(\Delta x < 0 \mid \text{data})$. Endpoint posterior draws were independent conditional on their separately fitted time-point models. The propagated displacement distribution therefore reflects marginal endpoint uncertainty and does not estimate cross-time posterior covariance from a joint longitudinal model. Full-posterior propagation was an optional uncertainty representation; it was not required to construct the primary replicate-decoupled forward estimator or the primary cross-replicate fluctuation estimator.

###### Transition-universe and estimand-support characterization

The assembled transition table was characterized and audited with `6-transition_support_characterization.py` (Step 6). This step did not estimate longitudinal dynamics. Instead, it quantified the generic transition universe and tested whether the fields required by the downstream estimands were structurally available on their intended support.

Coverage was summarized overall and by biological subject, subject-specific temporal interval, nominal $t_0 \to t_1$ interval, and temporal lag $\Delta t$. The composition of the reference TT/TF/FT/FF classes and the fractions of transitions satisfying AB, BA, and `common4` eligibility were also reported. When additional pseudo/null grouping variables were present, these extended the analysis-unit keys but were not counted as additional biological subjects.

Because observed log-frequency is intentionally undefined for a zero-count replicate, completeness was evaluated conditionally on the support where a field is required rather than by demanding that every observed log-frequency field be non-null on every transition. For example, `dx_observed_rep1` was required to be finite where replicate 1 was positive at both endpoints; AB readiness required finite replicate-1 initial conditioning and replicate-2 displacement on `forward_ab_eligible`; BA used the reciprocal requirement; and `common4` readiness required both observed displacements and $x_{\text{mid}}^{\text{latent}}$ to be finite. This support-conditional audit distinguished legitimate observation-boundary missingness from true field incompleteness.

Step 6 also independently checked structural consistency, including

$$\Delta t = t_1 - t_0,$$

agreement between read-positivity flags and observed counts, agreement of AB/BA/`common4` flags with their definitions, and agreement of endpoint observability with $p_{\text{value}} < \alpha_{\text{ref}}$. Optional model-based fields such as $x_{\star}^{\text{latent}}$, endpoint posterior uncertainties, approximate latent-displacement uncertainty, posterior-predictive detectability, and full-posterior propagated displacement summaries were characterized for completeness but were not required for the primary observed estimands.

All Step-6 checks were descriptive production-readiness audits. They did not change transition membership, select TT as a quality class, impose abundance or posterior-quality thresholds, or estimate forward drift, fluctuation covariance, temporal scaling, or observation-threshold effects. The final Step-5 transition table therefore remained the common finite-time substrate for the subsequent estimand-specific analyses.

##### (iii) Pseudo-longitudinal technical calibration and definition of the primary estimators

###### Pseudo-longitudinal technical-reference ensemble and inferential unit

Technical calibration used the pseudo 1×12 dataset and was implemented in two production analyzers, motivated by the established need to quantify technical variability and standardize AIRR-seq measurements using replicate or control material (Barennes et al., 2021; Trück et al., 2021): `7-pseudo_forward_technical_null_compact.py` (Step 7) and `8-pseudo_fluctuation_conditioning_validation_pairbank.py` (Step 8). The source dataset contained one unchanged biological source represented by 12 independent technical TCR-repertoire measurements. These measurements were reassembled upstream into 2,000 randomized configurations, each organized as six pseudo-time points with two technical replicates per pseudo-time point.

The randomization unit and the biological unit were explicitly distinguished. `configuration_id` identifies a randomized reassembly of the same 12 technical measurements and was never treated as an additional biological subject. Consequently, no subject bootstrap was performed. Across-configuration medians and 2.5th–97.5th percentiles describe the distribution induced by pseudo-longitudinal randomization and were reported as randomization intervals rather than biological confidence intervals. Likewise, pseudo-lag $\Delta t$ represents distance within an arbitrary technical ordering and must not be interpreted as elapsed biological time.

Step 7 used the one-step compact transition cache ($\Delta t = 1$) for forward technical-reference calibration. Step 8 evaluated fluctuation behavior over pseudo-lags $\Delta t \in \{1, 2, 3, 4, 5\}$. Neither analyzer counted configurations as biological replicates.

##### Mathematical coupling and rationale for replicate decoupling

The forward calibration explicitly addressed the algebraic coupling created when the same noisy baseline measurement appears both on the horizontal axis and inside the displacement, a classical source of spurious baseline–change association in repeated-measurement analyses (Oldham, 1962; Chiolero et al., 2013). Consider an unchanged underlying log-abundance $x^*$ observed with measurement error,

$$y_{r,t} = x^* + \varepsilon_{r,t}.$$

For same-replicate conditioning, the displacement between two pseudo-time points is

$$\Delta y_r = y_{r,1} - y_{r,0} = \varepsilon_{r,1} - \varepsilon_{r,0}.$$

Hence,

$$\text{Cov}(y_{r,0}, \Delta y_r) = \text{Cov}(\varepsilon_{r,0}, \varepsilon_{r,1}) - \text{Var}(\varepsilon_{r,0}).$$

If baseline and endpoint errors are independent within a replicate, this reduces to

$$\text{Cov}(y_{r,0}, \Delta y_r) = -\,\text{Var}(\varepsilon_{r,0}) < 0.$$

Thus, a restoring-like or mean-reverting pattern can arise purely from measurement geometry. For reciprocal replicate decoupling, however, conditioning and displacement are taken from different technical replicates. Under replicate-independent technical errors,

$$\text{Cov}\left(\varepsilon_{A,0}, \varepsilon_{B,1} - \varepsilon_{B,0}\right) = 0.$$

These relations motivated the matched same-versus-cross calibration but were not imposed as fitting constraints. Residual deviations from the ideal null were treated empirically as the combined consequence of finite sampling, technical measurement, state representation, binning, and arbitrary pseudo-time assignment.

##### Step-7 primary observed replicate-decoupled forward benchmark

The production Step-7 compact cache retained the quantities required for the final forward estimator: $x_0^{\text{latent}}$, $x_{\text{mid}}^{\text{latent}}$, $x_\star^{\text{latent}}$, $\Delta x^{\text{latent}}$, replicate-specific observed initial log-frequencies, replicate-specific observed displacements, and the `forward_ab_eligible`, `forward_ba_eligible`, and `common4` support flags.

The primary forward benchmark used the observed replicate-resolved channel. For configuration $c$ and abundance bin $b$, the AB fold was

$$\bar{d}_{\text{AB},c}(b) = E\left[\Delta x_2^{\text{obs}} \mid x_{1,t_0}^{\text{obs}} \in B_b, \text{AB eligible}\right],$$

where AB eligibility requires replicate 1 to be positive at $t_0$ and replicate 2 to be positive at both endpoints. The reciprocal BA fold was

$$\bar{d}_{\text{BA},c}(b) = E\left[\Delta x_1^{\text{obs}} \mid x_{2,t_0}^{\text{obs}} \in B_b, \text{BA eligible}\right].$$

AB and BA were always analyzed separately. The combined curve was formed only after bin-specific summaries had been calculated,

$$\bar{d}_{\text{cross},c}(b) = \frac{1}{2}\bar{d}_{\text{AB},c}(b) + \frac{1}{2}\bar{d}_{\text{BA},c}(b).$$

The same exact 0.5/0.5 rule was applied to the fold-specific median-displacement descriptors and to $P(\Delta x > 0)$. Row counts were retained for support auditing but were never used as AB/BA estimator weights.

Global forward bin edges were themselves constructed with equal fold influence. Within every selected configuration, equal numbers of eligible AB and BA baseline values were sampled for edge estimation. With production defaults, the total target was 2,000 sampled baseline values per configuration, split equally between AB and BA. Twenty global quantile bins were then constructed over the 0.5th–99.5th percentile range. A fold-specific bin required at least 50 observations, and curve-level descriptors required at least six valid bins.

For each valid curve, Step 7 retained the mean of bin-specific mean displacements, mean absolute bin displacement, root-mean-square and maximum absolute bin displacement, mean absolute directional-probability bias,

$$\frac{1}{B}\sum_{b} \left| P(\Delta x > 0 \mid B_b) - 0.5 \right|,$$

and the linear slope, intercept, and zero crossing of mean displacement against bin center. AB–BA concordance was summarized across matched valid bins using Pearson correlation and absolute fold differences.

**Matched same-versus-cross coupling control**

To isolate the effect of replicate decoupling from differences in measurement support, a matched comparison was performed on `common4`, for which both technical replicates were positive at both endpoints. Four curves were constructed on the same global forward bins:

$$\text{same A}: \quad x^{\text{obs}}_{1,t_0} \longrightarrow \Delta x^{\text{obs}}_{1},$$

$$\text{cross AB}_{\text{matched}}: \quad x^{\text{obs}}_{1,t_0} \longrightarrow \Delta x^{\text{obs}}_{2},$$

$$\text{same B}: \quad x^{\text{obs}}_{2,t_0} \longrightarrow \Delta x^{\text{obs}}_{2},$$

and

$$\text{cross BA}_{\text{matched}}: \quad x^{\text{obs}}_{2,t_0} \longrightarrow \Delta x^{\text{obs}}_{1}.$$

The two same-measure curves and the two matched cross-replicate curves were each combined with exact equal fold weight,

$$\bar{d}_{\text{same}}(b) = \frac{1}{2}\bar{d}_{\text{same A}}(b) + \frac{1}{2}\bar{d}_{\text{same B}}(b),$$

$$\bar{d}_{\text{cross,matched}}(b) = \frac{1}{2}\bar{d}_{\text{cross AB,matched}}(b) + \frac{1}{2}\bar{d}_{\text{cross BA,matched}}(b).$$

Step 7 additionally recorded the fraction of the same-measure mean absolute bin displacement retained after cross-replicate decoupling and the corresponding change in abundance-dependent slope. These quantities were calibration descriptors rather than hypothesis-test statistics.

##### Secondary latent conditioning-geometry diagnostic

The compact Step-7 cache also supported a secondary comparison of the latent conditioning coordinates

$$x_0^{\text{latent}}, \qquad x_{\text{mid}}^{\text{latent}}, \qquad x_\star^{\text{latent}}$$

against the same latent displacement $\Delta x^{\text{latent}}$. Only rows finite for the displacement and all three candidate coordinates entered this complete-case comparison. A common absolute support was defined from the overlapping central ranges of the three coordinates, using the 1st–99th percentile limits, and divided into 30 equal-width bins with production defaults. Equal numbers of rows were sampled from each randomization configuration for construction of the common support so that no configuration dominated the bin definition.

This block was explicitly descriptive. It did not select a universal conditioning coordinate and, in the current compact workflow, it was not restricted to TT or any other operational observation class because `obs_class_reference` is not part of the compact input contract. The latent conditioning diagnostic therefore should not be described as the historical TT-only benchmark.

##### Step-8 replicate-consistent fluctuation calibration

Fluctuation calibration used `8-pseudo_fluctuation_conditioning_validation_pairbank.py` and was restricted to `common4`, ensuring that both observed replicate displacements were finite at both endpoints. For a given configuration, pseudo-lag, and abundance bin, let

$$A = \Delta x_1^{\text{obs}}, \qquad B = \Delta x_2^{\text{obs}}.$$

The primary replicate-consistent fluctuation component was the signed cross-replicate covariance

$$C_{\text{cross}} = \text{Cov}(A, B).$$

The mean same-replicate displacement variance was

$$V_{\text{same}} = \frac{1}{2}\left[\text{Var}(A) + \text{Var}(B)\right],$$

and the replicate-specific excess was

$$V_{\text{rep}} = V_{\text{same}} - C_{\text{cross}}.$$

When $V_{\text{same}} > 0$, the auxiliary shared-fluctuation ratio was

$$F_{\text{shared}} = \frac{C_{\text{cross}}}{V_{\text{same}}}.$$

The covariance was kept signed throughout and was never truncated at zero. Sample variances and covariance used $\text{ddof} = 1$ in the production configuration. As a numerical identity audit, the implementation verified

$$\text{Cov}(A, B) = \text{Var}\left(\frac{A+B}{2}\right) - \frac{1}{4}\,\text{Var}(A - B),$$

with the residual retained as an implementation diagnostic.

The production primary conditioning coordinate was $x_{\text{mid}}^{\text{latent}}$. The abundance grid was derived once from equal-size samples of this coordinate from each selected configuration, using the central 1st–99th percentile range and 20 equal-width bins by default. The exact same grid was then reused for the conditioning sensitivities $x_{\star}^{\text{latent}}$ and `xmid_observed`, preventing changes in bin boundaries from being conflated with changes in conditioning representation.

For the primary and sensitivity coordinates, binned covariance calculations used finite paired observed displacements and finite values of the relevant conditioning variable. Each conditioning representation therefore used its own complete cases. Direct comparisons between conditioning representations were performed on matched valid bins. Production defaults required at least 10 transitions per binned cell and at least six valid abundance bins for a curve descriptor. Pooled configuration-level fluctuation summaries required at least 20 transitions.

Step 8 summarized $C_{\text{cross}}$, $V_{\text{same}}$, and $V_{\text{rep}}$ both by pseudo-lag and by abundance bin. To reduce dependence on unequal numbers of nominal intervals contributing to different pseudo-lags, equal-interval summaries were also calculated by first estimating the fluctuation quantity within each nominal interval and then averaging across valid intervals. Linear slopes across pseudo-lags 1–5 were retained as technical-randomization diagnostics only; pseudo-lag has no biological time interpretation.

##### Randomization summaries and final estimator policy

Step-7 and Step-8 quantities were first computed separately within each of the 2,000 randomized configurations. Ensemble summaries were then obtained from the across-configuration distribution, using the configuration median together with the 2.5th and 97.5th randomization percentiles. No subject bootstrap and no nested pseudo-time permutation were applied by these analyzers.

The technical calibration defined the production estimands without collapsing the distinct roles of the available representations. The primary forward estimator uses observed replicate-decoupled AB/BA measurements and exact equal fold weight after binning. The primary fluctuation estimator is the signed cross-replicate covariance

$$\text{Cov}\left(\Delta x_1^{\text{obs}}, \Delta x_2^{\text{obs}} \mid x_{\text{mid}}^{\text{latent}}, \Delta t, \text{common4}\right).$$

$x_{\star}^{\text{latent}}$ and the observed midpoint are retained as conditioning sensitivities, while $x_0^{\text{latent}}$, $x_{\text{mid}}^{\text{latent}}$, and $x_{\star}^{\text{latent}}$ against $\Delta x^{\text{latent}}$ form a separate latent-geometry diagnostic. No composite performance score, automatic dispersion-based selection rule, or universal representation ranking was used.

#### (iv) Replicate-decoupled forward dynamics and comparison with the pseudo-longitudinal technical reference

##### Observed replicate-decoupled forward estimator

The genuine longitudinal forward analysis was implemented in `9-observed_replicate_decoupled_forward_drift.py` using the generic Step-5 transition table. The primary lag was $\Delta t = 1$ week. The analysis used the replicate-resolved observed log-frequency fields and the estimand-specific eligibility flags generated during transition assembly; replicate-specific latent posterior summaries were not required.

For the AB orientation, the conditioning variable was the observed baseline log-frequency in replicate 1 and the outcome was the observed displacement measured entirely in replicate 2. For the reciprocal BA orientation, the roles of the replicates were exchanged:

$$\text{AB}: \qquad x_A(t_0) \longrightarrow \Delta x_B = x_B(t_1) - x_B(t_0),$$

$$\text{BA}: \qquad x_B(t_0) \longrightarrow \Delta x_A = x_A(t_1) - x_A(t_0).$$

AB eligibility required replicate 1 to be positive at $t_0$ and replicate 2 to be positive at both endpoints; BA used the reciprocal requirement. These conditions correspond to `forward_ab_eligible` and `forward_ba_eligible`. No primary restriction by operational TT/TF/FT/FF class was imposed.

##### Abundance binning and fold combination

One common observed-abundance grid was used for AB, BA and the matched same-versus-cross controls. With the production settings, 20 quantile bins were defined over the 0.005–0.995 range of an exact equal-mass mixture of the two primary conditioning distributions. Each AB conditioning value received weight $0.5/n_{\mathrm{AB}}$ and each BA conditioning value weight $0.5/n_{\mathrm{BA}}$ when the bin edges were constructed. Thus, bin geometry and combined estimates were invariant to unequal AB/BA row counts.

Within each fold and abundance bin, the analysis retained the number of transitions and contributing subjects, mean and median displacement, the probability of positive displacement, and the displacement standard deviation. A bin was considered valid when it contained at least 50 eligible transitions. For bins valid in both folds, the combined forward quantities were calculated with exact equal fold weight:

$$\overline{\Delta x}_{\text{cross}}(b) = \frac{1}{2}\left[\overline{\Delta x}_{\mathrm{AB}}(b) + \overline{\Delta x}_{\mathrm{BA}}(b)\right],$$

$$P_{\text{cross}}(\Delta x > 0 \mid b) = \frac{1}{2}\left[P_{\mathrm{AB}}(\Delta x > 0 \mid b) + P_{\mathrm{BA}}(\Delta x > 0 \mid b)\right].$$

AB and BA were therefore never concatenated and treated as independent biological observations. Fold-specific support was retained separately because the reciprocal folds can contain overlapping transitions.

##### Subject-level uncertainty and curve descriptors

Uncertainty in the longitudinal forward profile was quantified by resampling biological subjects with replacement (2,000 bootstrap replicates; seed 123), preserving the subject as the resampling unit for the clustered longitudinal data (Deen and de Rooij, 2020). The same resampled subject set was used simultaneously for AB, BA and the matched controls. Each fold was reconstructed separately within every bootstrap sample and reciprocal folds were then combined with the same 0.5/0.5 rule used for the point estimate. Percentile bootstrap intervals were obtained from the resulting subject-cluster distributions.

For descriptive summarization, an unweighted linear relation was fitted across the fixed set of valid abundance-bin centers:

$$\overline{\Delta x}(b) = \alpha + \beta x_b, \qquad x_{\text{zero}} = -\frac{\alpha}{\beta}.$$

The slope, intercept and zero-crossing were treated as compact descriptors of the nonparametric binned profile rather than as parameters of a prespecified generative drift model. Bootstrap descriptors were evaluated on the same valid-bin domain as the corresponding point estimate. Subject-specific and leave-one-subject-out curves were generated as robustness diagnostics.

##### Matched same-measure control

The direct effect of reusing the same technical measurement for both conditioning and displacement was assessed on `common4`, where both replicates were positive at both transition endpoints. On this matched support, same-replicate and cross-replicate constructions were evaluated on exactly the same transitions:

$$\text{same A}: \quad x_A(t_0) \to \Delta x_A, \qquad \text{cross AB}: \quad x_A(t_0) \to \Delta x_B,$$

$$\text{same B}: \quad x_B(t_0) \to \Delta x_B, \qquad \text{cross BA}: \quad x_B(t_0) \to \Delta x_A.$$

The same and cross controls were combined separately with equal A/B or AB/BA fold weight. This analysis was used as a technical coupling diagnostic and did not replace the broader estimand-specific AB/BA support of the primary forward analysis.

**Reference to the pseudo-longitudinal technical reference**

Comparison with the pseudo-longitudinal technical reference was implemented in `10-longitudinal_vs_pseudo_forward_null.py`. Step 10 required the final Step-7 and Step-9 outputs to implement the same observed replicate-decoupled AB/BA estimand and the same exact equal-fold combination policy; it did not introduce a TT/TF/FT/FF filter.

Only the methodological features required to interpret the comparison are summarized here. The absolute-abundance analysis retained the native longitudinal and pseudo bin geometries and compared them only on their shared observed-log-frequency support, without extrapolation; pseudo native bins missing in an individual configuration were not imputed. A complementary relative-abundance analysis assigned transition-level mid-rank percentiles separately within biological subject × fold for the longitudinal cohort and within randomization configuration × fold for the pseudo ensemble:

$$p = \frac{\text{average rank} - 0.5}{n}.$$

AB and BA were percentile-ranked and binned separately before equal-fold combination. Longitudinal intervals in this comparison remained biological subject-cluster bootstrap intervals, whereas pseudo intervals reflected the distribution across randomized pseudo configurations and were therefore randomization intervals rather than biological confidence intervals. Curve-level empirical comparisons used the pseudo-configuration distribution with a finite-sample +1 correction. The detailed comparison descriptors and their interpretation are presented with the corresponding Results.

#### (v) Replicate-resolved fluctuation dynamics, technical-reference comparison and temporal scaling

**Cross-replicate fluctuation estimand**

The genuine longitudinal fluctuation analysis was implemented in `11-cross_replicate_fluctuation_dynamics.py` from the generic Step-5 transition table. The primary support was `common4`, defined as transitions for which both observed technical replicates were positive at both endpoints. Operational TT/TF/FT/FF classes were not used as a primary filter. The conditioning coordinate was the latent midpoint, whereas the fluctuation outcomes were the two independently observed replicate-resolved displacements.

$$\Delta x_A = x_A(t_1) - x_A(t_0), \qquad \Delta x_B = x_B(t_1) - x_B(t_0).$$

Within a lag × abundance cell containing $n$ transitions, the primary estimator was the sample cross-replicate covariance

$$V_{\text{cross}} = \frac{1}{n-1} \sum_i \left(\Delta x_{A,i} - \overline{\Delta x}_A\right) \left(\Delta x_{B,i} - \overline{\Delta x}_B\right).$$

The within-replicate variance comparator and replicate-specific excess were

$$V_{\text{same}} = \frac{1}{2} \left[\text{Var}(\Delta x_A) + \text{Var}(\Delta x_B)\right],$$

$$V_{\text{excess}} = V_{\text{same}} - V_{\text{cross}}.$$

The complementary shared-fluctuation ratio, defined as cross-replicate covariance divided by mean within-replicate variance, was reported when the denominator was positive. All covariance and variance estimates used `ddof=1`. Importantly, cross-replicate covariance remained signed; negative estimates were not clipped to zero. The implementation additionally verified the covariance identity

$$\text{Cov}(A, B) = \text{Var}\left(\frac{A+B}{2}\right) - \frac{1}{4}\,\text{Var}(A - B)$$

numerically in pooled, subject-level and interval-level cells as an internal construction audit.

##### Abundance grid, lag-resolved profiles and uncertainty

A single abundance grid was reused across $\Delta t = 1$–5 weeks so that lag-specific profiles were evaluated on a common latent-midpoint geometry. With the production settings, 20 equal-width bins were defined between the 0.01 and 0.99 quantiles of the latent midpoint among finite `common4` transitions pooled across the selected lags. A pooled lag × abundance cell was considered supported when it contained at least 50 transitions.

For each supported cell, the analysis retained cross-replicate covariance, mean within-replicate variance, replicate-specific excess, shared-fluctuation ratio, replicate-specific mean displacements, transition count, number of contributing subjects and number of contributing subject-specific physical intervals. The primary cohort curve was transition-weighted: sufficient statistics were summed across subjects before reconstructing the covariance rather than averaging subject-level covariance estimates.

Biological uncertainty was quantified with 2,000 subject-cluster bootstrap resamples (seed 123), preserving within-subject dependence during resampling (Deen and de Rooij, 2020). Subjects were sampled with replacement, and the pooled sufficient statistics were reconstructed from the selected subject multiplicities before recalculating the covariance decomposition. A bootstrap draw contributed to a lag × abundance-cell interval only when its reconstructed transition count satisfied the same minimum-$n$ criterion as the point estimate. Subject × lag × bin and subject × physical-interval × bin estimates were also retained for downstream temporal and interval-position analyses.

##### Reference to the pseudo-longitudinal technical reference

The technical-reference comparison was implemented in `12-longitudinal_vs_pseudo_cross_replicate_fluctuations.py` using exactly the same primary estimand, support and conditioning coordinate in the genuine and pseudo-longitudinal datasets. Because covariance magnitude is abundance dependent, the comparison was performed on matched absolute latent-midpoint support separately at each lag; no percentile normalization was used. Stable pseudo native bins were identified from configuration-level Step-8 support, and pseudo configurations were required to provide directly valid values over the matched bin set. Missing pseudo bins were not interpolated or imputed. Only the genuine longitudinal curve was locally interpolated between adjacent valid Step-11 bins when required to evaluate it at the matched Step-8 native bin centers.

The lag-specific scalar comparison used the equal-bin mean of cross-replicate covariance across the retained matched abundance bins. Pseudo configurations represented randomization realizations rather than biological subjects, whereas longitudinal uncertainty remained based on biological subject resampling. This technical-reference comparison was not used as a preprocessing correction: Step-12 covariance was not subtracted from Step-11 values before temporal-scaling analysis.

##### Fixed-core temporal scaling across 1–5 weeks

Temporal scaling was implemented in `13-temporal_fluctuation_scaling.py` using the subject × lag × abundance-bin output of Step 11. Step 12 was not a computational input. To prevent lag-specific changes in abundance or subject composition from generating an apparent temporal trend, both the abundance domain and the biological cohort were fixed before fitting temporal models.

Let $N_{\text{available}}(\Delta t)$ denote the number of subjects structurally available at lag $\Delta t$. A Step-11 abundance bin qualified for the temporal core only when, at every selected lag, it contained at least

$$\lceil 0.90\, N_{\text{available}}(\Delta t) \rceil$$

subjects with a finite subject-level cross covariance and at least two transitions in that subject × lag × bin cell, the mathematical minimum for covariance with `ddof=1`. The largest contiguous run of qualifying bins defined the primary abundance core. The primary complete-case cohort then retained only subjects with valid values in every selected core bin at every lag.

##### Primary temporal estimator and sensitivity weighting

For complete-case subject $s$ at lag $\Delta t$, the primary subject-level core summary gave every selected abundance bin equal weight:

$$M_s(\Delta t) = \frac{1}{|\mathcal{B}|}\sum_{b\in\mathcal{B}} V_{\text{cross},s,b}(\Delta t),$$

where $\mathcal{B}$ is the fixed abundance core. The cohort temporal profile then gave every complete-case subject equal weight:

$$M(\Delta t) = \frac{1}{|\mathcal{S}|}\sum_{s\in\mathcal{S}} M_s(\Delta t).$$

This equal-bin/equal-subject construction was the primary estimator (`equal_bin_equal_subject`). A precision-oriented sensitivity (`df_weighted_within_subject_equal_subject`) weighted bins within each subject × lag by their covariance degrees of freedom, $n-1$, and then averaged subjects equally. The sensitivity weighting did not replace the fixed-geometry primary estimator.

Cross-replicate covariance was the primary temporal metric. The mean within-replicate variance and replicate-specific excess were analysed as technical comparators. The shared-fluctuation ratio was not used as a default temporal-scaling metric because it can become unstable when its variance denominator is small.

##### Signed temporal slope, bootstrap inference and finite-lag models

Temporal dependence was summarized by fitting the five lag-specific cohort values with the signed linear descriptor. Analyses of fluctuation magnitude as a function of temporal separation have previously been used to characterize T-cell clone-size dynamics, although with different estimands and time scales (Gaimann et al., 2020; Bensouda Koraichi et al., 2023).

$$M(\Delta t) = K + D(\Delta t - 1), \qquad \Delta t = 1, \dots, 5,$$

where $K$ denotes the fitted one-week level and $D$ is the signed slope per week. The sign of $D$, rather than a positive-only accumulation model, was the primary temporal descriptor. Biological uncertainty was estimated by a joint subject-cluster bootstrap with 2,000 resamples: one subject draw was generated per bootstrap replicate and propagated unchanged across all lags, metrics and estimators. The resulting distribution provided the bootstrap median, 2.5th–97.5th percentile interval, and the fractions of slopes above and below zero.

A temporal slope was also calculated separately for each complete-case subject, and exact sign-flip tests of the mean subject-specific slope were reported as a conservative complementary inference. In addition, three simple finite-lag descriptions were compared on each cohort curve:

$$\text{constant:} \qquad M(\Delta t) = K,$$

$$\text{signed linear:} \qquad M(\Delta t) = K + D(\Delta t - 1),$$

$$\text{non-negative incremental-linear:} \qquad M(\Delta t) = K + D(\Delta t - 1), \qquad D \geq 0.$$

Model comparison used AICc with the residual-variance parameter included, using the small-sample correction to Akaike's information criterion (Hurvich and Tsai, 1989). Because only five temporal lags were available, AICc preference was treated as a descriptive finite-lag comparison and was kept separate from signed-slope evidence. Within the selected core, bin-specific temporal slopes were additionally estimated as an abundance-resolved secondary analysis using the same complete-case subjects and joint bootstrap draws. No specific stochastic mechanism was inferred from the five sampled lag values.

#### References


Barennes P, Quiniou V, Shugay M, et al. Benchmarking of T cell receptor repertoire profiling methods reveals large systematic biases. *Nature Biotechnology*. 2021;39:236–245. doi:10.1038/s41587-020-0656-3.

Bensouda Koraichi M, Ferri S, Walczak AM, Mora T. Inferring the T cell repertoire dynamics of healthy individuals. *Proceedings of the National Academy of Sciences of the USA*. 2023;120(4):e2207516120. doi:10.1073/pnas.2207516120.

Chiolero A, Paradis G, Rich B, Hanley JA. Assessing the relationship between the baseline value of a continuous variable and subsequent change over time. *Frontiers in Public Health*. 2013;1:29. doi:10.3389/fpubh.2013.00029.

Clauset A, Shalizi CR, Newman MEJ. Power-law distributions in empirical data. *SIAM Review*. 2009;51(4):661–703. doi:10.1137/070710111.

Deen M, de Rooij M. ClusterBootstrap: An R package for the analysis of hierarchical data using generalized linear models with the cluster bootstrap. *Behavior Research Methods*. 2020;52(2):572–590. doi:10.3758/s13428-019-01252-y.

Desponds J, Mora T, Walczak AM. Fluctuating fitness shapes the clone-size distribution of immune repertoires. *Proceedings of the National Academy of Sciences of the USA*. 2016;113(2):274–279. doi:10.1073/pnas.1512977112.

Gaimann MU, Nguyen M, Desponds J, Mayer A. Early life imprints the hierarchy of T cell clone sizes. *eLife*. 2020;9:e61639. doi:10.7554/eLife.61639.

Gelman A, Meng XL, Stern H. Posterior predictive assessment of model fitness via realized discrepancies. *Statistica Sinica*. 1996;6:733–760.

Hurvich CM, Tsai CL. Regression and time series model selection in small samples. *Biometrika*. 1989;76(2):297–307. doi:10.1093/biomet/76.2.297.

Koch H, Starenki D, Cooper SJ, Myers RM, Li Q. powerTCR: A model-based approach to comparative analysis of the clone size distribution of the T cell receptor repertoire. *PLoS Computational Biology*. 2018;14(11):e1006571. doi:10.1371/journal.pcbi.1006571.

Minervina AA, Pogorelyy MV, Komech EA, et al. Primary and secondary anti-viral response captured by the dynamics and phenotype of individual T cell clones. *eLife*. 2020;9:e53704. doi:10.7554/eLife.53704.

Oldham PD. A note on the analysis of repeated measurements of the same subjects. *Journal of Chronic Diseases*. 1962;15(10):969–977. doi:10.1016/0021-9681(62)90116-9.

Pogorelyy MV, Minervina AA, Puelma Touzel M, et al. Precise tracking of vaccine-responding T cell clones reveals convergent and personalized response in identical twins. *Proceedings of the National Academy of Sciences of the USA*. 2018;115(50):12704–12709. doi:10.1073/pnas.1809642115.

Puelma Touzel M, Walczak AM, Mora T. Inferring the immune response from repertoire sequencing. *PLoS Computational Biology*. 2020;16(4):e1007873. doi:10.1371/journal.pcbi.1007873.

Shugay M, Bagaev DV, Turchaninova MA, et al. VDJtools: Unifying post-analysis of T cell receptor repertoires. *PLoS Computational Biology*. 2015;11(11):e1004503. doi:10.1371/journal.pcbi.1004503.

Trück J, Eugster A, Barennes P, et al. Biological controls for standardization and interpretation of adaptive immune receptor repertoire profiling. *eLife*. 2021;10:e66274. doi:10.7554/eLife.66274.
